# A compact vocabulary of paratope-epitope interactions enables predictability of antibody-antigen binding

**DOI:** 10.1101/759498

**Authors:** Rahmad Akbar, Philippe A. Robert, Milena Pavlović, Jeliazko R. Jeliazkov, Igor Snapkov, Andrei Slabodkin, Cédric R. Weber, Lonneke Scheffer, Enkelejda Miho, Ingrid Hobæk Haff, Dag Trygve Tryslew Haug, Fridtjof Lund-Johansen, Yana Safonova, Geir K. Sandve, Victor Greiff

**Author notes:** Correspondence: Victor Greiff. Equal contribution (Philippe A. Robert, Milena Pavlović).

## Abstract

Antibody-antigen binding relies on the specific interaction of amino acids at the paratope-epitope interface. The predictability of antibody-antigen binding is a prerequisite for de novo antibody and (neo-)epitope design. A fundamental premise for the predictability of antibody-antigen binding is the existence of paratope-epitope interaction motifs that are universally shared among antibody-antigen structures. In the largest set of non-redundant antibody-antigen structures, we identified structural interaction motifs, which together compose a commonly shared structure-based vocabulary of paratope-epitope interactions. We show that this vocabulary enables the machine learnability of antibody-antigen binding on the paratope-epitope level using generative machine learning. The vocabulary (i) is compact, less than 10^4^ motifs, (ii) distinct from non-immune protein-protein interactions, and (iii) mediates specific oligo- and polyreactive interactions between paratope-epitope pairs. Our work successfully leveraged combined structure- and sequence-based learning showing that machine-learning-driven predictive paratope and epitope engineering is feasible.

## Introduction

Antibody-antigen binding is mediated by the interaction of amino acids at the paratope-epitope interface of an antibody-antigen complex. A long-standing question in the fields of immunology and structural biology is whether paratope-epitope interaction is predictable. The predictability of paratope-epitope binding is a prerequisite for predicting antibody specificity and *in silico* antibody and vaccine design. So far, however, it remains unclear whether antibody-antigen binding is predictable (*1–3*).

The current paradigm is that antibody-antigen interaction is too high-dimensional of an entity to permit prediction. This long-standing belief most likely has its roots in the works of Karl Landsteiner and colleagues showing that antibodies can recognize a nearly infinite number of antigens (*4*). The complexity paradigm was reinforced over the subsequent decades by sequencing and crystallography studies suggesting that not only the interaction space is infinite, but also the sequence and structural space (with minor constraints on the antibody side) (*2, 5, 6*).

Antibody binding to the epitope is mainly formed by the three hypervariable regions termed complementarity-determining regions (CDRs) situated in both antibody heavy and light chains (*7*– *9*). The hypervariability of the CDR3 is key to the immunological specificity of antibodies (*10*) and is generated by somatic recombination of the variable (V), diversity (D, only for the heavy chain), and joining (J) genes of the B-cell genomic locus (*11*). Combinatorial diversity from rearranged germline gene segments, somatic hypermutation, and antigen-driven selection steps enable antibodies to interact specifically with virtually any given antigen (*11–13*).

The most reliable method for identifying paratope-epitope pairs is by solving the 3D structure of antigen-antibody complexes and determining which amino acids in the two partners make contact with each other (*14*). Over the last decades, the increasing amount of antibody-antigen structures has enabled quantitative insights into the physicochemical features of antibody-antigen interaction (*3, 15–25*). For example, it has been observed repeatedly that paratopes localize mostly, but not exclusively, to CDRs (*26*), and that certain amino acids are preferentially enriched or depleted in the antibody binding regions (*3, 27–30*). For epitopes, several analyses have shown that their amino-acid composition is essentially indistinguishable from that of other surface-exposed non-epitope residues if the corresponding antibody is not taken into account (*31–34*).

Recently, computational and machine learning approaches for the sequence-based and structural prediction of paratopes (*35–37*), epitopes (*38*) or paratope-epitope (antibody-antigen) interaction (*34, 36, 39–41*) are accumulating (for a more complete list of references see here: (*1, 2, 42–45*)). While the accuracy for the prediction of paratopes seems generally higher than that for epitopes, to our knowledge, no study has yet conclusively shown that antibody-antigen interaction is a priori predictable and if so, based on what theoretical and biological grounds (*1, 46*).

Recent reports have provided preliminary evidence for the potential predictability of antibody-antigen interaction: (i) The antibody repertoire field has now established that antibody sequence diversity underlies predictable rules (*5, 6, 47*). (ii) The presence of transferable “specificity units” between distinct antibody molecules was recently suggested by showing that tightly binding functional antibodies may be conceived by designing and improving seemingly unrelated paratopes (*48*). Previous efforts towards predicting paratope-epitope interaction have been stifled by both a one-sided investigation of either exclusively the paratope or the epitope and the failure to break down the problem of antibody-antigen interaction into its *fundamental units.* The fundamental units of antibody-antigen interaction are the sequence regions on the antibody and the antigen that compose the paratope-epitope interface. The 3D complex structure of an epitope typically emerges from different sub-peptides of the protein, folded in the same place. Therefore, the binding units go beyond a single linear peptide, hindering the power of sequence-based prediction tools. We conjectured that the comparison of those interaction units across antibody-antigen complexes may lead to the discovery of a general vocabulary of antibody-antigen interaction. If a general compact (finite) vocabulary for antibody-antigen interaction existed that unambiguously formed paratope-epitope pairs, then paratope-epitope interaction would be a priori predictable.

To investigate the veracity of our conjecture, we devised a domain-specific sequence language that encodes the units of paratope and epitope interaction as *structural interaction motifs* (henceforth termed interaction motif or simply motif). These structural interaction motifs (i) conserve the link between paratope and epitope and (ii) allow for the comparison of paratope-epitope structural interactions across antibody-antigen complexes.

Here, we screened the largest available set of curated non-redundant antibody-antigen structures for binding patterns and identified a vocabulary of structural interaction motifs (≈400 paratope and ≈1600 epitope motifs) that govern antibody-antigen interaction. This interaction motif vocabulary has the following characteristics: (i) it is compact (finite and restricted) and immunity-specific. Indeed, we found that the motif vocabulary – although tens of orders of magnitude less diverse than antibody receptors and antigen sequences – covers already 50% of all possible paratope and 15% of all possible epitope motifs. We excluded the possibility that the motif vocabulary discovered is a trivial subset of the protein-protein binding space. (ii) The paratope motif vocabulary shows specific and mutually exclusive (unambiguous) epitope reactivity. Specifically, a small number of paratope motifs were found to bind to as many as 50% of all observed epitope motifs. These polyreactive paratopes bound to distinct epitope spaces. (iii) The motif vocabulary is seemingly universal. Interaction motifs were shared across entirely unrelated antibody-antigen complexes (different antibody germline genes, different antigen classes). The existence of a vocabulary of antibody-antigen binding that nearly unambiguously links paratope to epitope implies that antibody-antigen binding is a priori predictable. To quantify the learnability of antibody-antigen binding, we leveraged statistical modeling (shallow learning) and deep learning. We showed that paratope-epitope interaction pairing can be learned with multi-class prediction accuracies ranging from 58–75% (baseline accuracy of randomly permuted data: 19–47%) at the motif level. Superimposing structural interaction motifs onto sequence-based prediction of paratope-epitope interaction improved predictability by up to 7 percentage points (with respect to sequence-only) suggesting that motifs, with respect to the primary sequence, occupy an orthogonal function in the mechanism of antibody-antigen binding.

## Methods

### A dataset of non-redundant and diverse 3D antibody-antigen complexes

A dataset of antibody-antigen complexes in the format of Protein Data Bank (PDB) (Fig. 1A) was obtained from the Antibody Database (AbDb) (*49, 50*). AbDb routinely crawls PDB to find existing antibody-antigen structures and preprocesses them by (i) identifying the antibody (VH-VL, variable [V] heavy [H] and light [L] chain domains) and the corresponding ligand, (ii) annotating the antibody variable region (Fv) by consulting the Summary of Antibody Crystal Structure (SACS) database (*51*) and (iii) applying a standardized numbering scheme for the antibody sequences. To obtain non-redundant structures, AbDb performs pairwise comparisons across the structures (both heavy and light chains). Structures comprising different amino acid residues in the same position are considered non-redundant. As of 15 June 2019, AbDb stored 866 complexes. From this initial dataset, we removed atoms labeled with PDB record type HETATM (non-protein atoms serving as co-factors) and structures with a resolution larger than 4.0Å (*44*). The final curated dataset comprises 825 antibody-antigen (protein antigen only) complexes with a median resolution of 2.5Å (Fig. 1A). To gain statistical power, we analyzed mouse and human antibody complexes as one entity. Mouse and human antibody-antigen complexes represent 90% of the AbDb (Suppl Fig. 11) and we found only minor differences in paratope motif spaces used by mouse and humans (Suppl Fig. 4C, 14). In support, it was recently reported that neither CDR/FR length nor the distribution of interface residues in human and murine antibodies differs substantially (*29, 52, 53*).

**Figure 1.**
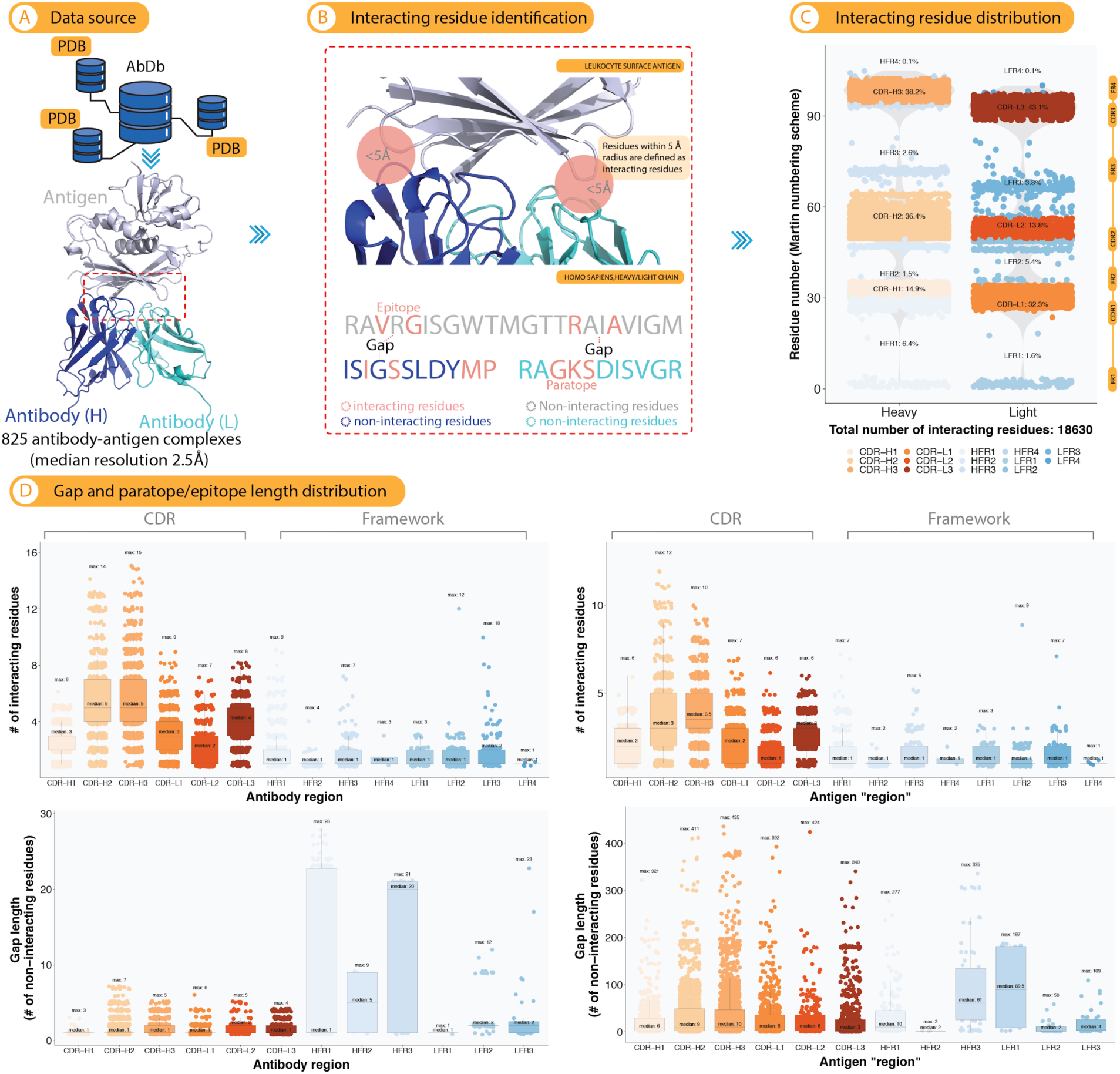
Characterization of interacting residues at the paratope-epitope interface. (A) We characterized antibody-antigen interaction using publicly available 3D structures deposited in the Antibody Database (AbDb) (see Methods), a total of 825 antibody-antigen complexes (dark/light blue: antibody V_H_/V_L_, grey: antigen). Antigens are distributed across a wide range of antigen classes (Suppl Fig. 11) and antibodies are ≈90% of human and mouse origin (Suppl Fig. 11). **(B)** Interacting residues were defined as the set of residues at the antibody-antigen interface within a radius of <5Å from each other. Here we considered only the distance between heavy atoms (non-hydrogen atoms) of residues at the interface. These residues are termed *paratope* in antibody and *epitope* in antigen. More specifically, a paratope was defined as the set of interacting amino acid residues within a particular FR or CDR region of an antibody. An epitope is defined as the set of antigen amino acid residues that interact with a paratope. Epitopes are annotated according to the FR or CDR regions of the corresponding paratopes. Gaps are defined as the non-interacting residues that lie in between interacting residues. **(C)** The interacting residues mapped predominantly to the CDRs and less so to the framework regions (sequence annotation follows the Martin numbering scheme, see Methods). **(D)** The number of interacting residues ranged between 2–5 (CDR, median) and 1–2 (FR, median) in paratopes and 1–3.5 (CDR, median) and 1 (FR, median) in epitopes. Gap lengths (number of non-interacting residues) ranged between 1–2 (CDR, median) and 1–20 (FR, median) in paratopes and 3–10 (CDR, median) and 2–89.5 (FR, median) in epitopes.

Annotations for 113 broadly neutralizing antibodies were obtained from the database bNAber (*54*). 70 of these antibodies were represented as 24 non-redundant complexes in AbDb and were included herein (Suppl Fig. S5). The remaining 38 (without antigens) and 5 missing structures were excluded.

### Selection of antibody sequence numbering scheme

AbDb provides datasets with three numbering schemes: Kabat (*55*), Chothia (*56*), and Martin (*57*). These numbering schemes partition the antibody heavy and light chains into framework (FR1, FR2, FR3, FR4) and CDR (CDR1, CDR2, and CDR3) regions. In the Kabat scheme, gaps found within the alignment are based on the variability of the aligned sequences. As more three-dimensional (3D) structural information became available, Chothia and Lesk created a numbering scheme that takes spatial alignment into consideration. In particular, they corrected the positioning of the first CDR in both heavy and light chains. Abhinandan and Martin further refined the Chothia numbering scheme by making corrections, not only in the CDRs but also in the FRs. Here, we used the Martin numbering scheme to annotate the FRs and CDRs of antibodies as it was previously determined to be suitable for structural and antibody engineering (*58*). It is also the most recent of the presently available numbering schemes. Supplementary Table S1 summarizes the position of FR and CDR regions and the position of insertions according to the Martin numbering scheme. In the Martin numbering scheme, the CDR-H3 region excludes the V-gene germline part of the antibody gene (typically identified by the amino acid triplet CAR), as well as parts of the J-gene germline part (typically identified by “W”) as shown in Supplementary Table S2.

**Table 1.** Summary of machine learning use cases (see also Methods text and. **Figure 5**).

| Incorporation of amino acid position | Direction* | Data type |
| --- | --- | --- |
| No | Paratope to epitope | Motif |
| Yes | Paratope to epitope | Motif |
| No | Epitope to paratope | Motif |
| Yes | Epitope to paratope | Motif |
| No | Paratope to epitope | Sequence |
| No | Epitope to paratope | Sequence |
| No | Paratope to epitope | Aggregate (sequence + motif) |
| No | Epitope to paratope | Aggregate (sequence + motif) |

\* Use cases for PPI are organized in a similar fashion.

### Identification of interacting residues in antibody-antigen complexes

To identify interactions between amino acid residues in antibody-antigen complexes, a distance cutoff was set. Distance cutoffs between 4–6Å are routinely used when examining interactions between proteins or protein-ligand pairs as most noncovalent atomic interactions are short-range (e.g., hydrogen bonds and Van der Waals interactions range from 3–4Å (*43, 59*)). For instance, a recent study on contact-based protein structure networks by Viloria and colleagues found that a distance cutoff of <5Å (heavy atoms) is most sensitive to changes in residue interactions and variables such as force fields (*60*). Therefore, we defined interacting paratope-epitope residues by a distance cutoff of <5Å between heavy atoms. In other words, amino acid residues are considered to be interacting if they have heavy atoms with a distance <5Å from each other (Fig. 1B). For completeness, we evaluated the variation of the total number of interacting residues, their distribution across FR and CDR regions as well as the overlap of interaction motifs (for explanation see below) for the three commonly used distance cutoffs <4Å, <5Å and <6Å in Suppl Fig. S3 and Fig. S7. We confirmed that the overall trends in per-region residue distribution, overlap between paratope and epitope interaction motifs (Suppl Fig. S3), and (dis)continuity (Suppl Fig. S7) hold across the three distance cutoffs tested indicating that our definition of interacting residues is appropriate and robust.

### Definition of paratope, epitope, and paratope-epitope structural interaction motifs

(i) A paratope is defined as the set of interacting amino acid residues within a particular FR or CDR region of an antibody (e.g., residues colored in salmon in Fig. 1B). (ii) An epitope is defined as the set of antigen amino acid residues that interact with a paratope. Epitopes are annotated according to the FR or CDR regions of the corresponding paratopes. (iii) The length of a paratope or epitope is defined as the number of amino acid residues constituting the paratope or epitope (see paratope/epitope length in Fig. 1D). (iv) A gap is defined as the number of non-interacting residues separating two paratope or epitope residues (Fig. 1B, 1D, and 2A). (v) A paratope or epitope structural interaction motif is composed of interacting paratope and epitope amino acid residues as well as non-interacting ones (gap). Interacting residues are encoded with the letter `X` and non-interacting residues are encoded with an integer quantifying gap size (number of non-interacting residues, Fig. 2A). For example, the string `X1X` encodes a paratope or epitope interaction motif of two interacting amino acid residues (`X,X`) separated by one non-interacting residue (1).

**Figure 2.**
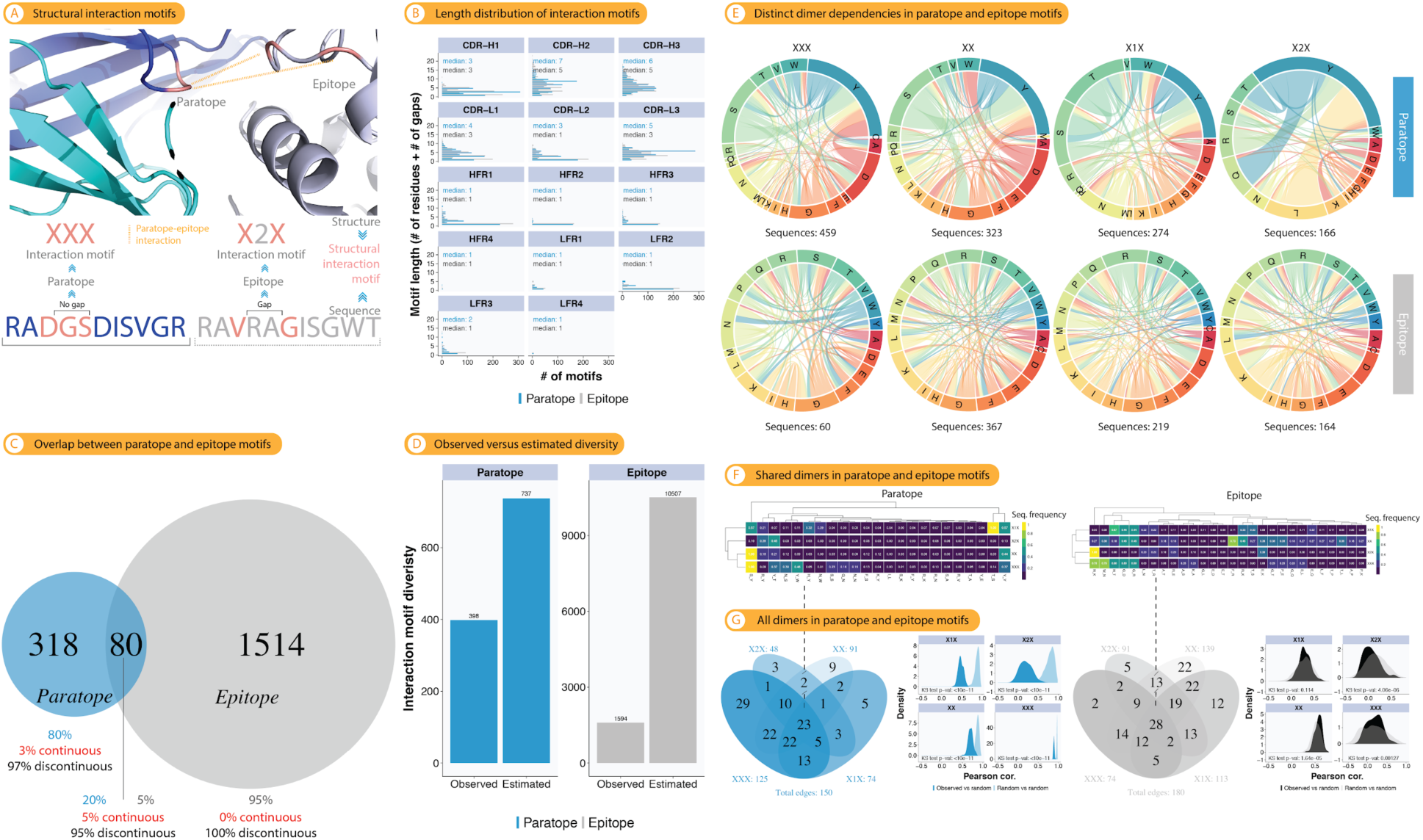
Structural interaction motifs represent a compact vocabulary for the composition of the paratope-epitope interface. (A) We devised a structural interaction motif notation that accounts simultaneously for gaps and residues in both paratopes and epitopes. A paratope or epitope structural interaction sequence motif is composed of interacting paratope and epitope amino acid residues as well as non-interacting ones (gap). Interacting residues are encoded with the letter `X` and non-interacting residues are encoded with an integer quantifying a gap size. For example, the string `X1X` encodes a paratope or epitope motif of two interacting amino acid residues (`X`,`X`) separated by one non-interacting residue (1). **(B)** Length distribution of motifs by paratope, epitope, and FR/CDR region. **(C)** Absolute and relative overlap of paratope and epitope motifs (Venn diagram). **(D)** Estimation of the potential (observed + unobserved) motif diversity using the Chao1 estimator (see *Methods*). **(E)** For each of the four most highly shared (across structures) interaction motifs (Suppl Fig. 4), the sequential dependency signature was determined. Briefly, for the ensemble of paratope/epitope sequences mapping to a given interaction motif, the 2-mer decomposition of each paratope/epitope sequence was determined by a bidirectional sliding window (2-mer frequency distribution). For each motif, these sequential dependencies were visualized as Chord diagrams where the 20 amino acids form the segments in a track (outermost ring) and the links are the frequency with which a 2-mer occurred (sequential dependency). **(F)** Hierarchical clustering of sequential dependencies (2-mers) that were shared between all four paratope or epitope motifs. **(G)** Venn diagrams: Overlap of sequential dependencies (two-mers) shared across paratope or epitope motifs. Density plots: We tested whether the 2-mer distribution (sequential dependencies) observed in (F) for each of the 4 motifs could be due to random effects. To this end, we sampled 100 times 2-mers from the number of 2-mers possible (G) according to the number of sequences mapping to each motif (E) and calculated the correlation either among all randomly drawn 2-mer distributions (grey: epitope, light blue: paratope) or between observed and randomly drawn one (black: epitope, dark blue: paratope). The significance in the difference between the distributions was tested using the Kolmogorov-Smirnov test.

### Definition of interaction motif angle

The angle of an interaction motif was computed by defining two vectors spanning the midpoint of a motif and its start and end positions (see inset in Suppl Fig. S8 for illustration), in a similar fashion to `AngleBetweenHelices`, a `Pymol` module for calculating the angle between helices (*61*). Larger angles would indicate that the structure of the interaction motif is more extended whereas small angles indicate that the interaction motif would tend to form a loop. Protein 3D structures were rendered and visualized in `Pymol 2.1.0` (*61*).

### Diversity analysis of interaction motifs

To estimate the potential (observed + unobserved) paratope or epitope sequence diversity, we used the Chao1 estimator (*62–64*), a non-parametric estimator of the lower bound of species richness [number of unique sequence motifs], as implemented in the R package `Fossil 0.3.7` (`Chao1`).

### Paratope-epitope amino acid contact map

Paratope(P)-epitope(E) amino acid contact maps were obtained by computing the log odds ratio, of the observed occurrence of each amino acid pair over the corresponding expected frequency as described in (*31*); where is the paratope amino acid, is the epitope amino acid, and is the region (FR/CDR in antibody-antigen complexes). Analogously, protein-protein amino acid contact maps were computed for inter- and intradomain in non-immune protein-protein complexes (PPI).

### Construction of bipartite paratope-epitope and PPI reactivity networks at motif and sequence level

A paratope-epitope motif interaction network (*reactivity network*) was constructed by connecting each paratope motif to its corresponding epitope motif (undirected edge). The degree distribution, the distribution of the number of connections (edges) to a node (degree), of the resulting interaction network was tested to fit a power-law distribution by calculating a goodness-of-fit value with bootstrapping using the `poweRlaw R 0.70.2` package (*65*) as described by Clauset and colleagues (*66*). Here, a network whose degree distribution fits a power-law distribution (exponent between 2 and 3) is defined as scale-free (*67*). Networks and the corresponding visualizations were constructed using the network analysis and visualization suite `Cytoscape 3.7.1` (*68*). Reactivity networks for sequence and aggregate encoding, see machine learning use cases (encoding) below, as well as PPI reactivity networks, were constructed as above described and are shown in Suppl Fig. S12 and Fig. S19, respectively.

### Analysis of sequential dependencies in interaction motifs

To quantify the sequential dependencies in paratope and epitope interaction motifs, we determined for each multi-residue motif, the 2-mer decomposition of each paratope/epitope sequence (bidirectional sliding window) of the ensemble of paratope/epitope sequences mapping to the respective motif (non-interacting residues were not taken into account). For each motif, these sequential dependencies were visualized as Chord diagrams where the 20 amino acids form the segments in a track (the outermost ring) and the links indicate the frequency with which a 2-mer sequential dependency occurred (sequential dependency). Chord diagrams were constructed using `Circlize 0.4.8` (*69*). Hierarchical clustering of the motifs’ sequential dependencies was performed using the `R` package `pheatmap 1.0.12` (*70*), distances between motifs were quantified by Euclidean distance or correlation and agglomeration was carried out using the complete-linkage method.

### Dataset of protein-protein interaction and definition of protein-protein interaction motifs

A dataset of protein-protein interactions (PPI) was sourced from 3did, a catalog of three-dimensional structure domain-based interactions (*71*). The database (i) collects high-resolution 3D-structures from PDB (`version 2019_1`) (*50*) and (ii) annotates the structures according to the protein domain definitions provided by Pfam (`version 32.0`, Table S3 summarizes the top 10 protein domains in the latest version 3did) (*72*). Interactions between domains originating from different chains were annotated as *interdomain* whereas interactions originating from the same chain as *intradomain*. Structures with Pfam domain description (i) immunoglobulin and (ii) Ig-like were excluded (as they overlap with structures from AbDb). As of 2 July 2019, 3did comprised a total of 18,599,078 contact residue pairs (100,888 protein structures), which is three orders of magnitude larger than the number of antibody-antigen contact residues (18,630 residue pairs, Fig. 1C). Protein-protein interaction motifs were constructed for each domain pair analogously to paratope-epitope interaction motifs (see the previous section). Motifs with gap length larger than seven were excluded from the analysis (to match the largest gap size found in paratopes, Fig. 1) as well as complexes larger than 300 residues long. The final non-immune PPI dataset comprises 9621 interdomain and 1043 intradomain complexes for a total of 299,141 contact residues, Suppl Fig. S6.

### Quantification of somatic hypermutation on antibody amino acid sequences

To quantify somatically hypermutated (SHM) amino acid residues in the dataset, we annotated the sequences with the corresponding species (here shown only human and mouse) and aligned the sequences against germline immunoglobulin V, D, and J genes sourced from IMGT.

The IMGT database (*73*) includes 570 (578), 34 (39), and 32 (26) human (mouse) germline immunoglobulin V, D, and J genes, respectively. We translated the nucleotide sequences of V and J genes according to their ORFs (open reading frame). As D genes can be truncated during the recombination process, we used amino acid sequences corresponding to all three ORFs (excluding non-productive translations). To compute alignments, we used the following scoring scheme: match reward = 2, mismatch penalty = –1, gap opening penalty = –5, and gap extension penalty = –2. For each sequence, we selected germline V, D (if the sequence corresponds to the heavy chain), and J genes with the highest alignment scores. SHMs were defined as differences in the alignment between the antibody sequence and the selected germline genes. Exonucleolytic removals during V(D)J recombination lead to deterioration of the alignment quality at the end (start) of V (J) genes. To reduce their impact on SHM quantification, we discarded SHMs corresponding to three amino acid residues at the end (start) positions of V (J) genes in the alignment as it was shown previously that three amino acids (up to 9 nt) correspond to the average lengths of exonucleolytic removals in V and J genes (*74*). To reduce the impact of exonucleolytic removals in D genes, we considered only SHMs emerging between the first and the last matches in the alignments. Suppl Fig. S10 shows inferred SHMs localize around CDR1s and CDR2s and thus partially correlate with the paratopes positions centered in all three CDRs. Suppl Fig. S10 shows only few SHMs in the CDR3s. We note that this may be a reflection of the limitation of our SHM quantification approach and not necessarily a biological feature of the immunoglobulin sequences here studied.

### Ramachandran plot analysis

Ramachandran angles (Phi-Psi pairs) were extracted from PDB files using the package `PDB` in `Biopython 1.74` (*75*). The package `pdb-tools 2.0.0` was used to preprocess PDB files and extract the chains/regions of interest (*76*). We examined six different groups: (i) residues in the CDR regions of the heavy or light chains of antibody structures (CDR); (ii) residues in the framework regions of heavy and light chains of antibody structures (FR); (iii) residues binding to the antigen (paratope), from the FR and CDR regions i.e. only the ‘`X`’ in the motifs (AbDb interacting residues); (iv) binding residues from the PPI dataset in inter- and intra-chain interactions (PPI interacting residues); (v) residues that belong to a motif (including gaps) in AbDb antibody structures, for instance `X-X--X` leads to 6 angles (ABDB motifs), and (vi) residues that belong to a motif in intra- or inter-chain interactions in the PPI dataset (PPI motifs). Finally, following Hollingsworth and colleagues (*77*), we classified the Phi-Psi pairs into groups of secondary structure types (also known as Ramachandran regions).

### Machine-learning based prediction of paratope-epitope pairing

To quantify the extent to which paratope-epitope is learnable with the available dataset, we leveraged both deep and shallow learning approaches using several encodings of the input.

#### Use cases (encoding)

Four levels of encoding in both directions, namely paratope to epitope and epitope to paratope, were used. (i) Structural motif level: a paratope structural motif `XXX` interacting with an epitope motif `X2X` yields an input-output pair `XXX–X2X`. (ii) Position-augmented structural motif level: a paratope structural motif `XXX` interacting with an epitope motif `X2X` yields an input-output pair `X X X` –`X 2 X`, the positions index each character in the sequence consecutively. (iii) Sequence level: a paratope sequence NMA interacting with an epitope sequence RA yields an input-output pair NMA–RA. (iv) Finally, an aggregate representation which simultaneously takes into account amino acid information and motif by replacing the abstraction character ‘`X`’ with the corresponding residue: a paratope-epitope interaction defined by the paratope sequence GR and motif `X1X` together with the epitope sequence LLW and motif `XX1X` yields an input-output pair G-R–LL-W. The antibody-antigen (PPI) datasets comprise a total of 5,327 (25, 921) input-output pairs. Table 1 summarizes the setup for motif, sequence, and aggregate learning.

#### Deep learning

We leveraged a model based on Neural Machine Translation (*78*) to learn an epitope from (to) a paratope at motif, sequence and aggregate levels. Specifically, pairs of input-output sequences were translated via a combination of two components: encoder and decoder with gated recurrent units (GRU, see Fig. 5 bottom panel and the workflow graphic below). During the decoding phase via an attention layer, a context vector is derived to capture relevant input-side information necessary for the prediction of an output. Utilizing the context vector, the decoder part of our deep model generates each paratope or epitope motif/sequence character by character (generative model). For the translation task, we abstracted the gaps within a motif by replacing them with dashes, for example, all motifs of the form `X X` (where is any integer) were encoded simply as `X-X`. The dataset was split into 80% training and 20% test set. The numerical representation of the input pairs was learned by vector embedding. Pairwise parameter combination: (i) embedding dimension (1, 2^1^, 2^2^,…, 2^10^) and (ii) number of units (hidden dimension) (1, 2^1^, 2^2^,…, 2^10^) was used to parameterize the models. Here, the embedding dimension is the length of the vector representing the input whereas the number of units is the number of cells in the GRU otherwise known as the length of the hidden dimension. The training procedure was carried out for 20 epochs with Adaptive Moment Estimation (Adam) optimizer (*79*) and was replicated ten times. Each replicate comprises 121 models for a total of 1,210 models (121×10, see workflow). The model from the last epoch of each replicate was used to generate predictions on the test dataset.

#### Shallow learning

The shallow model takes into account the conditional probability of the output with respect to the input and a prior corresponding to the output with the highest marginal probability (the most frequent class).

#### Evaluation

Discrepancy (error) between predictions and the true motifs (sequences) was determined by the normalized Levenshtein distance, between the predicted motifs (sequences) and true motifs (sequences). Baseline prediction accuracies were calculated based on label-shuffled data where antibody and antigen-binding partners were randomly shuffled. To ensure robustness when evaluating the deep models, instead of showing the error obtained from the “best model” in each replicate, we showed the mean of median error across all replicates and pairwise parameter combinations. Ratios of training and test datasets, as well as error computation for the shallow model were identical to the above-described computation for deep models except for input motifs that were not present in the training dataset where the error was set to 1 (maximum error).

Deep learning models were constructed in `TensorFlow 1.13.1` (*80*) with `Keras 2.2.4-tf` (*81*) in `Python 3.6.4` (*82*), while the statistical (shallow) model was constructed using `pandas 0.25.1` (*83*). Computations for deep models were performed on the high-performance computing cluster Fram (Norwegian e-infrastructure for Research and Education sigma2.no/fram).

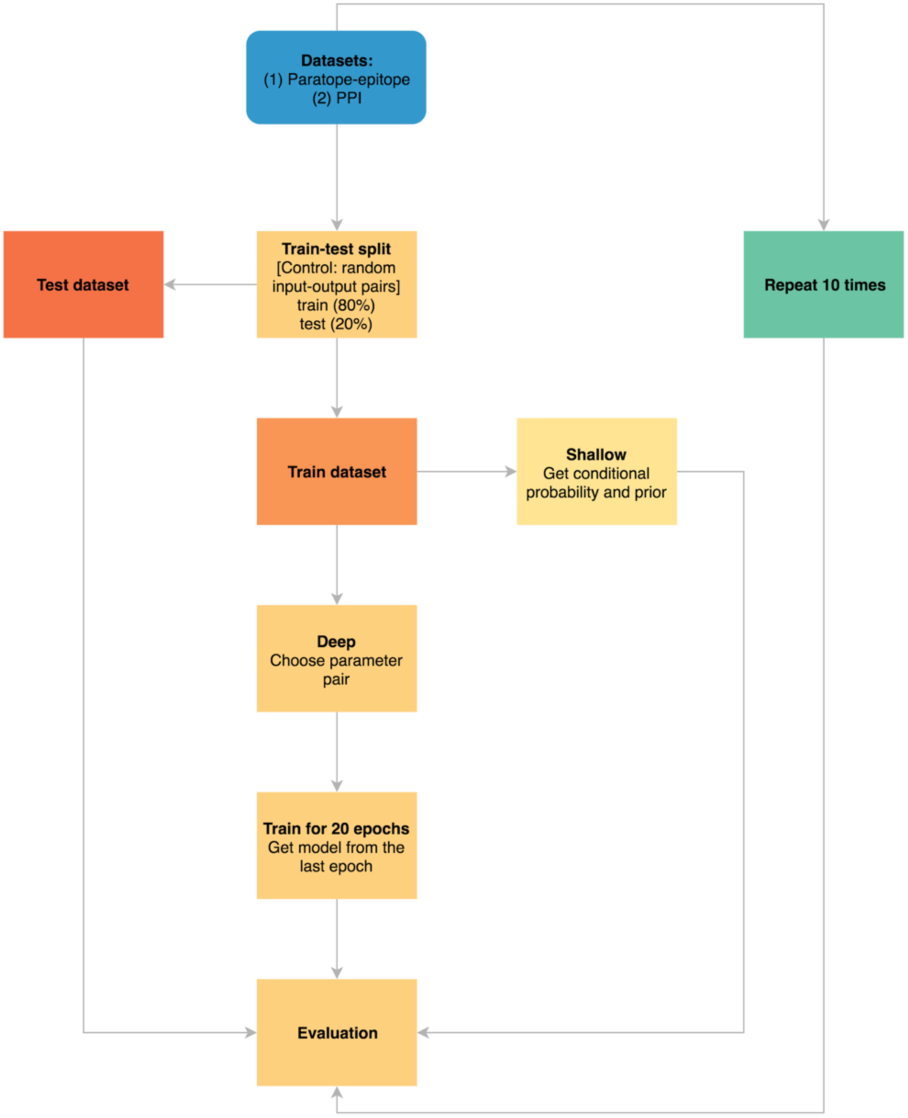
**Workflow for the machine learning prediction of paratope-epitope and PPI at interaction motif, sequence and aggregate level.** At the start of the workflow, a dataset comprising 5327 pairs of paratope-epitope motifs (or sequences or aggregate pairs) was split into a train (80%) and test (20%) datasets (PPI dataset comprises 25,921 pairs). For a deep model, a pair of parameter (embedding dimension and GRU unit) was selected from a set of 121 pairs. Each pair defines a model and each model was trained for 20 epochs on the training dataset. Once trained, the model was evaluated on the dataset by calculating (i) the correlation between the length of the predictions the true outputs (for deep model only) as well as (ii) normalized Levenshtein distance (see above) between the predictions and the true outputs. The procedure was replicated ten times. To accelerate the training process, parallelization was carried out in two stages: (i) jobs in the replication step (“repeat 10 times”) were sent to a set of nodes via jobarray in `SLURM` (*80*) and (ii) the 121 models were scattered across 121 CPUs via `mpi4py` (*81*). Baseline prediction accuracies were calculated based on label-shuffled data where paratope and epitope-binding partners were randomly shuffled (random pairing of paratope-epitope pairs). Shallow models follow identical train-test split and evaluation schemes (see Methods above).

### Graphics

All non-network graphics were generated using the statistical programming environment `R 3.5.2` (*84*) with the grammar of graphics R package `ggplot2 3.1.0` (*85*), the R package `Venn`Diagram 6`.20` (*86*), and the ggplot2 theme `themeakbar 0.1.2` (*87*). Figures were organized and schematics were designed using `Adobe Illustrator CC 2019`.

### Dataset and code availability and results reproducibility

(Preprocessed) datasets, code, and results figures are available at: https://github.com/GreiffLab/manuscript_ab_epitope_interaction.

## Results

### A diverse structural dataset for antibody-antigen binding

To gain a representative picture of antibody-antigen 3D interaction, we retrieved a dataset of all known 825 non-redundant antibody-antigen complexes (protein antigen only) archived in the Antibody Database (AbDb) (*49, 50*) (Fig. 1A). Antibody sequences, originating from this dataset, mapped to a diverse set of 64 (mouse) and 104 (human) V genes (Suppl Fig. S1A) and antigen sequences belonged to a diverse set of antigen classes from both humans and mice (Suppl Fig. 11). Both antibody (Ab) and antigen (Ag) sequences showed a high median sequence distance (>4) among either antibody framework (FR) and complementarity determining regions (CDR) (Suppl Fig. S1B) or antigen classes (Suppl Fig. S1C, median = 199). Thus, the dataset is diverse and not overly biased to one type of antibody or antigen class or to sequences of high similarity.

### The majority of paratope interacting residues are located in the antibody complementarity determining regions

We identified the set of interacting residues at the interface of antibody-antigen structures by using a heavy-atom distance cutoff of <5Å (*88*) (see Methods and see Suppl Fig. S3 and Fig. S7 for an examination of the robustness of the distance cutoff). Antibody-antigen amino acid pairs within this distance were designated as interacting residues (Fig. 1A–B). Together, the sets of antibody and antigen interacting residues form paratope-epitope pairs. In accord with previous reports (*20, 89*), paratope residues mapped overwhelmingly to the complementarity-determining regions (CDR) 1–3 (V_H,CDR1–3_: 89.5% and V_L,CDR1–3_: 89.2%) and, consequently, were only rarely found in the framework regions (FR) 1–4 (V_H,FR1–4_: 10.5% and V_L,FR1–4_: 10.8 %) (Fig. 1C). While the percentage of interacting residues found in the CDR3 was consistent between heavy and light chains, only half as many interacting residues mapped to CDR-H1 (14.8%) as compared to CDR-L1 (32.5%), whereas the reverse was true for CDR-H2 (36.1%) and CDR-L2 (13.9%). We verified that the number of interacting residues per FR/CDR is not a function of the underlying FR/CDR length (Suppl Fig. S9F). Since we used the Martin numbering scheme for CDR and FR annotation (see Methods), which mostly excludes germline gene residues from the CDR3, the above numbers demonstrate that germline-gene residues surrounding the CDR3 (FR3, FR4) contribute relatively little to antibody-interaction and that CDR3 paratope-epitope interaction is essentially non-germline gene residue driven (*57, 58*). Finally, we found that the position of paratope interacting residues correlates significantly (p<0.05) with the sites of (inferred) somatic hypermutation hotspots (SHM), Spearman (Pearson) correlation: 0.31–0.52 (0.44–0.58), Suppl Fig. S10) suggesting that interacting residues investigated herein have been subjected to antigen-driven selection.

### Paratopes are enriched in aromatic and polar residues whereas epitopes are enriched in charged residues

The residues tyrosine (V_H_: 18.2% and V_L_: 20.0%), and serine (V_H_: 11.2% and V_L_: 15.7%) were the two most frequent interacting residues found in paratopes. For epitopes, lysine (V_H_: 8.9% and V_L_: 9.6%), and arginine (V_H_: 8.7% and V_L_: 11.2%) were found to be the most abundant residues at the paratope-epitope interface (Suppl Fig. S2A–B). The enrichment of aromatic residues (namely tyrosine) in paratope sequences and the abundance of polar and charged residues in epitopes (e.g., lysine and arginine) are in accord with published literature (*23, 27*) validating further the robustness of our definition of interacting residues. While overall, the amino acid usage was more uniform in epitopes, both in paratope and epitope methionine and cysteine were heavily selected against. In gap (non-interacting) residues of paratope and epitope, cysteine was, however, more common (Suppl Fig. S2).

In terms of amino acid usage, the correlation between heavy and light chain paratope sites was very high (>0.82) as was the Pearson/Spearman correlation between the epitopes that were contacted by heavy and light chains (r_Spearman/Pearson_: >0.87, Suppl Fig. S6E). Paratope versus epitope residue usage correlation was far lower (r_Spearman/Pearson_:0.13–0.49). We also investigated the correlation between paratope and epitope residue usage with that of non-immune protein-protein interaction (PPI) residues and found that PPI and paratope were not correlated (r_Spearman/Pearson_: 0.06–0.29) whereas PPI and epitope were moderately correlated (r_Spearman/Pearson_: 0.57–0.71, Suppl Fig. 6D,E).

Finally, we investigated to what extent amino acid contact pairs of paratope and epitopes differed across CDR/FR regions and to what extent these amino-acid binding preferences overlap with those of non-immune protein-protein interaction (Suppl Fig. S15). We found substantial cross-type (type: charged, polar, aromatic, hydrophobic/nonpolar) interactions both at a local (CDR/FR region wise) and a global level (composite; Suppl Fig. S15A), while in contrast, PPI amino acid interaction preferences were much clearer where amino acids predominantly interact within-type rather than across types.

### Structural interaction motifs enable a unified comparison of paratope-epitope interfaces of unrelated antibody-antigen complexes

The fundamental units of antibody-antigen interaction are the sequence regions on the antibody and the antigen comprising the interacting and non-interaction residues at the binding interface termed paratope and epitope respectively. Consequently, paratopes and epitopes may manifest in two ways: (i) as a continuous stretch of interacting residues or (ii) discontinuously, separated by one or more non-interacting residues (gaps) due to protein folding (*8, 89*) (Fig. 1B).

The distributions of the number of interacting residues (termed paratope lengths) and the number of non-interacting residues (gap lengths) were analyzed within each region (FR or CDR) in (Fig. 1D). Paratope lengths ranged between 1 and 15 (median: 1–5) in the CDRs and 1–12 (median: 1–2) residues in the FR regions whereas the length of gaps ranged between 1 and 7 (median: 1–2) residues in CDRs and 1–28 (median: 1–20) residues in the FR regions (Fig. 1D). Epitopes can span up to 12 residues long with gaps up to 435 residues long (Fig. 1D). Therefore, the paratope-epitope interface can not be described exclusively by continuous stretches of amino acids.

As of now, it remains challenging to compare paratope-epitope interfaces across unrelated antibody-antigen complexes. To resolve this challenge, we devised a structural interaction motif notation that accounts simultaneously for gaps and residues in both paratopes and epitopes, to describe a structural pattern of binding. A paratope or epitope structural interaction motif is composed of interacting paratope and epitope amino acid residues as well as non-interacting ones (gap). Specifically, we encoded any interacting residue as capital `X` and any gaps as integers. Here, the integer quantifies the number of non-interacting amino acid residues (Fig. 2A). The combination of amino acid and gap encodings is termed *structural interaction motif* (henceforth interaction motif or simply, motif). Therefore, motifs describe the spatial conformation of the binding and can be used in addition to residue information to characterize antibody-antigen binding. For instance, the paratope DGS (three consecutive interacting residues) is denoted as an interaction motif and the epitope VRAG (two interacting residues, V and G, separated by two non-interacting residues, R and A) is denoted as an motif. Both motifs (and) have the same length: three. Our motif notation for antibody-antigen interaction places the paratope-epitope interface into a unified coordinate system that preserves the link between paratope and epitope and enables computational traceability of both continuous and discontinuous (structural) antibody-antigen interaction across antibody-antigen complexes. We will show that structural interaction motifs, although only defined on structural bases, inherently contain biological information on the underlying paratope and epitope amino acid usage, diversity and sequential dependency (see next Section and Figures 2E–G, Suppl Figs. 13, 14).

Paratope and epitope motif lengths varied across FR and CDR regions but remained below a length of 10 (median length: 1–7, Fig. 2B). Motifs were generally shorter in FRs (median length: 1–2) compared to CDRs (median length: 3–7). Interestingly, motifs of CDR-H2 (median length: 7) were longer than those in the CDR-H3 (median length: 6). Paratope and epitope motif lengths showed consistently positive correlation in CDRs (r_Pearson,CDR-H1–3 (L1–3)_ = 0.73–0.77 (0.5–0.84), r_Spearman, CDR-H1–3 (L1–3)_ = 0.7–0.74 (0.48–86), Suppl Fig. 9E). Furthermore, motifs were substantially shared across CDR and FR in both paratope and epitope (although in epitopes to a lesser extent than in paratopes) and were thus not exclusive to a specific CDR or FR (Suppl Fig. S4A, B lower triangle). On average, three to four motifs were found per antibody heavy or light chain (Suppl Fig. S9A).

With our motif vocabulary, we can now query key parameters of antibody-antigen recognition: (i) motif sequence diversity, (ii) structural diversity (motif angle and (dis)continuity), (iii) co-occurrence across complexes and (iv) predictability and learnability of paratope-epitope interaction. We note that we found no difference between interaction motifs found in HIV broadly neutralizing antibodies (bNAb), widely regarded as hallmarks of unusual antibody-binding behavior, and non-bNAb (Suppl Fig. S5A). Furthermore, the combined set of paratope and epitope motifs were generally distinct from those found in non-immune protein-protein interaction (PPI). Specifically, (1) PPI motifs were generally a few characters larger than paratope and epitope motifs (median length 1–7 in paratopes (CDR_1–3,L/H_) vs 7 in PPI (inter- and intradomain), Fig. 2B, Suppl Fig. S6B), and (2) while 57% of paratope motifs were found in PPI motifs, only 12% of epitope motifs were shared with PPI motifs (Suppl Fig. S6A).

### The diversity of paratope and epitope interaction motifs is restricted (compact)

We asked to what extent interaction motifs of paratopes and epitopes overlap. Out of 1594 and 398 unique paratope and epitope motifs, only 80 motifs overlapped (Fig. 2C). Only 5% of the shared motifs were continuous and 95% were discontinuous (Fig. 2C).

Next, we asked how much of the potential (observed + unobserved) paratope and epitope motif diversity is covered by our dataset of 825 antibody-antigen structures? To answer this question, we estimated the potential motif diversity using the Chao1 estimator (Chao, 1984) – a method often used in ecology for estimating population sizes (Fig. 2D, see *Methods* for details). For paratopes, the set of unique motifs in our dataset covered about 50% (398) of the potential diversity of all paratopes (estimated total: 737), whereas the set of unique epitope motifs in our dataset covered 15% (1, 594) of the total epitope diversity (10,507, Fig. 2D). The estimated potential size of the paratope motif space is one order of magnitude smaller than the theoretical size (≈10^5^, see Suppl Text for analytical derivation). Of interest, the size of the potential epitope motif space is similar to that of the PPI motif space (SupplFig. S6H).

To summarize, the estimated potential motif space is smaller (<10^4^) than the total number of antibody sequences (>10^14^, (*5, 90*)) by at least 10 orders of magnitude. Our dataset captures a substantial portion of the total motif space indicating the restriction of the paratope-epitope interaction motif space.

### Interaction motifs have a unique sequential amino acid signature indicating immunological function

Since structural interaction motifs retain association with their underlying sequence, we were able to ask whether structural interaction motifs group paratope and epitope sequences with common sequence signatures. If so, it would indicate that structural interaction motifs bear distinct immunological and biochemical function. To investigate the sequence dependencies within selected multi-residue (length>1) paratope and epitope interaction motifs, we determined the 2-mer decomposition of the sequences mapping to the four most abundant paratope/epitope sequences (see Methods). We visualized thus computed sequential motif dependencies (for 4 interaction motifs, top shared 3 from both paratope [`XXX, XX, X1X`] and epitope [`XXX, XX, X2X`]) as Chord diagrams where the 20 amino acids form the segments in a track (the outermost ring) and the links indicate the frequency with which a 2-mer sequential dependency occurred (Fig. 2E).

Visually, paratope and epitope motif sequential signatures differed from one another (Fig. 2E). To quantify this difference, we showed that sequential dependencies were largely non-overlapping (Fig. 2G) and if overlapping, were numerically different (Fig. 2F). Numerical differences were statistically confirmed for all four tested paratope motifs and three of the four epitope motifs (see caption of Fig. 2G for details).

Finally, we compared the paratope and epitope sequence signatures to those of non-immune protein-protein interaction (PPI) and found that immune and non-immune interaction motifs each have distinct sequence dependencies for the same motifs (Suppl. Fig. S13).

In summary, our data suggest that each motif, in paratopes as well as epitopes, has a unique sequential amino acid signature, which indicates that structural interaction motifs cluster together sequences in a non-random fashion that bears immunological function. Supporting this claim is that sequence dependencies discovered in paratope-epitope interaction cannot be derived from non-immune protein-protein interaction.

### The structure of paratope and epitope motifs differs across CDR and FR regions

Next, we asked whether interaction motifs differ structurally. To address this question, we measured the “angle” of each motif. The angle of an interaction motif was computed by defining two vectors spanning the midpoint of a motif and its start and end positions. Larger angles would indicate that the structure of the motif is more extended whereas small angles indicate that the motif would manifest as a loop (see more details in the *Methods* section). In general, median epitope motif angles were only maximally as high as paratope motif angles. We found that median motif angles in heavy and light chain of CDR3 did not substantially differ between paratopes (V_H_ 57° and V_L_ 58°) and epitopes (V_H_ 61° and V_L_ 62°), however, the difference is statistically significant indicating that epitopes assume a more extended conformation (Suppl Fig. S8A). This observation agrees with previous studies that found epitopes prefer to localize on the planar parts of the antigen (*91*) manifesting as flat oblong elliptical shapes (*32*), at least for the CDR3 region. Differences of CDR motif angles among regions were more apparent for example for paratope (median): CDR-H1: 130°, CDR-H2: 42.5°, CDR-H3: 57°, reflecting the structural diversity of antibody-antigen interaction. Specifically, CDR-H2 and CDR-H3 were “loopier” whereas CDR-H1 tended to be linear. In contrast to CDRs, median FR paratope angles were mostly larger than epitope angles. However, since the data points were much sparser for FR compared to CDR, these observations should be viewed with caution. Paratope and epitope angles correlated moderately positively in the majority of the regions (max r_Pearson,CDR-H1–3_ = 0.57, max r_Spearman, CDR-H1–3_ = 0.55, Suppl Fig. 8B).

To further substantiate our structural analysis of the motifs, we investigated the distribution of backbone dihedral angles (using so-called “Ramachandran plots”, Suppl Fig. S21). Specifically, we compared Ramachandran plot statistics between paratope and epitope motifs, PPI motifs, CDR and FR regions and antibody and PPI interacting residues. In addition to verifying that FR and CDR use different angles (Suppl Fig. 8), we found that PPI mostly manifests as alpha-helix and antibodies mostly manifest as beta-strand/sheet, P_II_ spiral, and delta turn, thus underlining the uniqueness of immune protein interaction.

Taken together, we showed paratope and epitope motifs vary across FR and CDR regions and are structurally distinct from PPI motifs and, thus, to a large extent unique to immune recognition.

### Paratope motifs are shared across antibody-antigen complexes

Since our structural motif encoding of the paratope and epitope interface discriminates between continuous and discontinuous motifs, we asked to what extent paratopes or epitopes are continuous or discontinuous. To this end, we quantified both the number of continuous and discontinuous motifs for each FR/DDR region (Fig. 3A) and the number of complexes that share identical motifs (Fig. 3B). We found that: (i) CDR-H3 is an obligate region for antibody-antigen interaction because the CDR-H3 is the only region that had interacting residues in each antibody-antigen complex investigated whereas all other CDR/FR regions were at least partially dispensable for antigen interaction, (ii) paratope motifs are predominantly continuous (except in CDR-H2, CDR-L1, and CDR-L3) and (iii) continuous paratope motifs, more so than discontinuous, are shared across antibody-antigen complexes; epitope motifs exhibited substantially less sharing and were more discontinuous. Specifically, we found only ten paratope motifs (`X, XXX, XX, X1X, XXXX1X, XXXX, XXXXX, X2X, XXXXXX, XXX1X,` Suppl Fig. S4H) and five epitope motifs (`X, XX, X1X, X2X, X3X,` Suppl Fig. S4I) were present in at least 10% (or 82 in absolute numbers) or more complexes. Six out of the ten top-shared paratope motifs were continuous, all of which are no more than seven residues long (including gaps) indicating that short motifs mediate a substantial proportion of antibody-antigen interactions. Indeed, the 20 top shared paratope motifs made up 78% out of all encountered motifs and the 20 top shared epitope motifs made up 58.5% (Suppl Fig. 4D). Importantly, shared paratope interaction motifs were not specific to a given class of antigens (Suppl Fig. S4F), nor to a specific germline gene (Suppl Fig. S4G) and 13 of the most shared interaction motifs were also found in HIV-bNAb (Suppl Fig. S5B) underlining the generality and diversity restriction of the interaction motifs here investigated.

**Figure 3.**
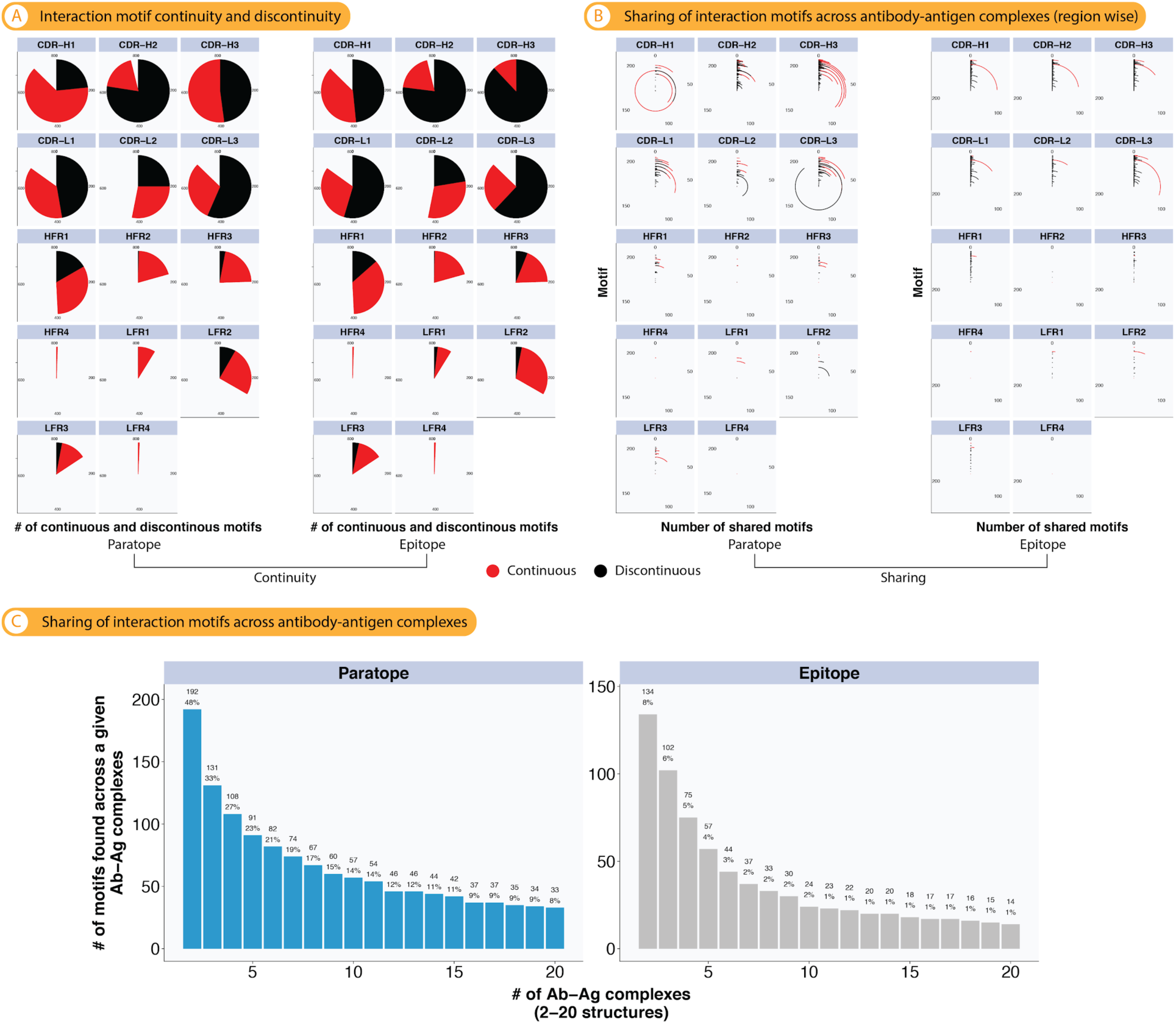
Paratope interaction motifs show a higher extent of continuity and sharing across antibody-antigen complexes than epitope interaction motifs. (A) Ratio of continuous (absence of non-interacting residues) and discontinuous (presence of at least one non-interacting residue/gap) paratope and epitope interaction motifs across antibody-antigen complexes. For example, for paratope CDR-H3, the pie chart signifies that in ≈50% of the complexes, CDR-H3 motifs are continuous and in 50% discontinuous. Gaps in pie charts indicate that for a given region not all structures showed interacting residues. **(B)** Absolute number of antibody-antigen structures a given interaction motif is found in (by CDR/FR). **(C)** Absolute and relative number of motifs found across at least 2–20 antibody-antigen complexes (x-axis).

**Figure 4.**
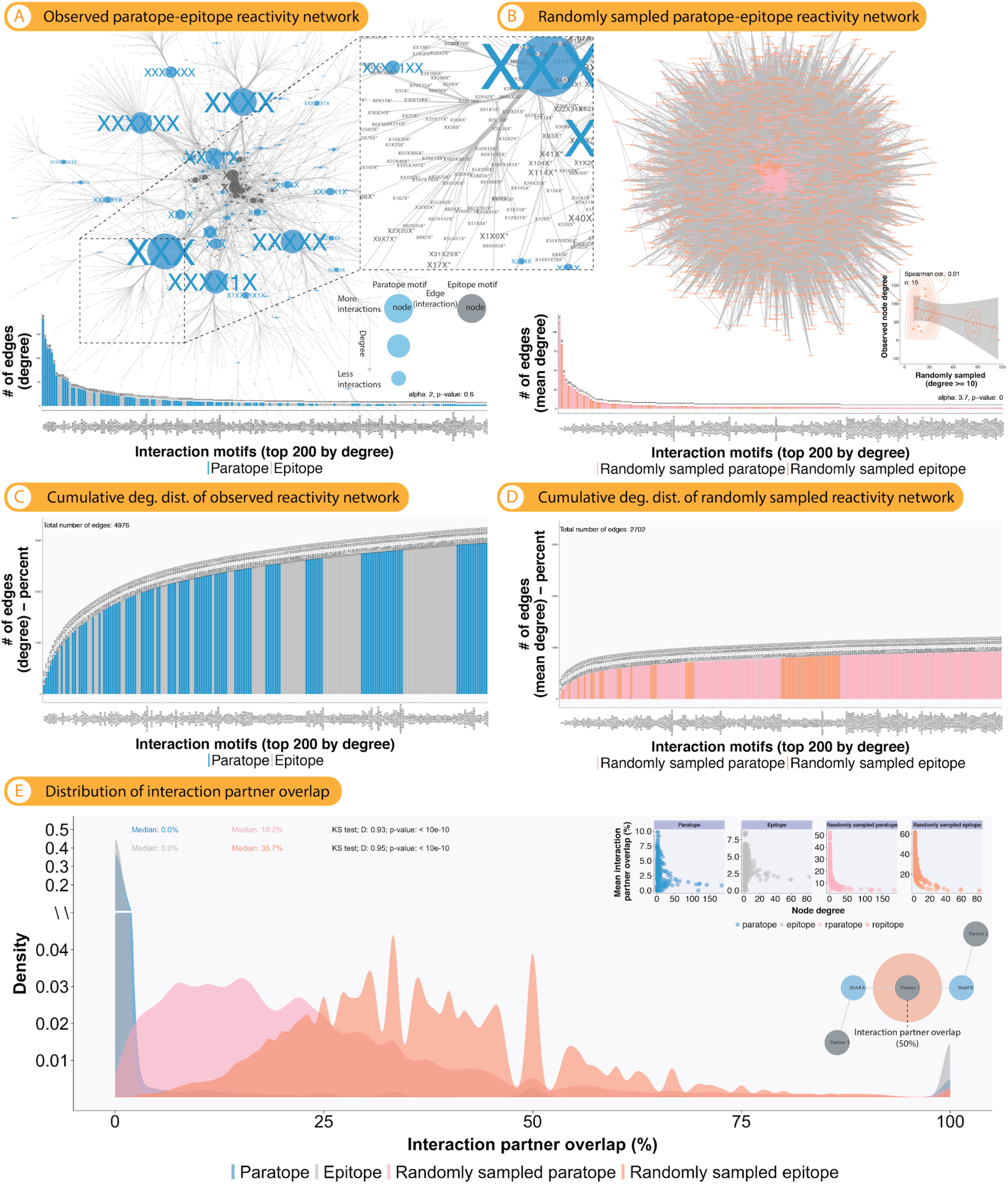
The majority of paratope and epitope motifs are oligoreactive while a small number of paratope motifs show broad polyreactivity towards distinct epitope motif spaces. **The reactivity network of paratope-epitope interaction motifs indicates a priori predictability of antibody-antigen binding. (A)** A bipartite reactivity network capturing paratope-epitope motif interaction was constructed by connecting each paratope motif to its corresponding epitope motif (undirected edge) – given that paratope and epitope motifs may occur more than once across antibody-antigen structures, paratope and epitope motifs may have multiple network connections. Network vertices were scaled by their number of connections (degree). Only the largest connected portion of the network was visualized (for full network, see Suppl Fig. 16). The network degree (a node’s degree is the count of its connections from other nodes) distribution was tested to fit a power-law distribution by calculating a goodness-of-fit value with bootstrapping using the `R` package `poweRlaw` (*65*) as described by Clauset and colleagues (*66*). Here, a network whose degree distribution fits a power-law distribution with an alpha [exponent] between 2 and 3 is defined as scale-free (*67*). A p-value higher than 0.1 means that a power law cannot be ruled out. Inset shows a zoomed-in section of a paratope motif (blue) connecting to a diverse set of epitope motifs colored in grey (polyreactivity). **(B)** To confirm that the reactivity network architecture observed in (A) was unlikely to be observed by chance, we randomly sampled 100 times 1000 motifs from paratope and epitope motif distributions (Suppl Fig. 4D,E). Inset: The Spearman correlation of node degree correlation of observed and randomly sampled networks is shown. For both networks, the respective node degree distribution is shown (for B, the standard error of the mean is also shown). The power-law fit was done as described in (A). **(C, D)** Cumulative degree distributions of networks (A) and (B**). (E)** Distribution of interaction partner overlap for networks (A,B). Briefly, for example for all paratopes in (A), the pairwise overlap of bound epitope motifs was calculated. The statistical significance of the difference between overlap distributions from (A) and (B) was computed using the Kolmogorov-Smirnov (KS) test. Inset: the correlation of node degree and interaction partner overlap was determined.

More generally, 8% of all paratope motifs were shared across at least twenty complexes where that was only the case for 1% of the epitopes (Fig. 3C). Epitopes were thus similar in sharing behavior to PPI motifs were only 3% were shared across twenty or more complexes (Suppl Fig. S18C).

### A selected number of paratope motifs shows broad polyreactivity towards mutually exclusive epitope motif spaces demonstrating a priori predictability of antibody-antigen binding

We next asked whether paratope and epitope motifs have preferred motif partners which would indicate a priori predictability of paratope-epitope binding or whether their interaction is mostly random. To answer this question, we constructed a paratope-epitope-motif network (a bipartite graph) by connecting each epitope motif to its cognate paratope motif (Fig. 4A, given that paratope and epitope motifs may occur more than once across antibody-antigen structures, paratope and epitope motifs may have multiple network connection). We term such a network a *reactivity network.* In this network, we found that the top 7 connected motifs were paratope motifs (mostly continuous or with one gap) that made up 17% of all connections in the network (Fig. 4A,C). Together, these top 7 paratope motifs made 829 connections to predominantly different epitopes (Fig. 4E, inset) thereby collectively binding to ≈50% of all unique epitope motifs (Fig. 2D). Thus, while these paratope motifs showed broad polyreactivity, they bound to largely entirely different epitope motif groups. This finding is in line with the fact that each motif occupies a different sequential dependency space (Fig. 2E–G). Apart from highly connected paratope (and very few epitope) motifs, paratope and epitope motifs were lowly connected and thus specific with respect to their interaction partners. Indeed, we found that the degree distribution of the reactivity graph was power law-distributed and scale-free (p-value = 0.6 (≥0.1), alpha = 2) (*66*). To examine whether the connectivity patterns observed were simply due to the fact that there are more epitope than paratope motifs, we generated control reactivity networks by randomly sampling 100 times 1000 paratope and epitope motifs and analyzed the networks’ degree distributions (Fig. 4B). In contrast to the observed reactivity network (Fig. 4A), the reactivity network obtained from random sampling showed (i) a more even distribution of connectivity for both paratopes and epitopes that was not power law (p-value<0.1) and with lower maximal node degrees and (ii) an increased overlap in bound partner motifs and thus significantly lower specificity (Fig. 4B,D,E). We also excluded the possibility that the observed reactivity network originated from the underlying frequency distribution of motifs (Suppl Fig. S4) as motifs that occur most often are not those that are mostly connected (Suppl Fig. S19A).

Furthermore, we asked whether the reactivity network observed for antibody-antigen interaction was immunity-specific. Therefore, we drew the PPI motif reactivity network for all available non-immune protein structures (that were of similar size to antibody-antigen structures) (>25,000 motif pairs corresponding to >10,000 3D structures, see Methods). We found that PPI reactivity network degrees are exponentially distributed with a lower slope (both on the entire network and a network subsampled to the size of that of the antibody-antigen dataset), meaning less motif hubs dominate interactions (Suppl Fig. 12). Thus, similarly to earlier observations in motif sequential dependencies and the amino acid usage, paratope-epitope reactivity characterizes specific properties and cannot be derived from generic protein-protein interactions.

To summarize, the top-connected motifs in the reactivity network show polyreactivity towards epitope spaces that are non-overlapping (polyreactive specificity). Most motifs, however, are oligoreactive (low number of partner interaction motifs) and thus highly specific (oligoreactive specificity). The combined high specificity and distinctiveness of paratope-epitope interaction indicates that paratope-epitope binding is a priori predictable. The observed reactivity network is fundamentally different from networks constructed from randomly sampling the observed paratope and epitope degree (connectivity) distributions and networks derived from generic protein-protein interaction.

### Quantification of machine-learnability of paratope-epitope interactions

The paratope-epitope reactivity map indicates a priori predictability of antibody-antigen binding. To quantify the accuracy (learnability) with which one can predict (translate) one paratope interaction motif (or sequence) into the cognate epitope interaction motif (or sequence) and vice versa, we leveraged both shallow and deep learning. Specifically, in the shallow model, we used the probabilities of output motifs (or sequences) and a prior we derive from marginal output probability in order to predict the output motif. The deep model leveraged deep learning-based Neural Machine Translation trained on pairs of paratope-epitope interaction motif/sequence (Fig. 5, bottom panel). Unlike the shallow model wherein the probability of a (whole) motif/sequence was inferred given its partner, the decoder part of our deep model generates each motif/sequence character by character (generative model). We rationalized the use of Neural Machine Translation, a deep-learning sequence to sequence model, due to the fact that the prediction of one motif (or sequence) from another is inherently a sequence to sequence translation problem and due to the fact that we observed sequential amino acid signatures in our dataset. We evaluated model performance by comparing (i) the magnitude (length) and (ii) edit distance (error) of the predictions and the true motifs or sequences (see Methods).

We separated different prediction tasks (use cases) by encoding: motif, motif and position, sequence (residues), and overlay of sequence and motif (aggregate). A total of eight use cases were trained depending on the input information (epitope prediction from paratope and vice versa on the motif, sequence and aggregate level, see Table 1 and Fig. 5 and Methods Section for further details). For each use case, the baseline prediction accuracy was calculated based on label-shuffled data where we trained models on randomly permuted paratope-epitope pairs with identical parameter combinations to the original translation machine learning tasks (Suppl Fig. 17, Fig. 5). The accuracy of each method and use case was evaluated by the average Levenshtein distance between the model output and the expected output. Nota bene, this approach for quantifying the prediction accuracy is similar to a multi-class prediction scenario and thus more challenging that the wide-spread binary binder versus non-binder classification. As a more coarse-grained measure for learnability, we also verified whether the output motif length was consistent with the expected output (applicable only to deep learning) (Suppl Fig. 17).

**Figure 5.**
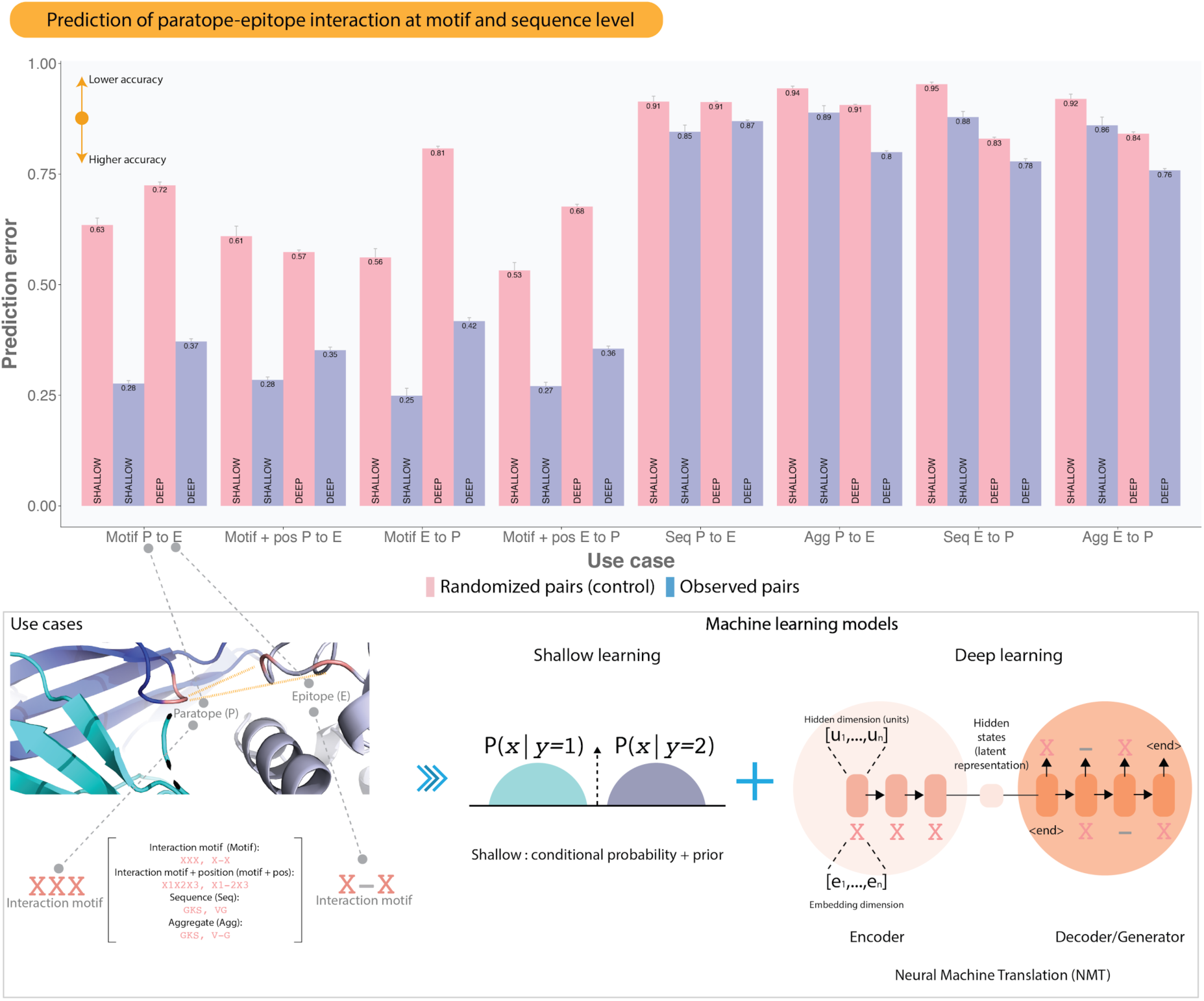
Quantification of machine-learnability of paratope-epitope interactions at motif and sequence level. **(Bottom)** Schematic of the paratope-to-epitope and epitope-to-paratope prediction task (use cases). To quantify the learnability of antibody-antigen interactions at motif and sequence levels, four distinct use cases were used: (1) interaction motif, (2) interaction motif with positional index, (3) sequence, and finally (4) motif and sequence aggregate (all use cases are explained with examples in the Methods section). We leveraged both deep and shallow machine learning approaches. Briefly, the shallow model takes into account the conditional probability of the output with respect to the input, but also the prior corresponding to the output with the highest marginal probability. In deep learning, pairs of input-output sequences were translated via a combination of two components: encoder and decoder with gated recurrent units. Unlike the shallow model wherein the probability of a (whole) motif/sequence was inferred given its partner, the decoder part of our deep model generates each motif/sequence character by character (generative model). The hyperparameters *hidden dimension* (1, 2^1^, 2^2^,…, 2^10^) and *embedding units* (1, 2^1^, 2^2^,…, 2^10^) were optimized during the 20 epochs-long training. A total of 121 models were trained for each use case with 10 replicates (see Methods for details). (**Top**) The median prediction error was obtained by calculating the median Levenshtein distance between the output and the predicted output for each use case across all parameters. The distance ranges from 0 (perfectly matching output-predicted-output, high prediction accuracy) to 1 (fully dissonant output-predicted-output pairs, low prediction accuracy). Here shown is the mean of the medians from the replicates of each use case. Use cases cover the bidirectional prediction tasks (paratope to epitope as well as epitope to paratope) of motif to motif, motif with position to motif with position, and finally amino acid sequence to amino acid sequence (see Table 1). Baseline prediction accuracies (control) were calculated based on label-shuffled data where antibody and antigen-binding partners were randomly shuffled (randomized pairs). Total unique pairs for motif, sequence, and aggregate levels are 2847, 3967, and 3986, respectively. Error bar: ± 2×standard error).

As the deep models scale with the complexity of the parameters (hidden dimension and embedding dimension), we observed an increasingly positive correlation between the predicted lengths and the lengths of the true motifs (sequences) (Suppl Fig. 17A–B, r_pearson_ 0.8–0.9). In contrast, models trained on randomized paratope-epitope pairs failed to recover the correct length indicating that the length for the motif or sequence is predictable. Although the length was learnable to a similar extent in both interaction motif and sequence task (Suppl Fig. 17), the sequence content (indicated by edit distance/error) is not equally predictable (Fig. 5, top panel).

Specifically, for both shallow and deep learning models, the medians of prediction error of interaction motif use cases 0.25–0.42 (accuracy 58–75%) were substantially lower than those of sequence 0.78–0.87 (accuracy 13–22%) use cases. The medians of prediction error for the control baseline use cases ranged from 0.53–0.81 (accuracy 19–47%) and 0.83–0.95 (accuracy 5–17%) for interaction motif and sequence use cases respectively. Together, these results indicate that the paired paratope-epitope interaction motif space reaches reasonable accuracy whereas the sequence space remained challenging to predict. We observed similar trends – prediction accuracy at interaction motif level is higher than at sequence and levels – when examining protein-protein interaction data (Suppl Fig. S20).

Given structural interaction motifs represent one of the layers of antibody-antigen binding, we asked whether integrating motif and sequence information improves sequence-based prediction. Indeed, when combining structural motifs and sequences to an “aggregate”, the prediction accuracy of the deep model, but not shallow models, improved by 2–7 percentage points as compared to the sequence-only use case (Fig. 5). Thus, adding structural information improves the sequence-based prediction accuracy of antibody-antigen binding as it removes interaction ambiguity from the paratope-epitope reactivity space (Suppl Fig. S19B, C).

## Discussion

### Summary

We set out to determine whether antibody-antigen binding is predictable. To this end, we performed an unbiased search for binding fingerprints in the largest available set of curated antibody-antigen structures and discovered a compact vocabulary of antibody-antigen interaction in the form of structural interaction motifs. We characterized this vocabulary by combining a (i) structural interaction motif language, (ii) network theory, and (iii) shallow and deep learning, we found that there exist paratope and epitope interaction motifs that are shared across antibody-antigen complexes. These motifs are predominantly simple (short and continuous), immunity-specific and their sequence diversity is restricted. We showed that each motif has unique sequential dependencies indicating that our motif definition captures underlying immunological principles. While most paratope and epitope motifs showed high specificity, we discovered a subset of paratope motifs with broad polyreactivity towards distinct epitope spaces thus partitioning the paratope-epitope interaction space. The structure of paratope-epitope reactivity network indicated a priori predictability of antibody-antigen binding. We quantified paratope-epitope interaction predictability at motif, sequence, and motif+sequence aggregate levels using shallow and deep learning. Importantly, combining the information of paratope motif and sequence improved predictability (albeit only slightly). This shows that the motif vocabulary is a valuable feature for the development of paratope-epitope prediction tools. To provide quantitative robustness to all of our findings, our study contains, to our knowledge, the most comprehensive statistical evaluation of antibody-antigen and non-immune protein-protein binding to date. Specifically, our study contains the largest volume of information on (i) the distribution of amino acid residues at the binding interface, (ii) the extent of binding interface (dis)continuity, (iii) quantification of interaction complex sequence similarity, (iv) a range of structural definitions of paratope and epitope interaction and (v) the relationship between somatic hypermutation sites and paratope-epitope contact residues. Together, our results demonstrate the existence of learnable sequence and structural rules in 3D-antibody-antigen interaction. Below, we discuss the implications of our findings for the understanding of antibody specificity and the engineering of epitope-specific antibodies.

### Antibody-antigen interaction operates via structural interaction motifs

In agreement with previous literature, we confirmed that most interacting residues are localized in the CDRs – albeit 10% of antibody-antigen contacts were localized outside the CDRs (Fig. 1C). Previous reports found that up to 20% of the residues lie outside the CDR region (*nota bene*: these percentages are to a certain degree subject to numbering-scheme dependent variation) (*26*). Epitopes were discontinuous to a larger extent than paratopes (Fig. 3), which is in accord with previous literature (*8, 32*). Furthermore, we confirmed the previously observed enrichment of key amino acid residues in paratopes such as tyrosine and serine (Suppl Fig. S2) (*92*). Importantly, the amino acid usage differed most between paratopes and non-immune PPI, whereas epitope amino acid usage was as similar to that of paratopes as to that of PPI (Suppl Fig. S6E).

Building on these confirmatory results, we showed that interaction motifs were shared across entirely different antibody-antigen complexes. The search for fingerprints of interaction has also been recently reported for protein-protein interaction (*93*). Thus, antibody-antigen interaction as a whole may be regarded as modular. Indeed, we found that only the VH-CDR3 region is an obligate region for antibody-antigen interaction. Seemingly, all other CDR and FR regions may be modularly added to fine-tune antibody recognition. The immune system is inherently a policing system that surveys both self and non-self for possible threats. Thus, it requires highly flexible recognition that is difficult to escape from via mutation – simple and small sequence motifs match this purpose. Interestingly, in HIV broadly neutralizing antibodies (bNAb, Suppl Fig. S5), the same motifs as in non-bNAb were used underlining the generality of the motif vocabulary here discovered and characterized (*94*).

The discovery of interaction motifs crucially depended on the FR/CDR-focused definition of paratope and epitope (Fig. 1, see Methods) as these are the locales of the fundamental binding units of antibody-antigen binding (In the future, once more antibody-antigen structures become available, one may attempt to search for motifs based on the entire antibody and antigen. A given antibody V_H_ and V_L_ has a median of 3–4 motifs, Suppl Fig. S9). Relatedly, Kunik and Ofran (*31*) showed the six antibody binding regions [ABR] (≈CDR-H/L1–3) differed significantly in their amino acid composition and that each ABR tends to bind different types of amino acids at the surface of proteins (*14*). Van Regenmortel interpreted this as follows *“Because the entire accessible surface of a protein is a continuum of potential epitopes* (*16*)*, it could be argued that it would be advantageous for Abs to bind any protein surface patch without requiring specialized sites of increased stickiness. It seems that antibodies are in fact able to achieve this because they have evolved a set of ABRs where each ABR binds different types of amino acids, while the combined preference of the entire set is for epitopes that are indistinguishable from the rest of the protein surface”* (*14*). While we were able to confirm that paratope-epitope amino-acid level contact maps differ across CDR/FR regions (Suppl Fig. 15), we found that paratope interaction motifs were shared substantially across CDR/FR regions suggesting that binding spaces of CDR/FR regions are not as mutually exclusive as previously thought. Indeed, our reactivity network analysis suggested that binding spaces are partitioned at the motif level and not at the amino-acid level (Fig. 4, Suppl. Fig. S19). Specifically, structural interaction motifs encode geometric information of the local structure and are therefore linked to the angle of folding (Suppl Fig. S8). Thus, linking sequence to motifs and motifs to binding in a 2-step process may connect, in the future, local folding to global specificity. Indeed we found that merging motif and sequence information increased the prediction accuracy of paratope-epitope interaction (Fig. 5).

### Interaction motifs are immunity-specific and not biased to certain germline genes, antigen classes, or species

The 825 antibody-antigen complexes here investigated represent naturally only a mere fraction of the potential antibody-antigen interaction space. Although the collection of these complexes is biased towards antibody-antigen structures of past and current scientific interest, we verified that our dataset, as well as the interaction motifs, were not overly biased towards interaction definition-dependent or interaction-independent confounding factors. Specifically, we demonstrated that (i) our structural definition [distance cutoff-based] for calling interacting residues is robust, (ii) the 825 protein antigens belonged to 84 different antigen classes (Suppl Fig. S11), (iii) shared paratope interaction motifs did not cluster by antigen class but instead were shared across antigen classes (Suppl Fig. S4), (iv) antibody and antigen sequences differed substantially in sequence across antibody-antigen complexes (Suppl Fig. S1), and (v) germline V genes across antibody-antigen complexes and species (human, mouse, Suppl Fig. S1, S4).

Furthermore, we assembled a dataset of 25,921 protein complexes that had motifs of similar size to antibody-antigen data with respect to maximum gap size, to answer the question of whether the motifs in antibody-antigen structures are merely a subset of the larger class of protein-protein interaction (PPI) motifs. Indeed, it had been reported previously that, for example, protein-protein and antibody-antigen interaction do not differ in terms of shape complementarity (*95*) whereas other research suggested that antibody-antigen interaction differs from protein-protein interaction (*24, 41*). Here, we found that paratope and epitope interaction motifs do not represent a subset of PPI motifs but are immunity-specific. Specifically, we found that protein-protein and paratope-epitope interfaces differ in terms of (i) amino acid composition and motif length (Suppl Fig. 6), (ii) sequential dependencies (Suppl Fig. 12), (iii) global reactivity (Suppl Figs. 13,16) and structure (Suppl Fig. 21). Further, we found that sequence-to-sequence prediction accuracy was higher for PPI than for antibody-antigen (Suppl Fig. S20). This finding is consistent with PPI amino acid-level binding preferences (Suppl Fig. S15B) that are more restricted than those of antibody-antigen interaction (Suppl Fig. S15A). The higher predictability of PPI may be a product of evolutionary optimization of protein interfaces whereas the microevolutionary processes of antibody-antigen optimization are occurring on shorter time scales. This hypothesis is supported by the larger size of PPI motifs (Suppl Fig. S6B). Indeed, antibody interaction is mediated by non-structured loops while PPI relies on specific structures encoded in the germline. Nevertheless, this does not mean that antibody-antigen interaction does not underlie evolutionary selection: indeed, we saw that mouse and human interaction motifs were remarkably similar despite differently composed germline gene sets and antigen environments.

Taken together, we validated in orthogonal controls that the rules of antibody-antigen interaction here described are universally shared across antibody-antigen complexes and not merely a byproduct of potential confounding factors. Nevertheless, it is important to acknowledge that each antibody-antigen structure provides only a snapshot of the continuum of interactions occurring between antibody and antigen (*86, 101, 102*). Thus, future NMR-driven kinetic interaction data might reveal that motifs identified for a given antibody-antigen complex are actually part of a “quasi-species” motif space. In this vein, an obvious extension of our approach is the addition of information on antibody-antigen affinity. While reliable affinity information was unavailable for the complexes studied herein, it will be of high interest to augment our interaction motif analysis with affinity data.

### Implications for repertoire-scale antibody specificity studies

Each of the paratope and epitope sequence interaction motifs possessed unique underlying amino acid sequence dependencies (Fig. 2E–G) that are distinct from those found in non-immune protein-protein interaction indicating that structural interaction motifs bear immunological meaning (Suppl Fig. S6, Fig. S13). Furthermore, we showed that the polyreactive paratope motifs contact mutually exclusive epitope-motif spaces (Fig. 4). These findings support the notion that for the prediction of antibody-antigen interaction, both antibody (paratope) and antigen (epitope) information is crucial (*43*). It is important to note, however, that while *epitope*-motif spaces might be mutually exclusive across paratopes, *antigen (collection of epitopes)* spaces might overlap since different epitope motifs may be contacted and located on the same antigen. Once more structures become available, we will need to stratify epitope motifs by antigen class to better understand to what extent epitope motifs are exclusive to a given class. In fact, here we found paratope motifs bound antigens of very diverse antigen classes (Suppl Fig. S4F). These further investigations will elucidate the question of whether paratope sequence motifs may be used as in silico scanning devices for epitope-specific sequence regions in high-throughput antibody repertoire data for at least a subset of medically important antigens. In fact, currently, it is already possible to reconstruct the CDR structural landscape of entire antibody repertoires (*2, 96, 97*). Understanding the diversity and structure of antibody specificity on the repertoire level is also of importance for enhancing the precision of immune receptor-based immunodiagnostics where one of the most pressing current problems is a low signal-to-noise ratio (few antigen-specific sequences within a large pool of unrelated sequences) (*1, 88, 98, 99*). Finally, the combination of repertoire-wide phylogenetics approaches (*100*) with antibody-antigen interaction information may enhance our understanding of the microevolutionary processes involved in affinity maturation.

### Antibody-antigen recognition is overall oligoreactive with islands of high polyreactivity: implications for humoral specificity

We identified not only predefined dependencies *within* paratope and epitope interaction motifs but also higher-order dependencies *among* paratope and epitope motifs. Specifically, we found that paratope-epitope interaction is power-law distributed (Fig. 4) with polyreactivity of few selected paratope motif ‘hubs’, while general oligospecificity of the majority of paratope and epitope motifs. The highly polyreactive paratope motifs were more continuous and contacted mutually exclusive epitope space indicating an overall high degree of humoral specificity already on the motif level and not only on the amino acid level as previously thought. We did not observe a power-law reactivity distribution in either randomly sampled antibody-antigen data (Fig. 4) nor randomly sampled protein-protein interaction (Suppl Fig. S12). In fact, we found that protein-protein reactivity networks showed characteristics of exponential networks – lying in between power law (antibody-antigen) and randomly sampled networks.

Power-law distributions have been repeatedly shown to occur in immune repertoires. We and others previously demonstrated that immune receptor repertoires store information in both the frequency as well as the sequence dimension in the form of power-law statistics (*6, 101, 102*). The interaction data here presented now suggest that paratope-epitope interaction may function as a third power-law governed layer of adaptive immunity. The great extent of organization in the paratope-epitope interaction space may be linked to the substantial antibody sequence bias observed in high-throughput antibody sequencing studies (recent high-throughput antibody sequencing studies (*90, 102–104*) have revealed high predictability and predetermination in both naive and antigen-specific antibody sequences). In future studies, we plan to investigate how sequence interaction motifs depend on the underlying immune receptor sequence generation probability landscape (*5, 105*). While we do not yet understand the evolutionary selection pressures (*14, 106, 107*) nor the functional significance of these power-law distributions in immune repertoires, their existence is a fundamental premise for the predictability of antibody-antigen interaction as they create a learnable space in which adaptive immunity operates.

Humoral specificity describes the capacity of the antibody immune response to selectively target a nearly infinite number of antigens. Given the large number of potential antigens, it is commonly thought that antibody-antigen interaction is very challenging to predict. But if one approaches the challenge of understanding antibody-antigen interaction from a motif perspective, it breaks down the problem into a lower-dimensional task. Indeed, our extrapolation of motif diversity suggests that the potential epitope motif space is small (Fig. 2D). It is tempting to speculate as to the evolutionary advantage of short motifs in antigen recognition (non-immune protein-protein interaction has evolved substantially larger motifs, Suppl Fig. S6B,C). Short motifs decrease the potential escape space for antigens but also render self and non-self recognition more difficult. It would be interesting to investigate in how far, for example, autoimmunity and infections (bacterial, viral) occupy different motif spaces. So far, however, we observed that epitope motifs were shared consistently across antigen classes (Suppl Fig. S4F).

### Predictability and learnability of the paratope-epitope interface

The problem of antibody-antigen binding is regarded generally as too high-dimensional. We conjectured that if a general vocabulary existed, which links paratope-epitope interaction in an unambiguous way, then paratope-epitope interaction would be predictable. The motif-based paratope-epitope reactivity network of antibody-antigen interaction shows (Fig. 4A) that motif-motif interaction is predictable whereas sequence-sequence interaction is inherently difficult to predict (Suppl Fig. S19). We found that such a vocabulary existed (Fig. 4, SupplFig. S19), leading to the conclusion that paratope-epitope pairing is indeed predictable.

To quantify the prediction accuracy (learnability) of paratope-epitope interactions, we leveraged both shallow and deep machine learning. We found that motif-based shallow and deep learning predicts paratope-epitope pairing reasonably well. By superimposing structural information on the paratope and epitope sequences into aggregate encoding, we were able to improve the prediction accuracy of sequence-based antibody-antigen binding since the motif-sequence-aggregates deconvolved the reactivity space (Suppl Fig. S19C). These results suggest that motif and sequence form two complementary layers in the antibody-antigen recognition space that can be leveraged to support sequence-based predictions (*34*).

More generally, paratope-epitope prediction is seemingly a subset of a more general protein-protein interaction (PPI) problem. In the structural bioinformatics field, this task is known as binding site prediction and is typically formulated as the problem of finding the set of residues (or patches) on the protein surface likely to interact (via non-covalent bonds) with other proteins (*108*– *113*). More formally, such a problem can be formalized as a binary classification task in which a model is trained to discriminate binders from non-binders typically at residue or sequence level.

That is, given a sequence VGRAISPRAS, the model would assign a probability to each amino acid signifying its likelihood to bind to a partner residue (Pbind(V) = 0.3, Pbind(G) = 0.05, Pbind(R) = 0.7 etc.) or Pbind(VGRAISPRAS) = 0.6. A target-agnostic approach, such as Paratome (*37*), finds a set of regions from a structural alignment of antibody-antigen complexes and use it to locate similar regions in new sequences. Antibody i-Patch (*41*) utilizes contact propensity data to score each amino acid while at the same time leveraging information from neighboring residues (a patch of resides). To be applicable for antibody-antigen prediction, the original i-Patch (*114*) algorithm, however, had to be adjusted with respect to two central assumptions: (i) multiple sequence alignment (MSA) signifying evolutionary (sequence/structural) conservation in protein-interacting domains and (ii) protein-protein derived amino acid propensity score both of which are less pertinent for antibody-antigen complexes as antibodies (as does surviving antigens) constantly evolve obscuring many forms of conservations, structural and sequence alike. Specifically, CDRs in antibodies manifest as unstructured loops with minimal structural conservation across antibodies and amino acid propensity differs between protein-protein and antibody-antigen complexes. This combination compounds the complexity in learning the rules that govern antibody-antigen interaction and necessitates a unique approach separate from the conventional approaches presently applied in the PPI field. The motifs here discovered fill this missing gap by capturing structural and sequence information in a single notation across antibody-antigen complexes and projecting antibody-antigen interaction on to substantially lower dimensions (10^2^ paratope and 10^3^ epitope motifs) which allowed us to observe conservation from a motif’s perspective. For instance, we showed in Fig. 3B–C and Suppl Fig. 4F–I that motifs are ‘conserved’ (shared) across different antigen classes, V genes, and structures. Tools such as Antibody i-Patch may, for instance, leverage a motif-driven alignment in place of the missing multiple sequence alignment data due to sequence diversity of antibody-antigen complexes. Further, the lower dimension in combination with the resulting reactivity network (at motif level) enabled us to probe beyond a binary (binder or non-binder) setting towards a more nuanced multi-class setting. Specifically, we asked for a given paratope motif the corresponding epitope motif instead of whether the motif binds or not (multi-class setting). Multi-class classification although a few magnitudes more difficult than the binary one was proven quite successful at least at the motif level (Fig. 5). Specifically, we note that transitioning into a higher dimensional multi-class setting, such as the transition between the sequence and aggregate encodings, adversely impacted our shallow model more so than the deep one suggesting that as the class diversity tends to infinity the deep model would increasingly outperform the shallow one.

Beyond target-agnostic approaches, accumulating evidence has demonstrated the utility of integrating the information from the interacting partner in improving state-of-the-art performance (*115*). Townshend and colleagues achieved state-of-the-art performance for the prediction of protein-protein interaction by training a model that comprises two separate convolutional neural networks (one from each interacting partner) and concatenating them to produce the final output (*116*). Similarly, Pittala and Bailey-Kellog used an attention layer on top of two separate convolutional layers (one each for antibody and antigen) to produce superior predictions to target agnostic approaches such as DiscoTope and Antibody i-Patch (*41, 117, 118*). Finally, Deac et al. eclipsed the performance of the target-agnostic Parapred approach by building a model that cross-modally attend antigen residues (*35, 119*). Although much more sophisticated in terms of model complexity and architecture in addition to ‘target-aware’-ness, these models remain anchored to the problem of delineating binders and non-binders (binary prediction) and have yet to venture to a multiclass setting. We note as well that antibody-epitope prediction is typically treated separately: (i) to predict residues or sequences in antibodies that bind to epitopes (paratope prediction) and (ii) to predict residues or sequences in antigens that bind to paratopes (epitope prediction). In this dichotomy, paratope prediction typically fares several folds better than epitope prediction (for context, a state-of-the-art predictor from Pittala and Bailey-Kellog yielded areas under the precision-recall curve AUC-PR of 0.7 and 0.212 for paratope and epitope prediction respectively). Au contraire, we observed a notably less dramatic difference at least at the motif level where accuracy ranges from 0.63–0.72 and 0.58–0.75 for paratope and epitope prediction respectively (Fig. 5). Thus, we speculate that ‘motif-awareness’ may further extend the performance of these approaches similar to the added benefit of target-awareness earlier described and bridge the dichotomy between paratope and epitope prediction.

Finally, the paratope-epitope reactivity network (Fig. 4, Suppl Fig. S19) allowed us to draw several conclusions and future directions with regards to the prediction of paratope-epitope binding. Given that there exist a few paratope motifs with broad epitope motif reactivity, the motif-based prediction accuracy of paratope-epitope interaction cannot reach, by definition, 100%. However, as the epitope reactivity of the polyreactive paratopes was mutually exclusive (distinct), focusing prediction efforts on branches of the paratope-epitope reactivity network may improve the performance of sequence-based paratope-epitope prediction models – especially since we discovered that (i) motifs possess unique and distinct sequential dependency signatures and that (ii) motifs aid sequence-based prediction of paratope-epitope pairing (sequence-motifs aggregates, Fig. 5). Thus, the motif vocabulary of antibody-antigen interaction helps partition the high-dimensional antibody-antigen interaction space into smaller, less daunting prediction tasks thereby rendering antibody-antigen interaction learnable.

### Structural interaction motifs provide ground truth for benchmarking of immune receptor-based machine learning

Adaptive immune receptor repertoires represent a major target area for the application of machine learning in the hope that it may fast-track the in silico discovery and development of immune-receptor based immunotherapies and immunodiagnostics (*1, 120–123*). The complexity of sequence dependencies that determine antigen binding (*124, 125*), immune receptor publicity (*47*) and immune status (immunodiagnostics) (*88, 126*) represent a perfect application ground for machine learning analysis (*47, 98, 122, 127–130*). As we extensively reviewed recently (*1*), the development of ML approaches for immune receptor datasets was and is still hampered by the lack of ground truth datasets. Ground truth datasets are defined by the property that the link between the class of a sequence (class = disease/antigen specificity) or repertoire and the underlying sequence structure is known *a priori*. Thus, by definition, ground-truth datasets are those that are synthetically generated in silico (*131, 132*). One of the current bottlenecks for generating nature-like synthetic datasets for predicting, for example, antigen-binding from the immune receptor sequence alone is the lack of knowledge on the complexity of paratope and epitope interaction motifs. Our research resolves this important knowledge gap. Specifically, our atlas of structural interaction motifs now allows for the faithful simulation of native-like ground truth immune receptor datasets for the development of immune-receptor based machine learning methods (*131*). For example, paratope and epitope sequence motifs may be implanted into synthetic 3D antibody-antigen structures (*133*) to subsequently perform benchmarking of different ML architectures with regard to their capacity to (i) predict paratope-epitope binding and (ii) recover the exact sites of antibody-antigen interaction (native-like antigen-specific sequence motifs). A similar approach may be taken for repertoire-based disease classification.

Nota bene: While we firmly maintain that for a profound understanding of antibody specificity, machine learning is of utmost necessity, machine learning alone will not suffice. Indeed, the discovery of general interaction sequence motifs was encoding-*dependent* and machine-learning-*independent*. As previously described at length (*1*), we believe that machine learning approaches adapted to the complexity and intricacies of immune receptor sequence data will have to be combined with (ideally learned) sequence encodings that aim to answer specific questions. In fact, prediction tasks may be aided or complicated by a given encoding. In this work, for example, breaking down antibody-interaction into modular structural interaction motifs has greatly simplified machine learning tasks (Figure 5).

### Implications for machine-learning driven antibody, epitope and vaccine engineering

Monoclonal antibodies are of substantial importance in the treatment of cancer and autoimmunity (*134*). Thus, their efficient discovery is of particular interest. Given that our work is unbiased towards both paratope and epitope analysis, it demonstrated the feasibility of the reconstruction, via generative machine learning, of potential neo-epitopes for neo-epitope design or the discovery of neo-epitope specific antibodies. Our analyses suggest that the number of antibody binding motifs is relatively restricted (Fig. 1–2). Monoclonal antibody discovery is predominantly performed using synthetic antibody libraries. The number of developable hits of such libraries may be increased by tuning sequence diversity towards the interaction motifs (and their corresponding sequential bias) here discovered. Relatedly, engineering-driven computational optimization of antibody-antigen binding, as well as docking algorithms, might benefit from incorporating interaction-motif-based heuristics (*39, 41, 95, 122, 135, 136*). Specifically, if we assume that the interaction motif sequential dependencies discovered here were evolutionarily optimized, they may be used to substitute for the lack of available multiple sequence alignments that are used to calculate high-propensity interacting residues in protein-protein docking (*41*). Furthermore, it will be of interest to investigate whether sequential dependencies are already predictive by themselves as to the antigen targeted (more paratope-epitope-paired data is needed for such investigations).

In contrast to earlier work (*24*), we found that contact residues and somatic hypermutation are in fact correlated (Suppl Fig. S10). For antibody optimization, this suggests that linking the antigen-contacting and somatically hypermutated positions (Suppl Fig. S10) in a high-throughput fashion and predicting whether the paratope prior to SHM was already binding or not, may enable, in theory, the construction of a hierarchy of evolutionary-driving driving SHM sites. Furthermore, it would be of interest to investigate in how far somatic hypermutation preserves binding motifs or, relatedly in how far, a reversal to germline would change interaction motifs. The latter is a particularly important question as there is likely an overrepresentation of high-affinity antibodies in the dataset here investigated.

The here studied antibodies are diverse and can harbor specific structural features such as glycans, for binding envelope proteins of HIV or influenza. Further, those antigens may also harbor glycans depending on the mode of protein synthesis before crystallization. Our dataset therefore inherently already contains the effect of post-translational modifications to antigen-antibody binding. Interestingly, we did not find substantial differences in the motif usage between bNAbs against viral glycoproteins, supporting that the vocabulary of motifs is shared among antigens with diverse structural features in the dataset. However, we cannot tell whether this holds true for proteins that cannot be crystallized or for unstructured loops of antigens, that are typically missing in structural databases.

Finally, in the future, it may also be of interest to correlate interaction motifs with antibody developability parameters (*122, 137–139*). Antibody developability depends on a multitude of parameters that are calculated based on the entire antibody complex (*139, 140*). Thus, all non-interacting residues also contribute to antibody developability calculations. Therefore, future studies will have to delineate to what extent non-interacting residues correlate with specific interaction motifs. For example, for our dataset, we did not observe a correlation between motif usage and germline gene (one proxy for non-interacting residue) usage (Suppl. Fig. S4). And differences in non-interacting residues were less stark compared to interacting residues in terms of amino acid frequencies (Suppl Fig. S2).

### Concluding remarks

Antibodies constitute only half of the adaptive immune system. To our knowledge, a similar work on *TCR*-peptide interaction does not yet exist. Comparing motifs between TCR and antibody-antigen motifs would shed light on mechanistic similarities and differences in antibody and TCR antigen-interaction (*88, 124, 125, 141–145, 145–147*). Furthermore, it remains to be investigated in how far the motif-based rules uncovered here for VH-VL-antigen complexes are transferable to scFV, nanobody, and other next-generation antibody constructs (*53, 148*). Finally, we wish to state that, while one of the main aims of this paper was to advance our quantitative understanding of antibody-antibody recognition, the second main aim was to develop computational approaches that may help study antibody-antigen interaction in the years to come. Indeed, future studies may also investigate other motif definitions (possibly identified by end-to-end machine learning) that may unveil further structure in the antibody-antigen interaction space. Our systems-level approach of combining orthogonal statistical, network and machine learning approaches for the study of antibody-antigen interaction was necessary to reach the conclusions drawn in this work.

*∼ ∼ ∼ Fin ∼ ∼ ∼*

## Acknowledgements

We thank Andrew Macpherson (University of Bern, CH), Anne Corcoran (Babraham Institute, UK), Ludvig M. Sollid (University of Oslo, Norway), Thérèse Malliavin (Institut Pasteur, Paris, France), Marieke Kuijjer (University of Oslo, Norway) and Luca Piccoli (Institute for Research in Biomedicine, Switzerland) for their comments and suggestions that have led to an improvement of the manuscript.

## 1 Supplemental Text. Derivation of an analytical solution for the theoretical number of motifs within one CDR/FR region

### 1.1 Objectives and motif definition

Our aim is to determine the number of possible structural interaction motifs for any motif length. A given sequence motif is defined as follows:

*•* An amino acid is encoded as **X**.
*•* A gap is encoded as integer *n* where *n* quantifies the length of the gap.
*•* Each motif starts and ends with an amino acid **X**.
*•* There can be *>* 1 amino acids in sequential positions but not *>* 1 gaps.

Let us give two different definitions of motif length. By simply “motif length” we mean the number of **X**s in it plus the number of gaps, we note this lengths *L*. By “amino acid length”, we mean the number of amino acids included in the sequence, i.e. the number of **X**s plus the sum of all gap lengths. Please, refer to the section 1.2 for a few examples.

As the interaction sequence cannot exceed the size of the CDR/FR it is located in, we need to add one more constraint:

- *•* The amino acid length of the motif is not bigger then a predefined number.

Let us denote the number of unique motifs of lengths *L* and amino acid length *A* as N*_L,A_* and the number of unique motifs of length *L* with amino acid length *not exceeding A* as N^-^*_L,A_*= 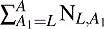

### 1.2 Examples

To derive a formula for N*_L,A_*, we inspect a few examples first for intuition purposes.

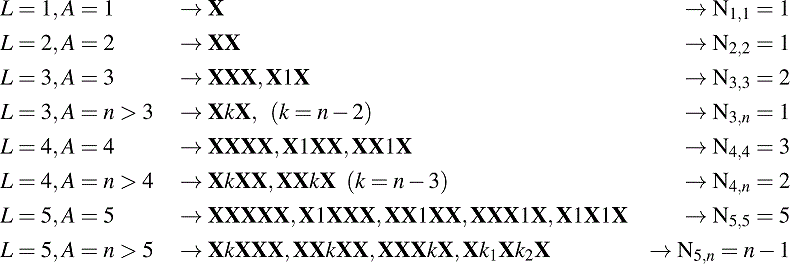

In the last line, *k* = *n -* 4 and *k*_1_ + *k*_2_ = *n -* 3. Let us clarify this last line: there are only 3 motifs with a single gap, but if there are two gaps, their lengths can vary: *k*_1_ = 1*, k*_2_ = *n -* 4; *k*_1_ = 2*, k*_2_ = *n -* 5*,...,* so that we have *n -* 4 double-gapped motifs in total.

### 1.3 General formula

Now we can proceed to derive a general formula for N*_L,A_*. Let us note the number of **X**s in a motif as *n_x_* and the number of gaps as *n_g_*. We can count the motifs for fixed *n_x_* and *n_g_* and then we will just have to sum the results over all *n_x_* + *n_g_* = *L*. Thus, we have *n_x_* **X**s and *n_x_* 1 slots for gaps – between any two neighbouring **X**s there can be a gap. First, we have to choose *n_g_* slots: the number of ways to do this is

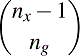

Now we have *n_g_* gaps of total amino acid length *A - n_x_*, and we need to distribute the lengths between the gaps. In other words, we need to split the number *A - n_x_* into a sum of *n_g_* nonzero terms. The number of ways to do this is the number of *n_g_-compositions* of *A - n_x_*, which equals

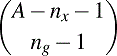

Now we can write down the formula for N*_L,A_* as

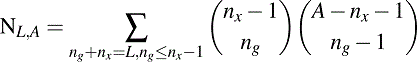

We did not take into account the all-**X** case, so for *A* = *L* we should have

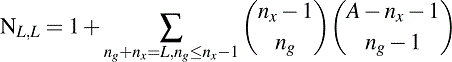

Similarly, the formula for N^-^*_L,A_*is

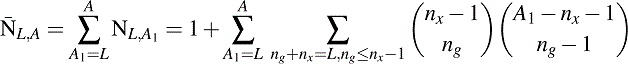

Figures 1 and 2 show the growth of N^-^*_L,A_*for *L* in 1*,...,* 10. We set 10 as maximum motif length based on our observations (Fig. 2B).

**Figure 1:**
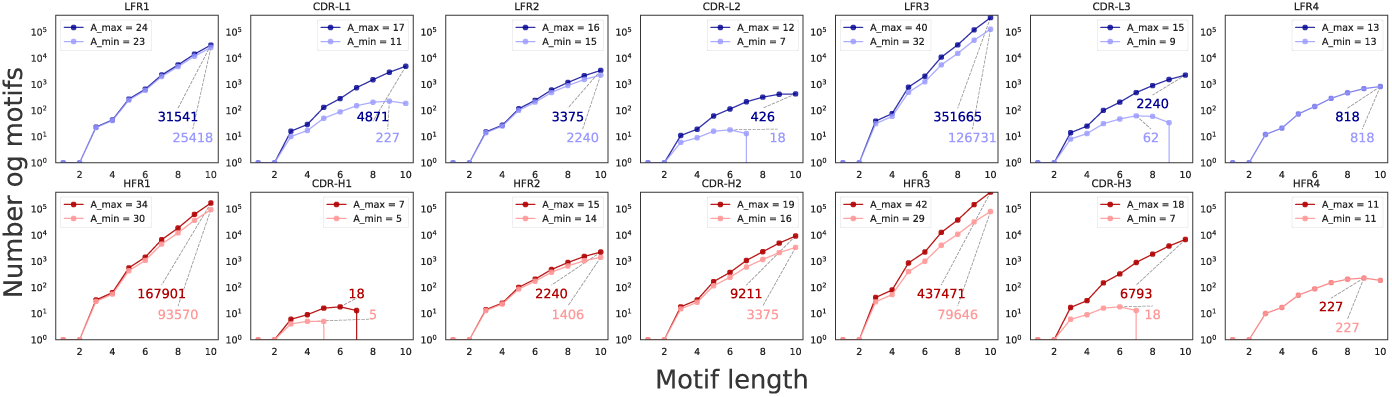
The number of unique motifs (Y axis) for a given motif length (X axis) that could be located in a certain FR/CDR (see possible FR/CDR lengths in Supplementary Table S1). The amino acid length of the motifs is bounded by the minimum and maximum possible region length (Supplementary Table S1).

**Figure 2:**
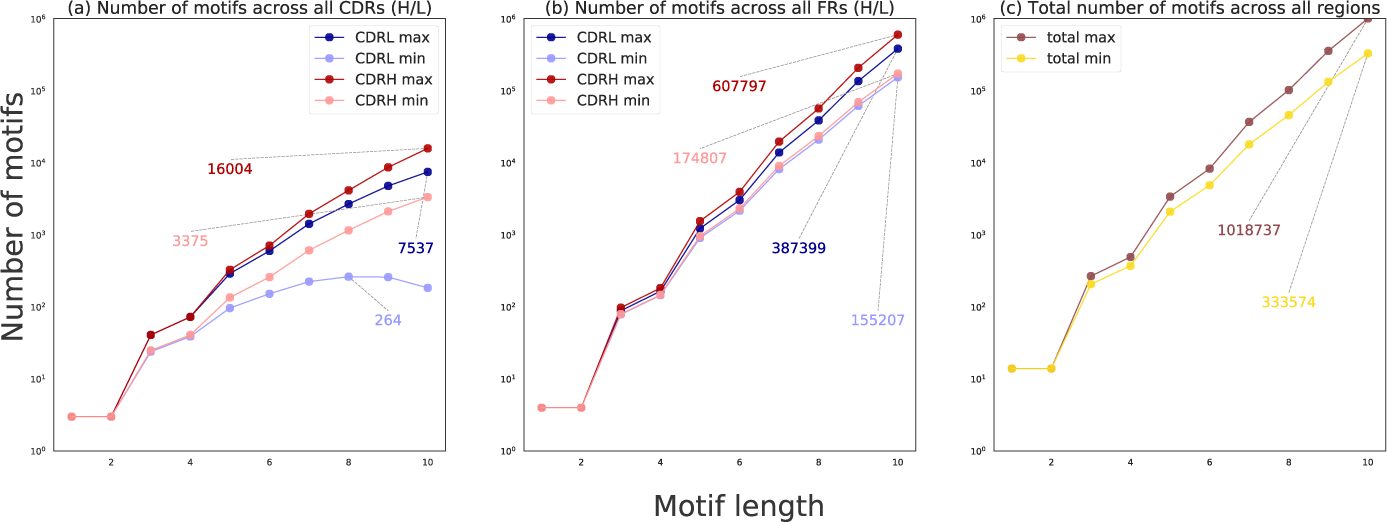
The total number of unique motifs (Y axis) for a given length (X axis) across all CDR-Ls and CDR-Hs **(a)**, across all LFRs and HFRs **(b)**, across all regions **(c)**.

## Supplementary Material

**Supplemental Figure S1.**
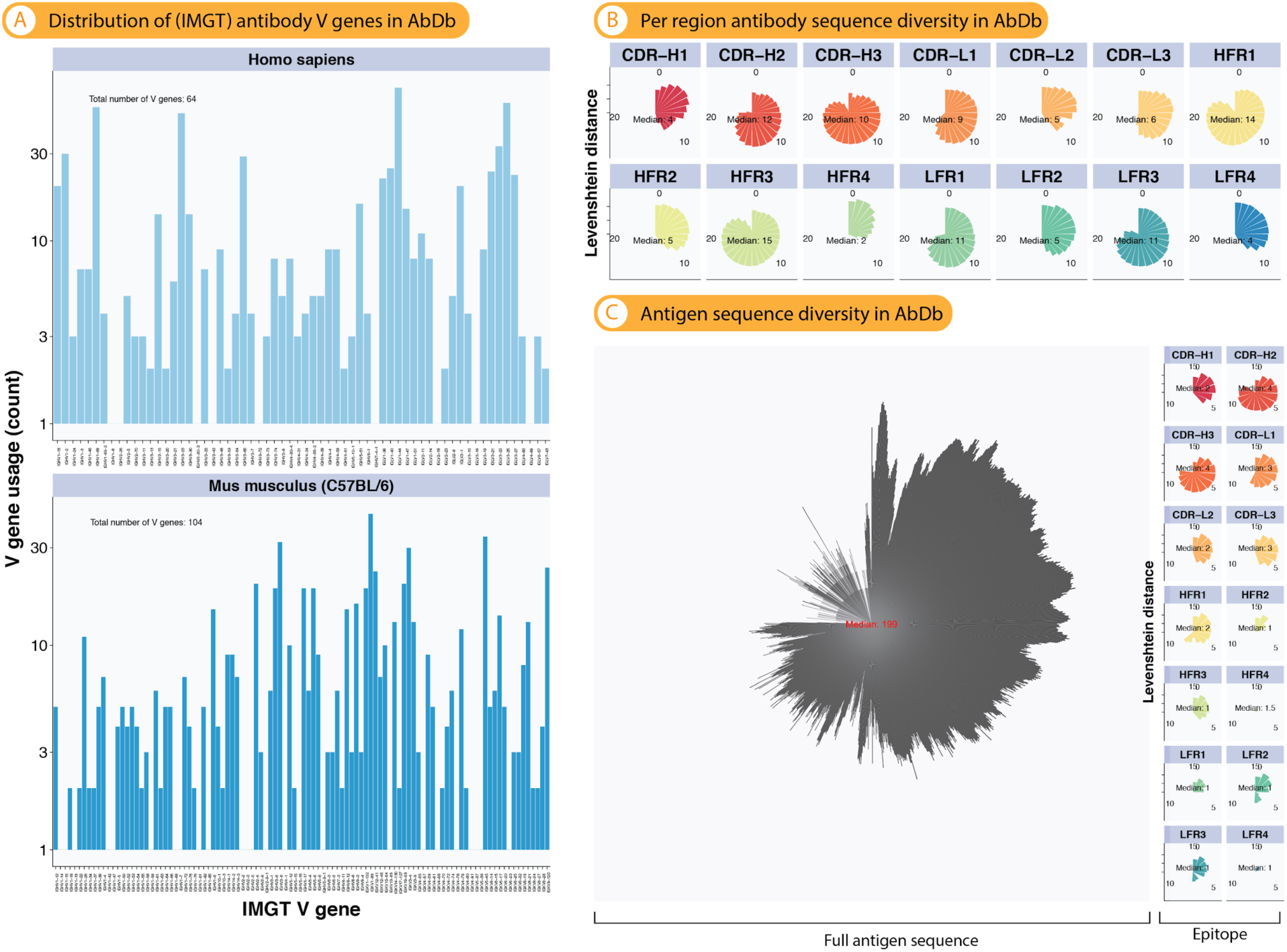
The AbDb dataset covers a broad antibody and antigen sequence diversity. (A) The 825 antibody-antigen structures show a large diversity in germline gene usage (human: 64, mouse: 104) covering a notable portion of the potential V germline gene diversity. **(B)** Distribution of pairwise sequence similarity across antibodies measured by Levenshtein distance. The median distance by region (CDR, FR) ranges between 2 (HFR4) and 14 (HFR1). **(C)** Distribution of pairwise sequence similarity across antigen sequences (both full and by epitope region) measured by Levenshtein distance. The median distance by region (CDR, FR) ranges between 1 (HFR2, HFR3, LFR1–4) to 4 (CDR-H3). The median pairwise distance of full length amino acid antigen sequences is 199.

**Supplemental Figure S2.**
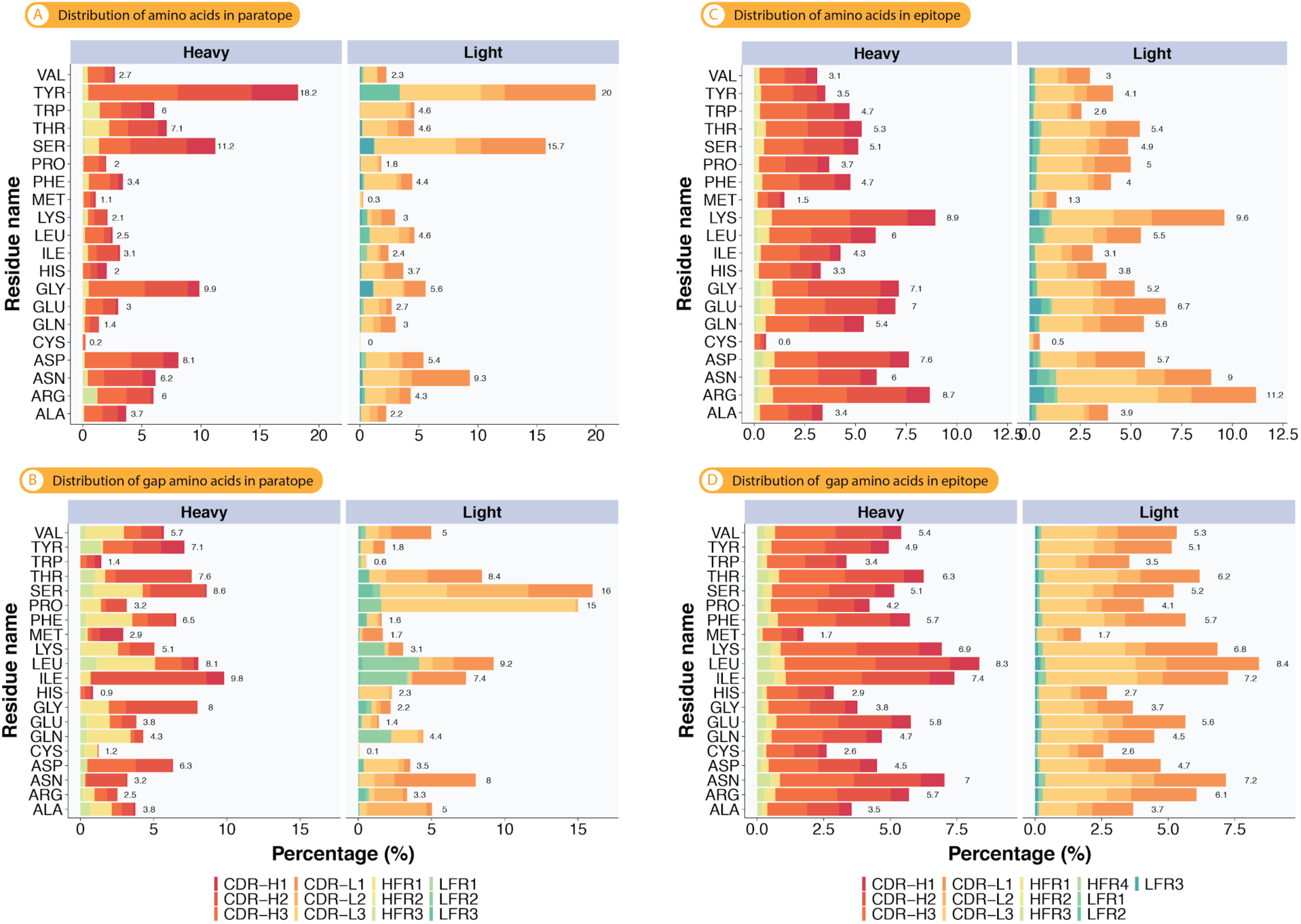
Amino acid residue distributions in paratopes and epitopes as well as gaps (non-interacting residues). (A) Relative residue distribution in paratopes. Aromatic residues, namely tyrosine and serine, are enriched in paratopes. Cysteine and methionine are the least used amino acids. **(B)** In contrast, gap residues in paratopes are more evenly distributed and generally enriched by a different set of residues. **(C)** In epitopes, polar and charged residues such as lysine and arginine are the two most frequently observed residues. In addition, epitope interacting residues are more evenly distributed than those in paratopes. **(D)** Similarly to paratopes (B), epitope gap residues are more evenly distributed and enriched by a different set of residues.

**Supplemental Figure S3.**
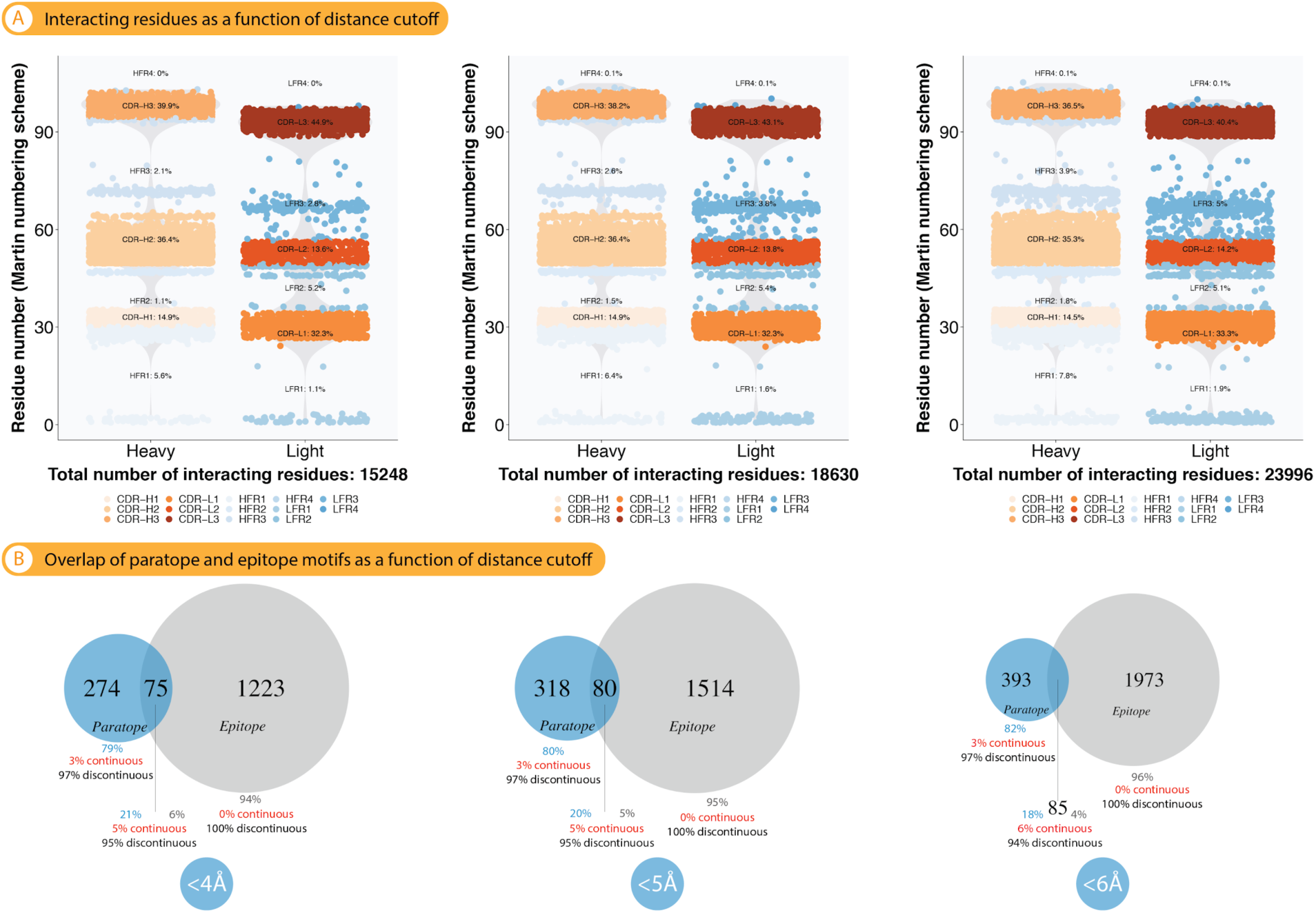
Number of interacting residues and paratope and epitope motifs as a function of the distance cutoff that defines interacting residues. (A) The three panels (columns) show the total number of interacting residues as well as their relative fraction by CDR/FR for the three investigated and commonly used distance cutoffs: <4Å, <5Å, and <6Å. For all analyses in this study, we adopted the distance cutoff of <5Å (see explanation in Methods). The relative distribution of interacting residues across CDR/FR differed only marginally across distance cutoffs. (B) While the number of epitope motifs varied considerably across distance cutoffs, the number of overlapping motifs remained nearly constant.

**Supplemental Figure S4.**
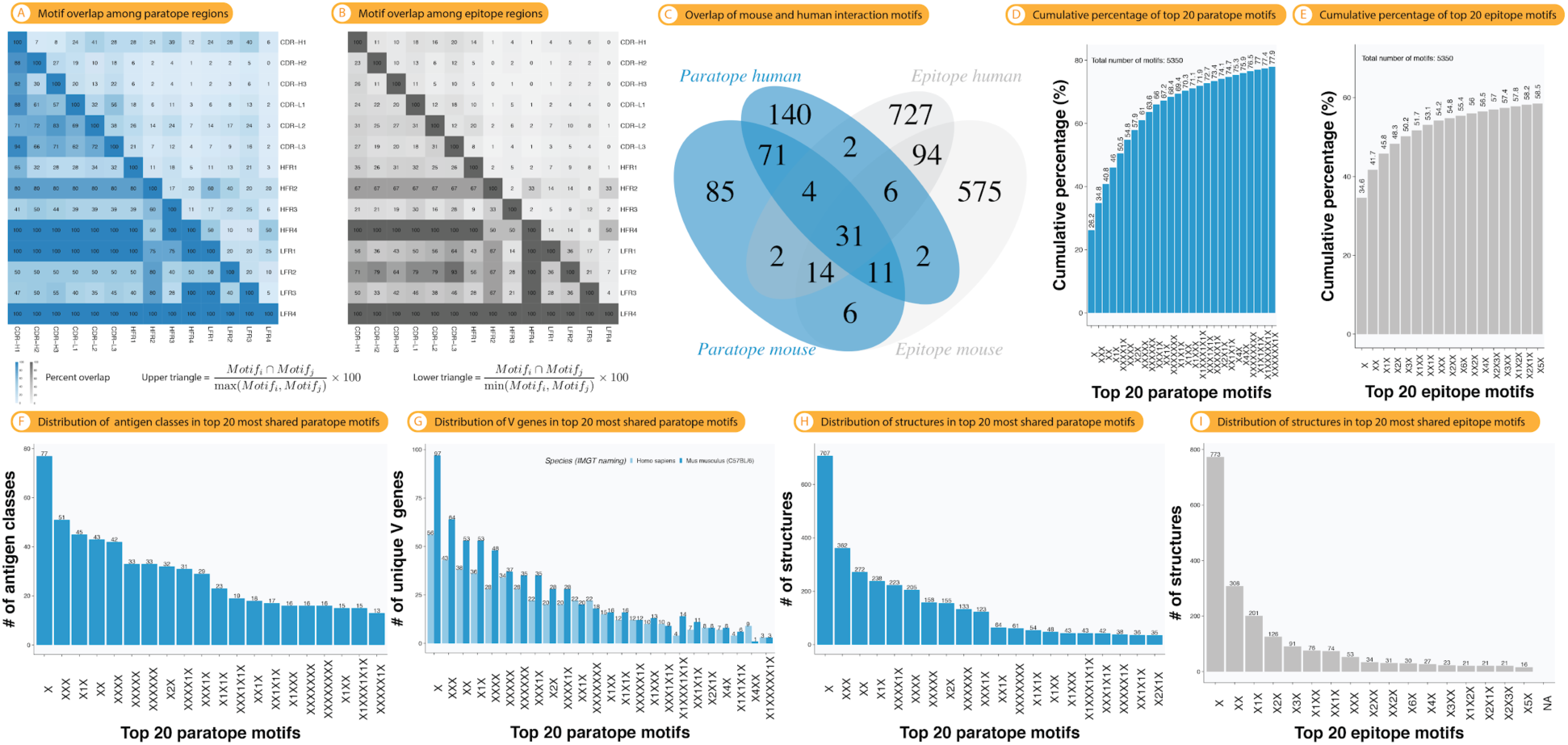
Interaction motifs shared across antibody-antigen structures cover a large antibody and antigen sequence space. Extent of motif sharing (overlap) across antibody regions (CDR/FR) in **(A)** paratopes and **(B)** epitopes. There is considerable overlap across paratope and epitope regions albeit to a slightly lesser extent in epitopes compared to paratopes. **(C)** Overlap of mouse and human paratope and epitope motifs. The cross-species relative overlap is higher for paratopes than for epitopes. **(D)** Cumulative motif distribution of paratopes and **(E)** epitopes ordered by frequency. Motif frequency denotes the number of times a motif has been detected within or across structures. **(F)** Number of antigen classes, **(G)** V genes (separated by human and mouse), and structures **(H)** paratope and **(I)** epitope motifs belong to. The top 20 most shared (across structures) motifs are shown for subfigures E–I.

**Supplemental Figure S5.**
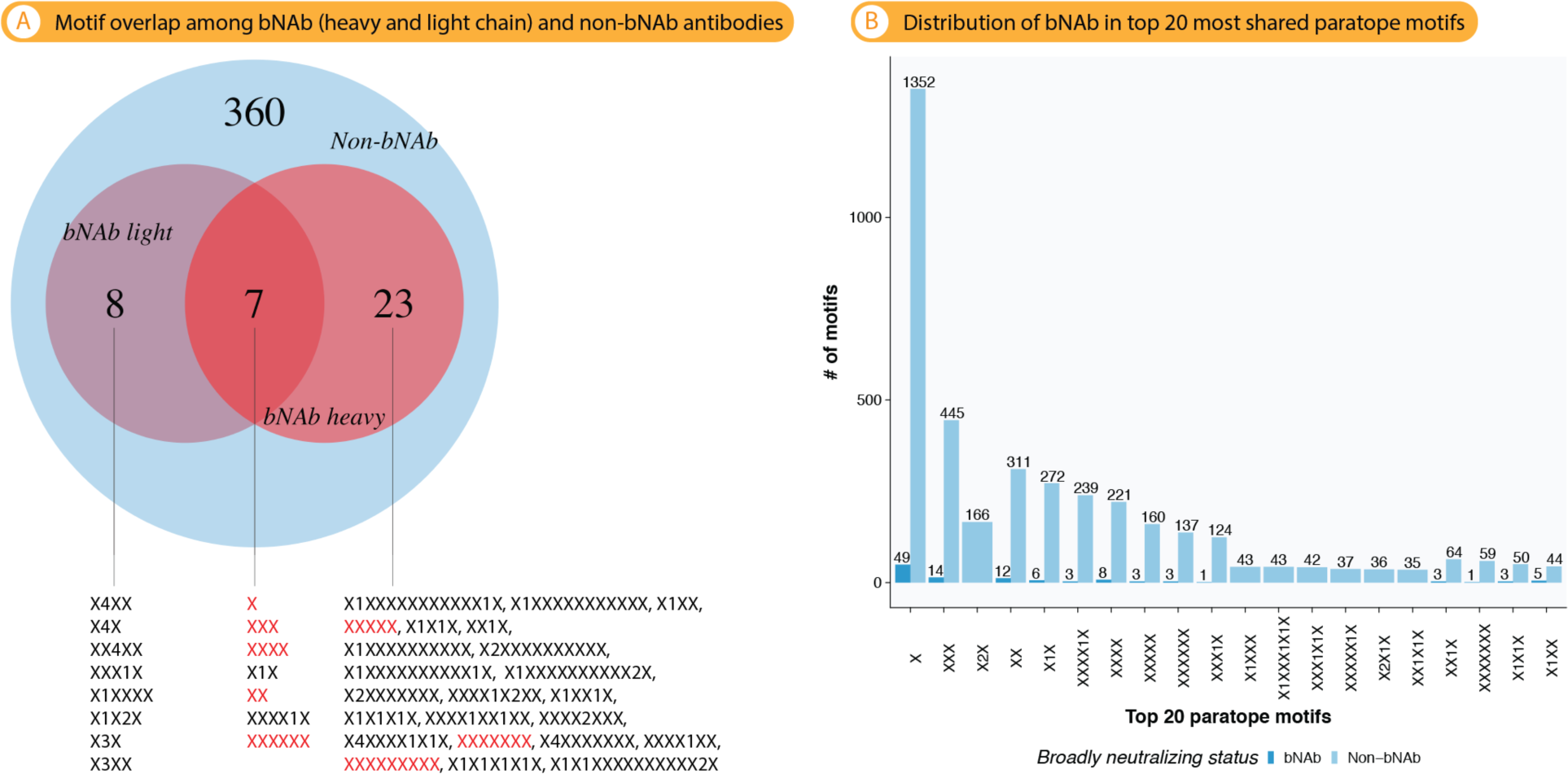
Interaction motifs of broadly neutralizing antibodies (bNAb) do not differ from those found in non-nbNAb antibodies. (A) All 38 motifs identified in the 24 HIV-bNAB structures from the bNAber database (*54*) (see Methods) are also found in non-HIV-bNAb structures. Out of the 38 unique interaction motifs identified, 8 are continuous and 30 are discontinuous. **(B)** Out of the 38 bNAb interaction motifs, 13 motifs were part of the 20 non-bNAb motifs that were shared most across structures (Suppl Fig. S4).

**Supplemental Figure S6.**
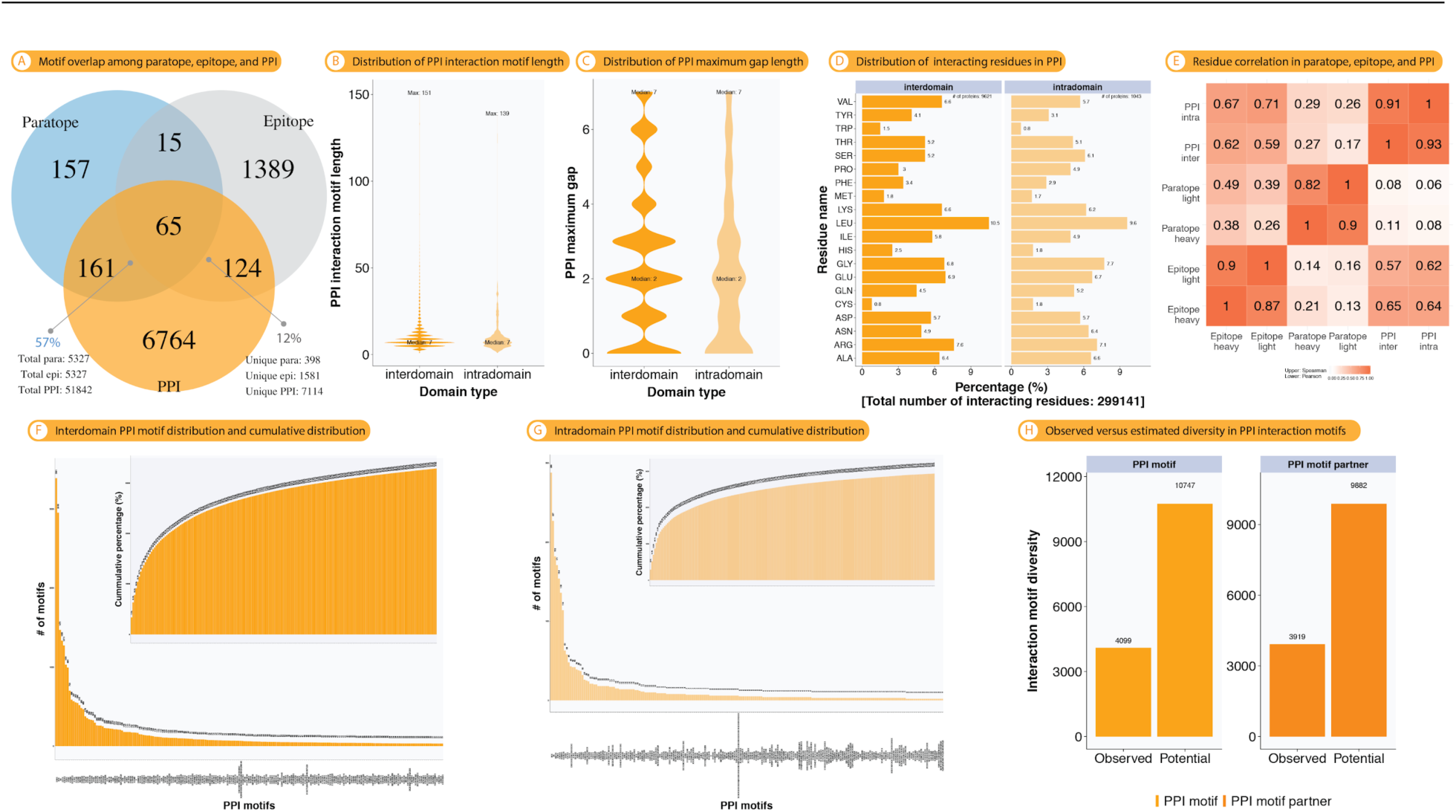
Characterization and comparison of interaction motifs found in non-immune protein-protein interaction (PPI) with those found in antibody-antigen complexes. (A) Overlap of paratope, epitope and PPI interaction motifs. Whereas paratope motifs partly overlap with motifs found in PPI (57%), most epitope motifs do not overlap with those of PPI (12%). The number of total non-unique motifs is also displayed. (B) Distribution of PPI interaction motif lengths. (C) Distribution of maximum gap lengths of PPI interaction motifs. (D) Amino acid distribution of interacting residues in PPI. (E) Pearson (lower triangular matrix) and Spearman (upper triangular matrix) correlation of the amino acid usage in paratope (heavy, light chain), epitope (heavy, light chain) and protein-protein interaction (intradomain and interdomain). (F) Inter- and (G) intradomain (cumulative) frequency distribution of the top 200 inter- and intradomain PPI motifs. Motif frequency denotes the number of times a motif has been detected within (intra) or across (inter) structures (1,043 and 9,621, respectively). (H) Estimation of the potential motif diversity using the Chao1 estimator (see *Methods*).

**Supplemental Figure S7.**
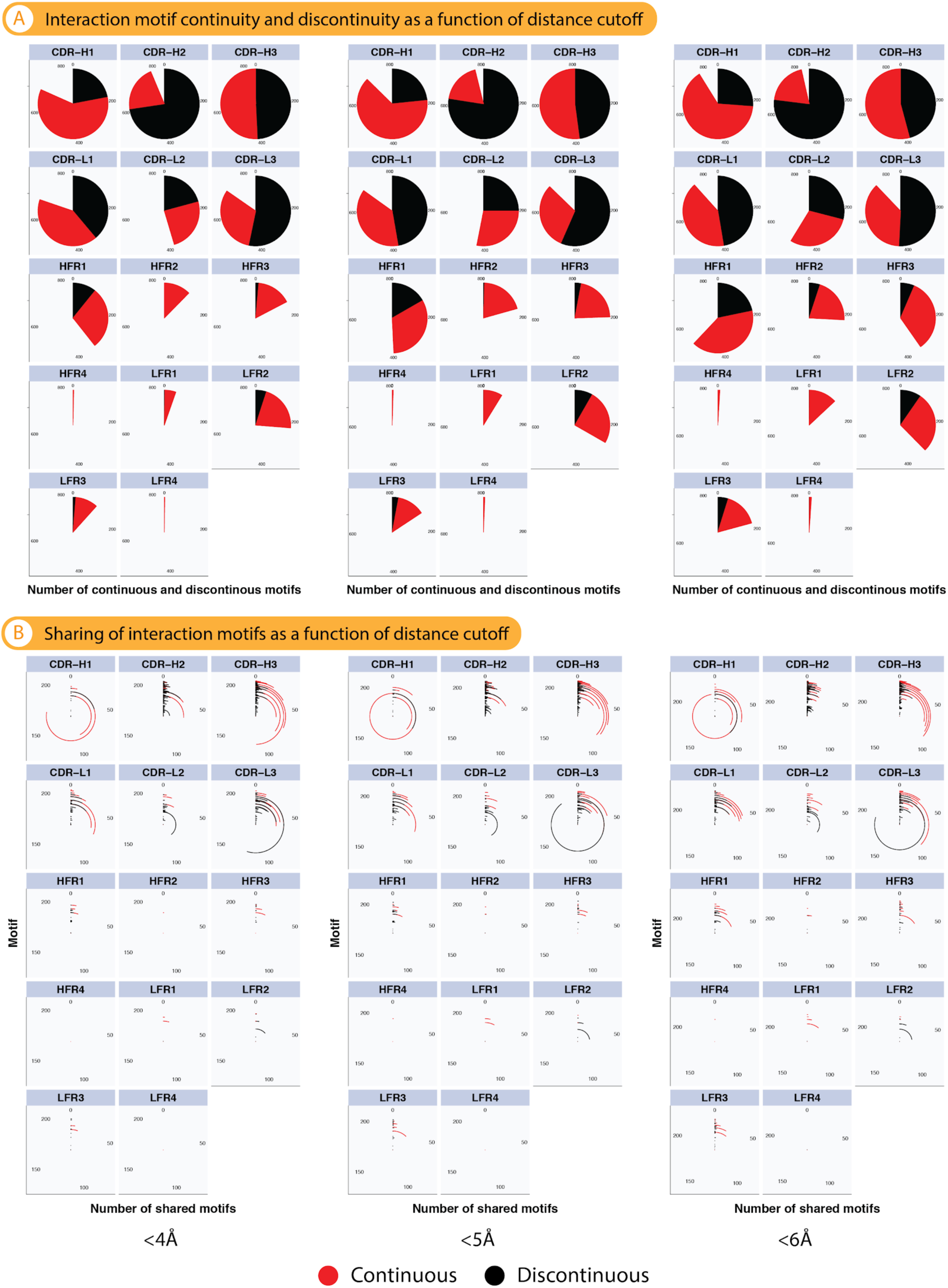
The extent of motif continuity and sharing across antibody-antigen structures is robust to the distance cutoff chosen. (A) Relative motif continuity and discontinuity are shown as a function of the distance cutoff (<4,5,6 Å). Ratios of continuous to discontinuous motifs are robust across distance cutoffs. **(B)** The extent of motif sharing across structures as a function of the distance cutoff is shown. The sharing profiles are robust across distance cutoffs.

**Supplemental Figure S8.**
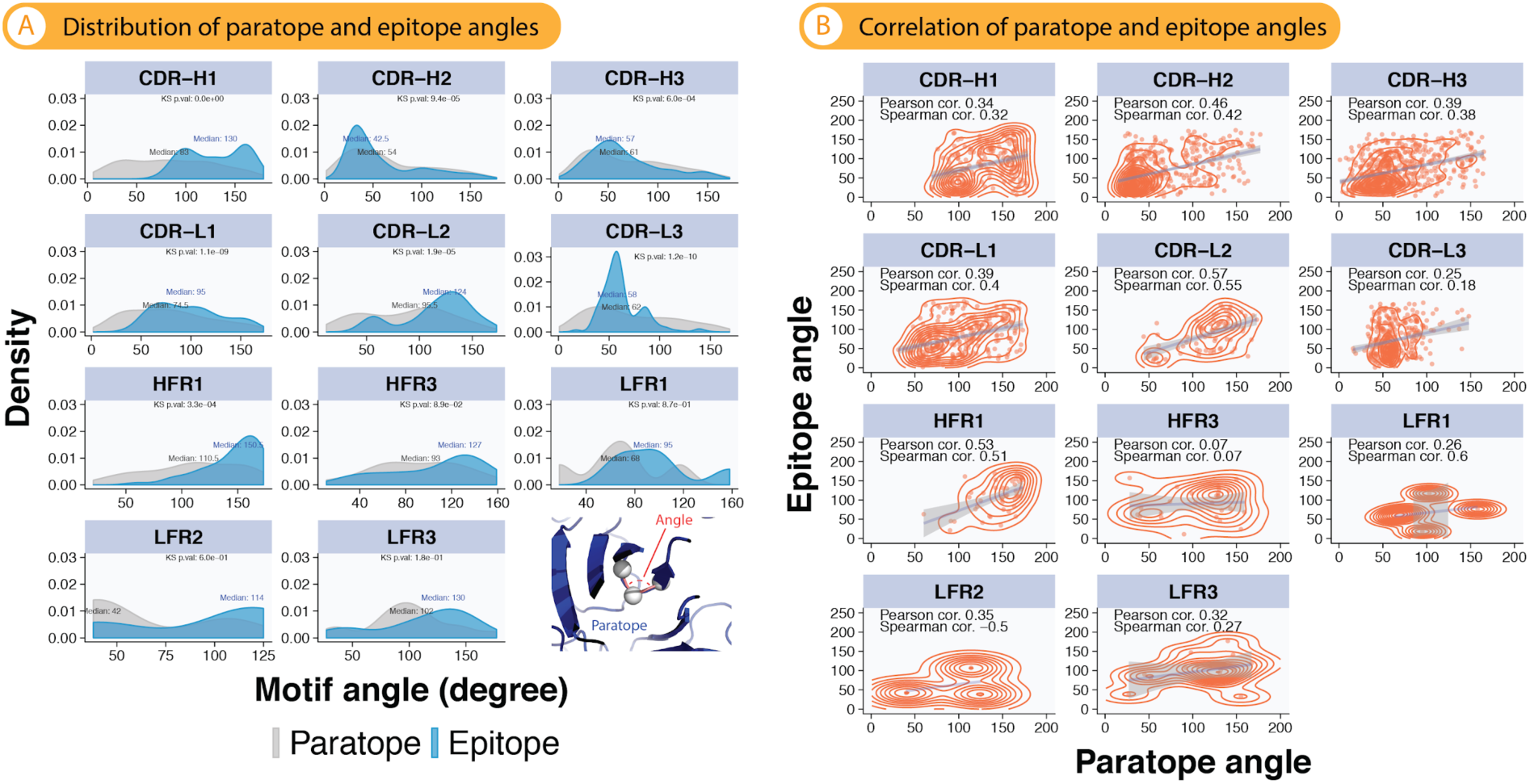
Angles of structural interaction motifs showcase diverse conformations in antibody-antigen interaction. (A) Motif angles in paratopes and epitopes (see Methods). For CDR-H2_VH,VL_ and CDR-H3_VH,VL_,the median angles in paratopes and epitopes are smaller than CDR-H1_VH,VL_ indicating that the former are looped and the latter are extended. Median motif angles in heavy and light chain of CDR3 did not substantially differ between paratopes (VH 57° and VL 58°) and epitopes (VH 61° and VL 62°), however, the difference is statistically significant indicating that epitopes assume a more extended conformation. **(B)** Angles between paratope and epitopes correlate moderately in the majority of the regions examined.

**Supplemental Figure S9.**
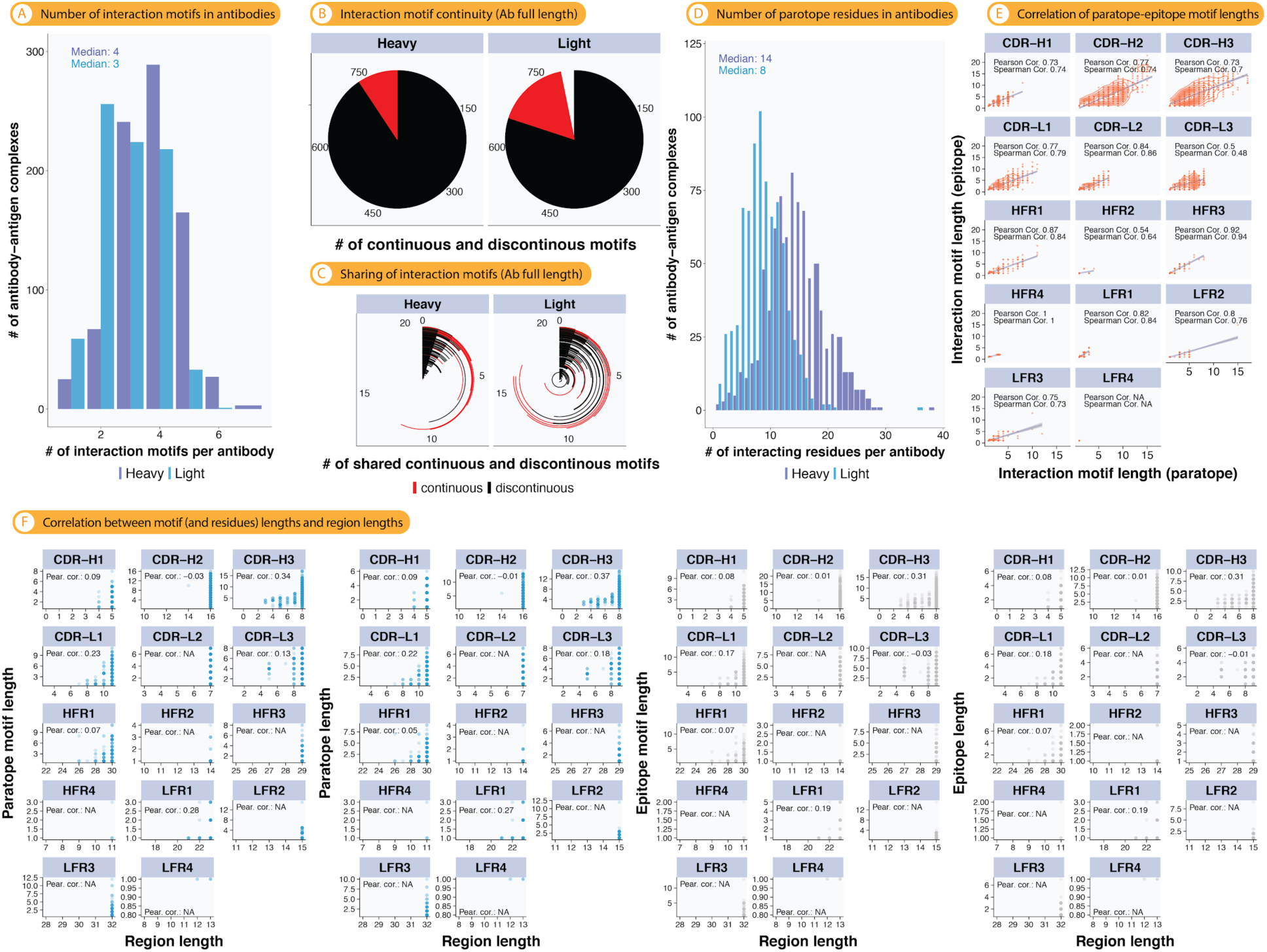
Analysis of interaction motifs on full length antibody sequences. (A) Number of paratope interaction motifs found in antibody heavy (median: 4) and light chains (median: 3). **(B)** Extent of ensembles of discontinuous (at least one discontinuous interaction motif per antibody heavy or light chain) or continuous (all motifs per antibody heavy or light chain are continuous) interaction motifs. **(C)** Sharing of discontinuous and continuous ensembles of interaction motifs across structures (heavy-chain maximum: 12 structures, light-chain maximum: 17 structures). Length distribution of paratope and epitope motifs. The length of a motif is defined as the number of characters in a motif. For example, motifs X1X and X2X are both of length 3. **(C)** Pearson (range: 0.5–1.00) and Spearman (range: 0.48–1.00) correlation of paratope and epitope lengths. **(D)** More interacting residues are found in antibody heavy chain (median: 14) in comparison to the light chain (median: 8). **(E)** The lengths of paratope and epitope motifs correlate positively in each CDR/FR segment. **(F)** Interacting residues and interaction motif lengths in general do not correlate.

**Supplemental Figure S10.**
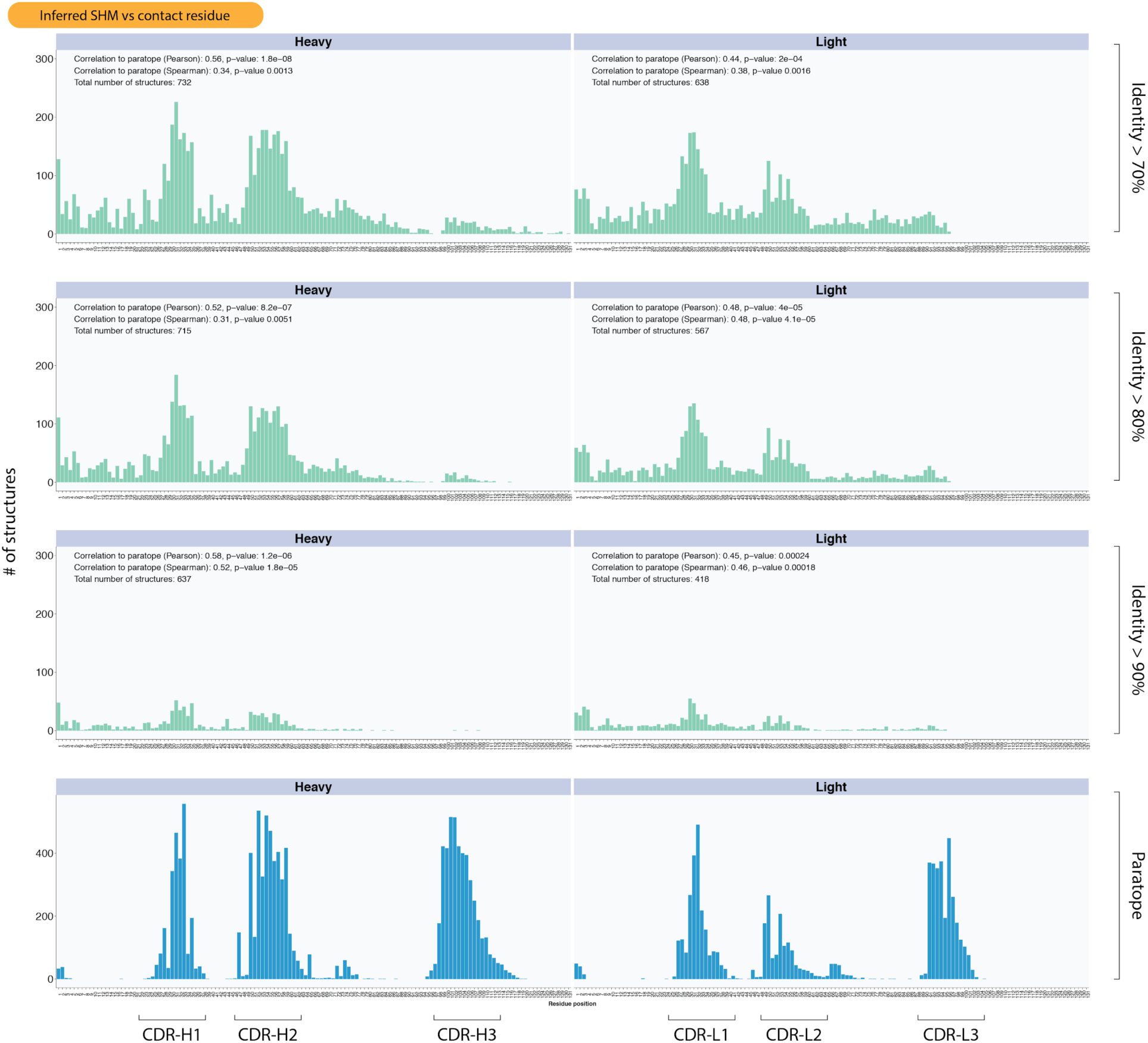
Moderate concordance between somatic hypermutation sites and contact residues. Somatic hypermutation was inferred for all human and mouse antibodies (total number of antibody-antigen complexes: 732, see Suppl Fig. S11, see Methods for details on SHM quantification). Green graphs: for each position in the heavy and light chain, the number of structures mutated in that position was quantified. Positional mutability is shown as a function of identity to germline (>70%, >80%, >90%) where identity is calculated as: (length of the sequence – the total number of SHMs) / the length of the sequence. The higher the identity, the more trustworthy the SHM count is. Blue graphs: number of structures with antigen-interacting residues in the given residue position. In each green-colored SHM graph, the correlation with the number of paratope contact sites per position is shown. Spearman (Pearson) correlation ranged between 0.31 (0.52) and 0.52 (0.58) for V_H_ and 0.38 (0.44) – 0.48 (0.48) for V_L_, indicating moderate concordance between mutated and antigen-interacting residues (all correlation values were significant, p<0.05).

**Supplemental Figure S11.**
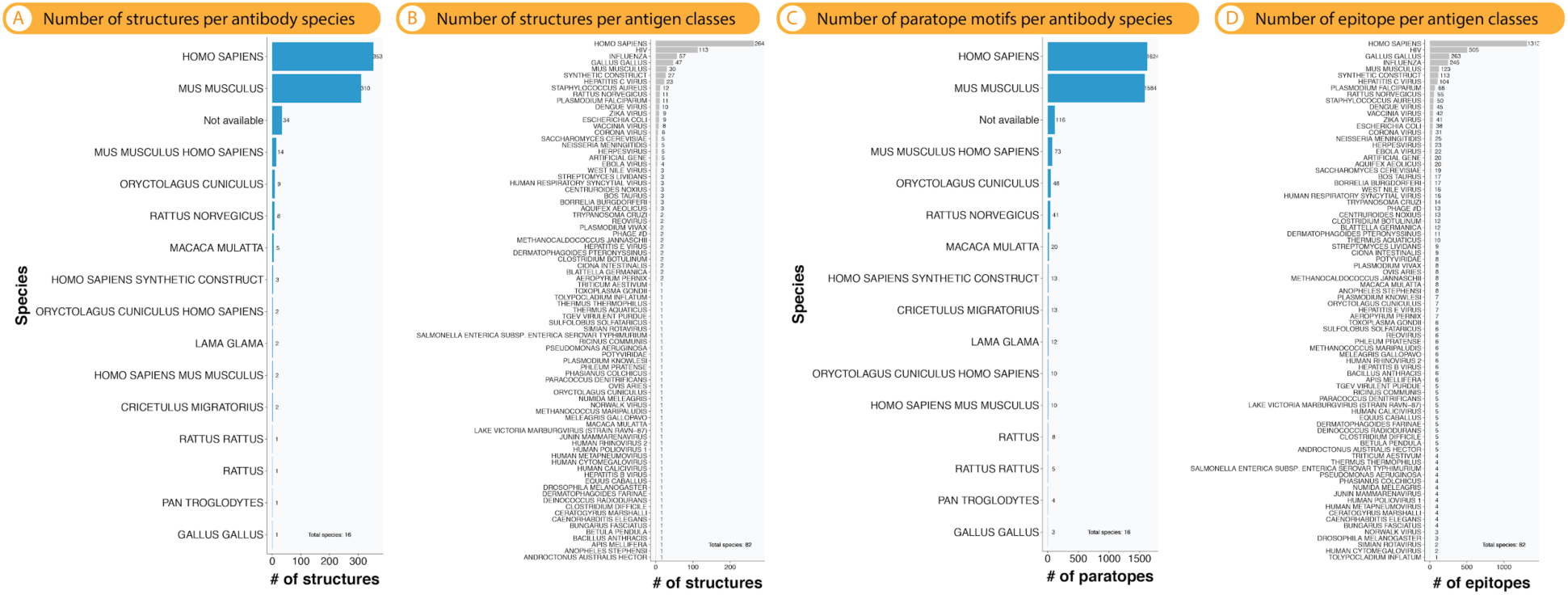
Abdb statistics: distribution of structures across species and antigen classes. (A,C) Distribution of structures (antibodies) and paratopes across species. The classes “human” and “mouse” make up together ≈90% of structures here studied. **(B,D)** Distribution of structures (antigens) and epitopes across antigen classes/species.

**Supplemental Figure S12.**
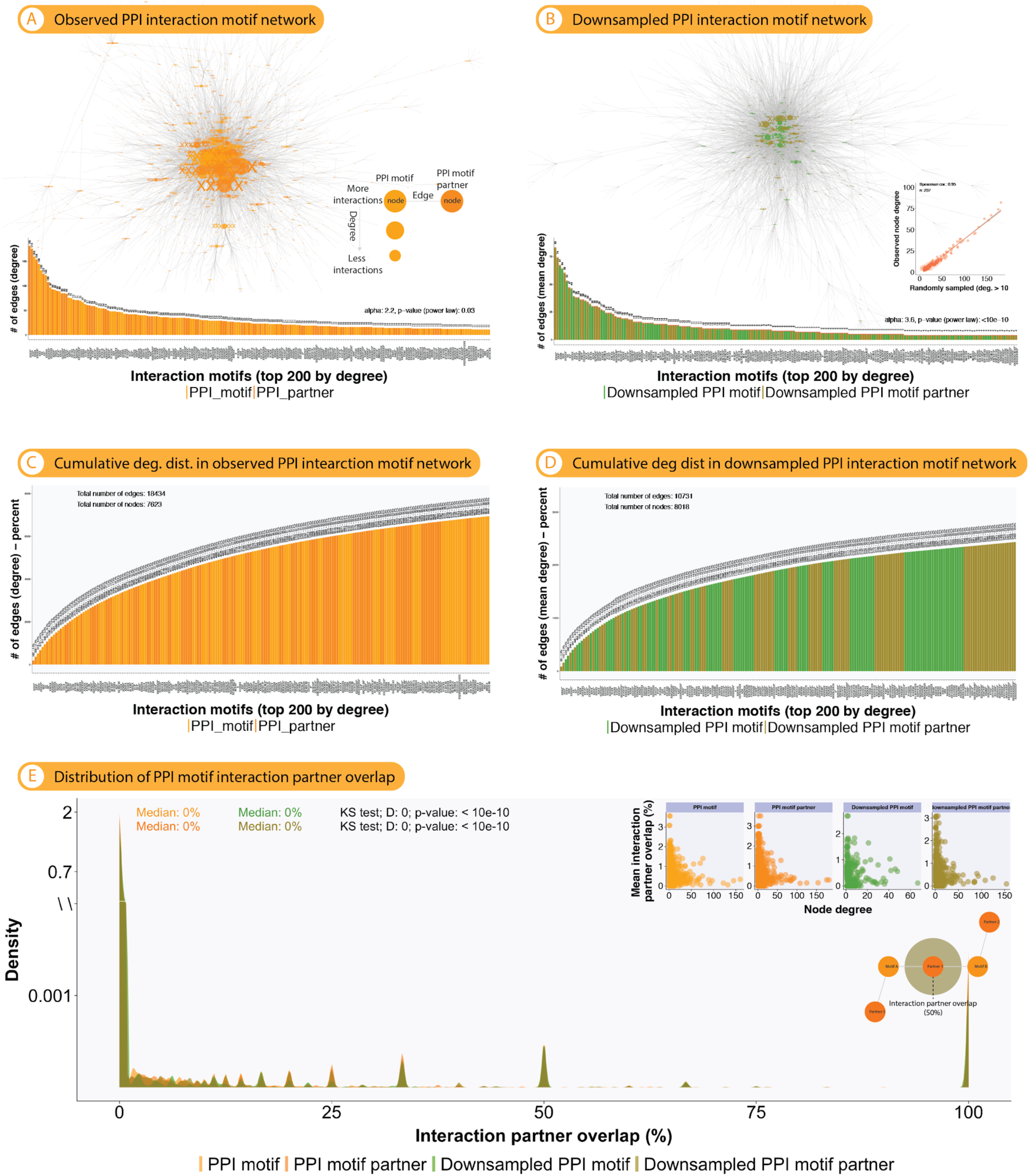
Protein-protein reactivity networks show exponential network architecture. (not a power-law distribution as observed in antibody-antigen reactivity networks [Fig. 4A], but also no Poisson-distributed networks [Fig. 4B]). The figure replicates (with a few modifications explained below) Fig. 4. Only the largest connected portion of the network was visualized, for full network see Suppl Fig. 16. **(A)** A bipartite network capturing protein-protein binding from interdomain protein-protein protein interaction (Suppl Fig. S6) was constructed. The PPI interdomain reactivity network was constructed by connecting each protein motif to its corresponding partner protein motif (total number of edges: 18,434; total number of nodes: 7,623). We here used exclusively interdomain PPI (as opposed to intradomain PPI) as interdomain PPI is most similar to antibody-antigen interaction (Suppl Fig. S6E). Network vertices were scaled by their number of connections (edges). **(B)** To more directly compare PPI reactivity network architecture to antibody-antigen interaction (Fig. 4A), we randomly sampled 100 times 5000 edges (similar to the number of edges found in the paratope-epitope reactivity network, Fig. 4A) from the reactivity network in (A). Inset: The Spearman correlation of node degree correlation of observed and randomly sampled networks is shown. For both networks, the respective node degree distribution is shown (for B, the standard error of the mean is also shown). A node’s degree is the count of its connections incoming from other nodes. **(C, D)** Cumulative degree distributions of networks (A) and (B). **(E)** Distribution of interaction partner overlap for networks (A,B). Briefly, for example for all protein binding partners in (A), the pairwise overlap of PPI motifs was calculated. The statistical significance of the difference between overlap distributions from (A) and (B) was computed using the Kolmogorov-Smirnov (KS) test. Inset: the correlation of node degree and interaction partner overlap was determined.

**Supplemental Figure S13.**
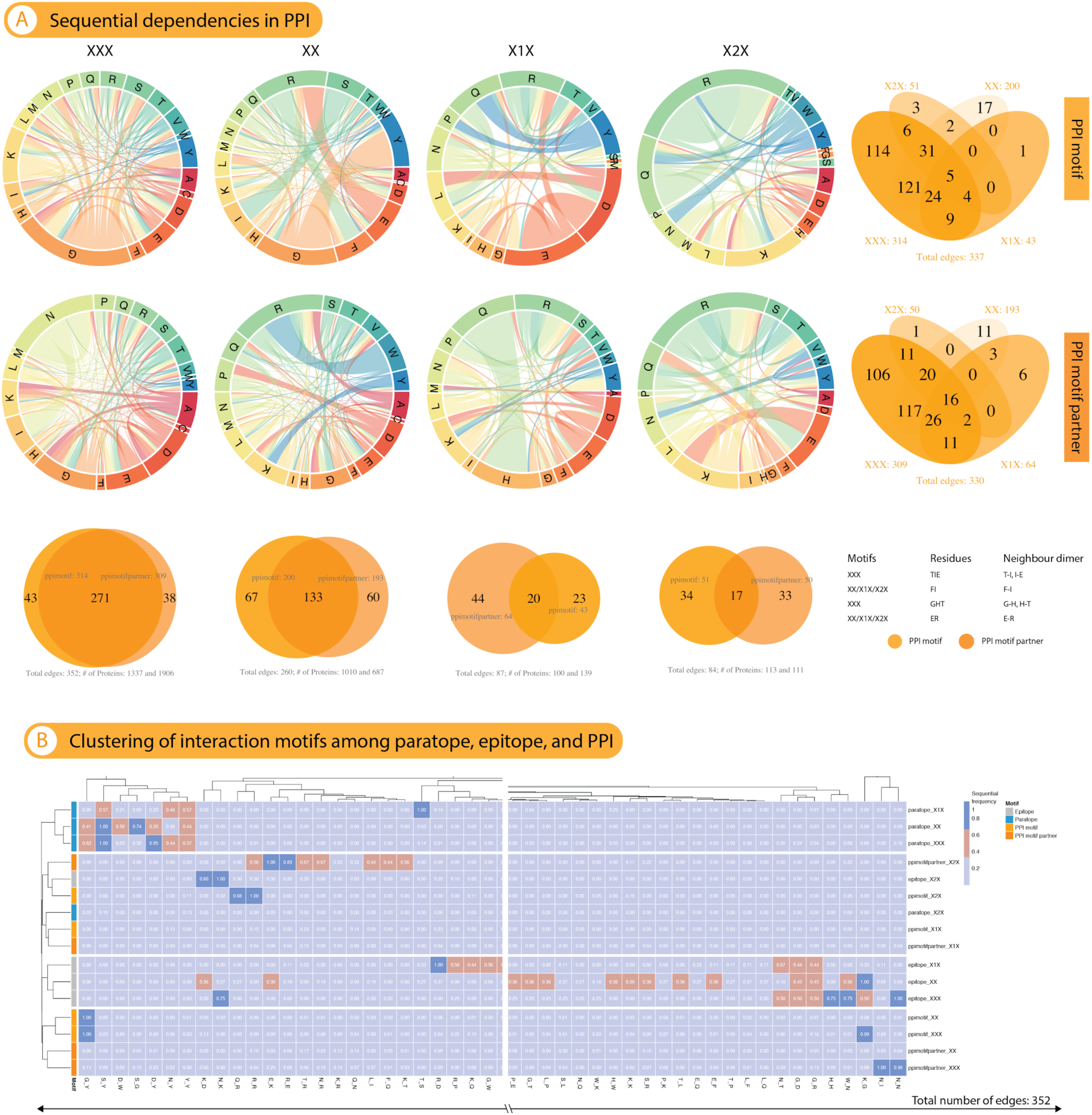
Sequential dependencies in interaction motifs of protein-protein interaction. (A) For each of the four most highly shared (across antibody-antigen structures) motifs (Suppl Fig. 4), the sequential dependency signature of PPI motifs was determined. Briefly, for the ensemble of paratope/epitope sequences mapping to a given interaction motif, the 2-mer decomposition of each paratope/epitope sequence was determined by a sliding window. For each motif, these sequential dependencies were visualized as Chord diagrams where the 20 amino acids form the segments in a track (the outermost ring) and the links indicate the frequency with which a 2-mer sequential dependency occurred (sequential dependency). In addition, Venn diagrams show the overlap of sequential dependencies (2-mers, network edges) between motifs (vertically arranged Venn diagrams) or binding partners (horizontal Venn diagrams). **(B)** Hierarchical clustering of sequential dependencies (the links are shown in (A) and Fig. 2E).

**Supplemental Figure S14.**
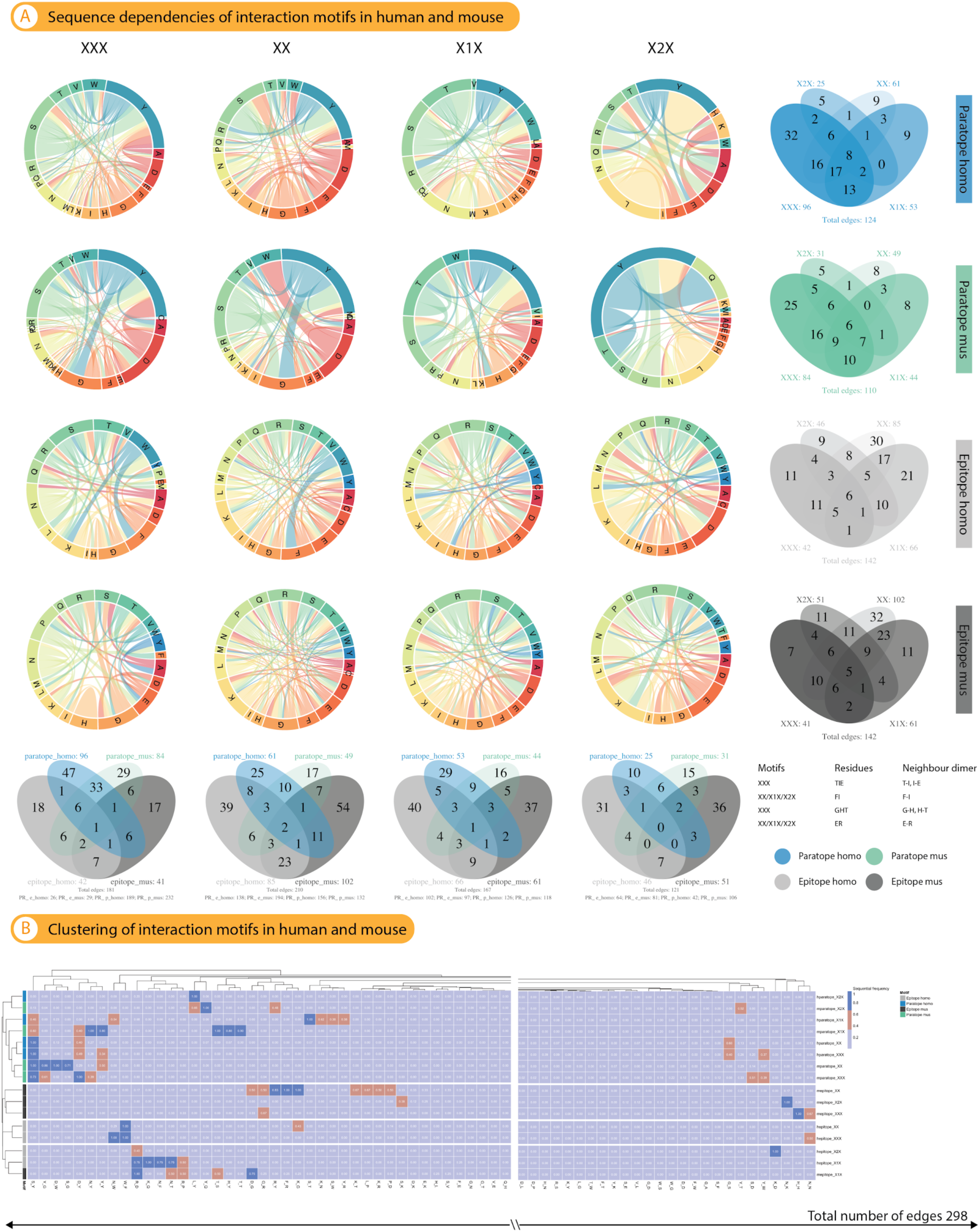
Sequence dependencies of interaction motifs in humans and mice are similar. (A) Sequence dependencies of mouse and human paratope and epitope interaction motifs were compared for the 4 most highly shared interaction motifs. Briefly, for the ensemble of paratope/epitope sequences mapping to a given interaction motif, the 2-mer decomposition of each paratope/epitope sequence was determined by sliding window. For each motif, these sequential dependencies were visualized as Chord diagrams where the 20 amino acids form the segments in a track (the outermost ring) and the links indicate the frequency with which a 2-mer sequential dependency occurred (sequential dependency). Overlap of two-mer combinations is shown by Venn diagram. **(B)** Hierarchical clustering of mouse and human paratope and epitope sequential dependencies (the links are shown in (A)).

**Supplemental Figure S15.**
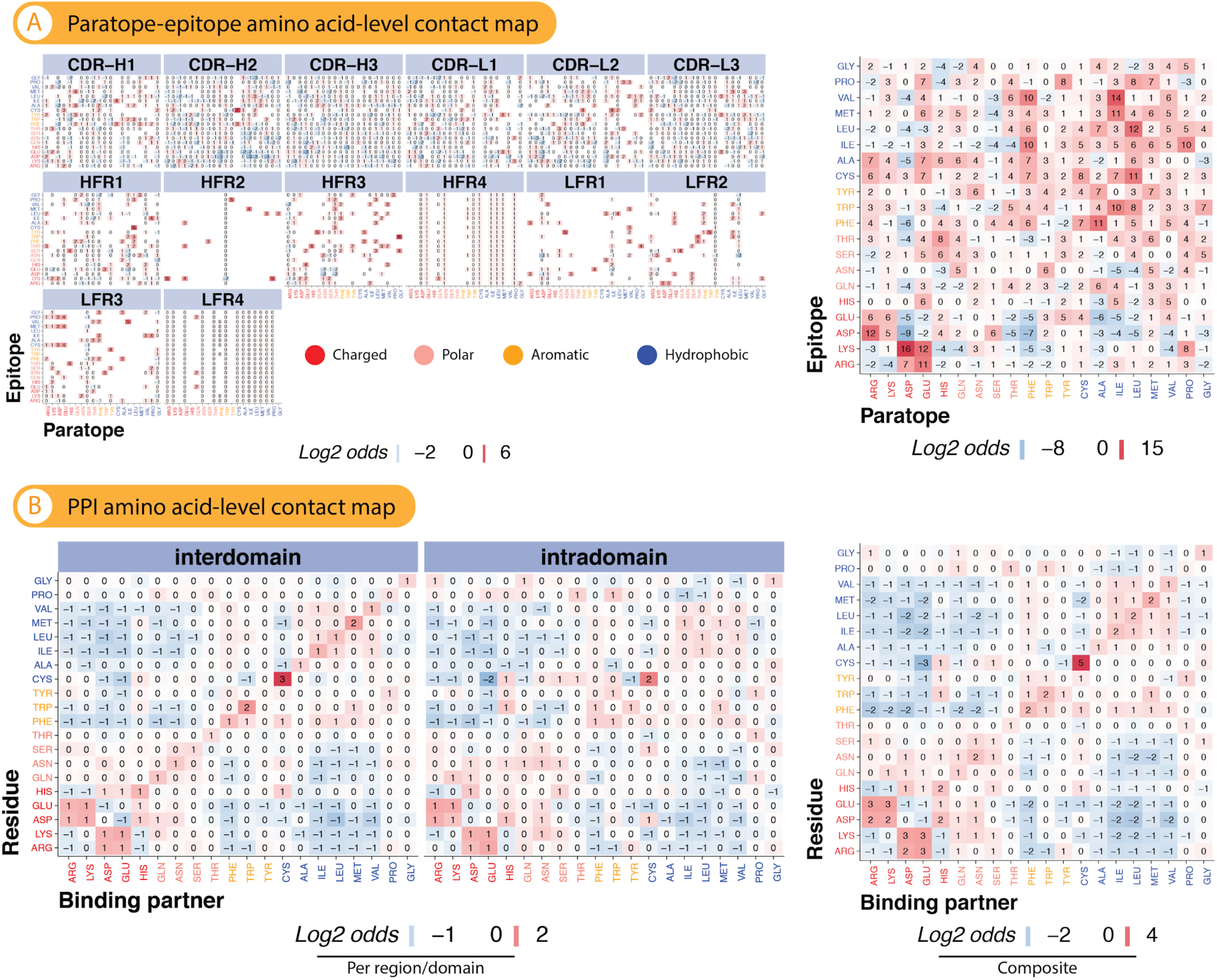
Paratope-epitope amino acid contact maps. (A) The y-axis denotes epitope residues and the x-axis denotes the paratope residues. Left: Each panel represents one CDR/FR. Right: Composite map for V_H_ and V_L_ combined. The cells are color-coded according to the preference of the amino acid pairs to interact. A red square indicates that the interaction is favored, and a blue square indicates that it occurs less frequently than expected at random (log2 odds ratio, see Methods). Amino acids are color-coded by property: charged, polar, aromatic, hydrophobic (nonpolar). **(B)** Identical to (A) except for non-immune protein-protein interaction (PPI). While there exist clear trends for PPI, namely off-diagonal amino acid interaction being disfavored and on-diagonal amino acid interaction being favored, paratope-epitope amino acid contact preferences (CDR/FR) cover a broader range in the interaction space as few amino acid interactions seem to be strictly disfavored (composite).

**Supplemental Figure S16.**
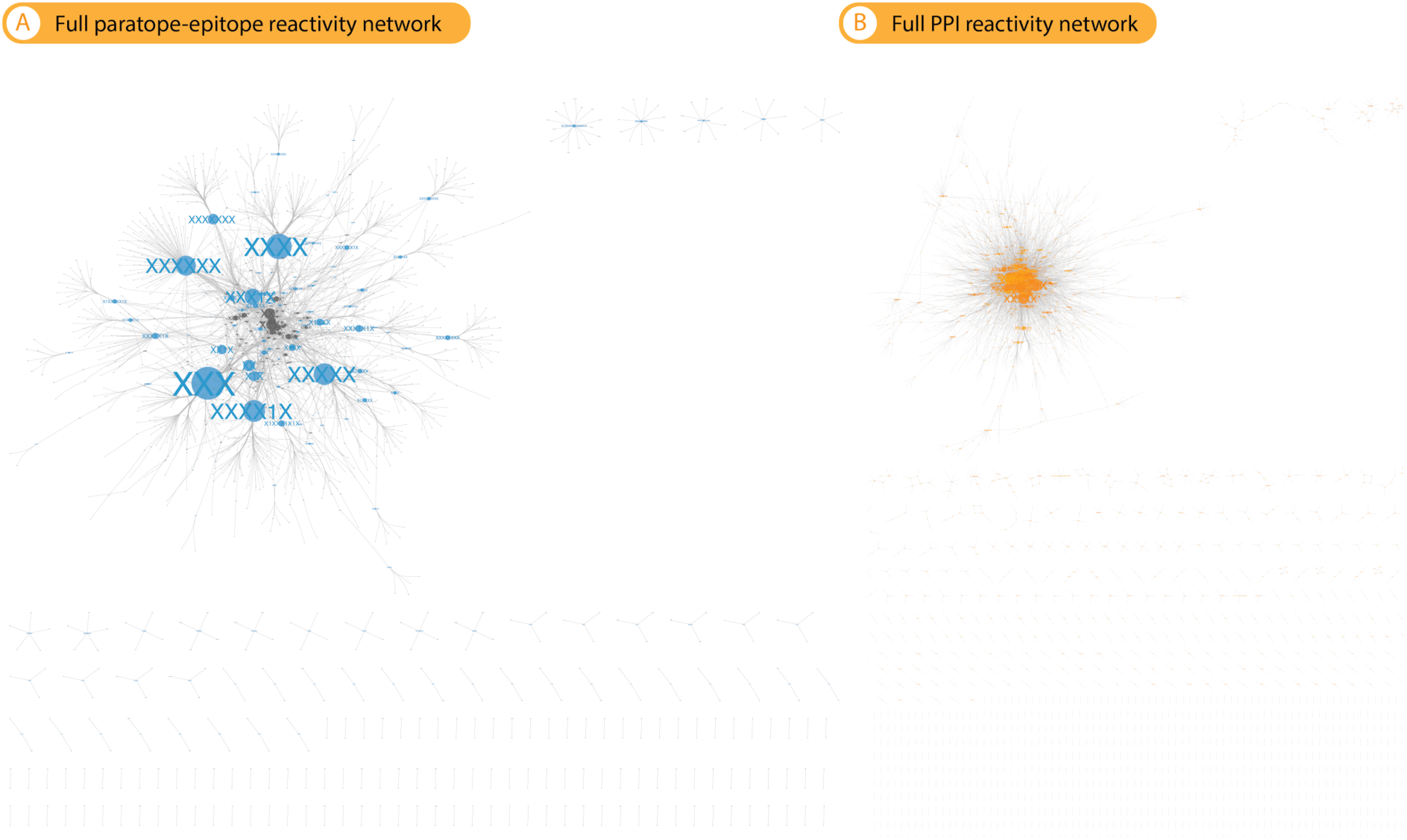
Full reactivity networks of paratope-epitope and protein-protein interaction. (A) Full paratope-epitope reactivity network which includes nodes and edges that are detached from the main network. Here shown the total number of edges 4,976 and nodes 1,981. Please refer to Fig. 4A (main network) for a methods description. **(B)** Protein-protein reactivity network (interdomain), similar to (A), the network includes nodes and edges that are detached from the main network. Here shown the total number of edges 18,434 and nodes 7,623. Please refer to Suppl Fig. S12A (main network) for a methods description.

**Supplemental Figure S17.**
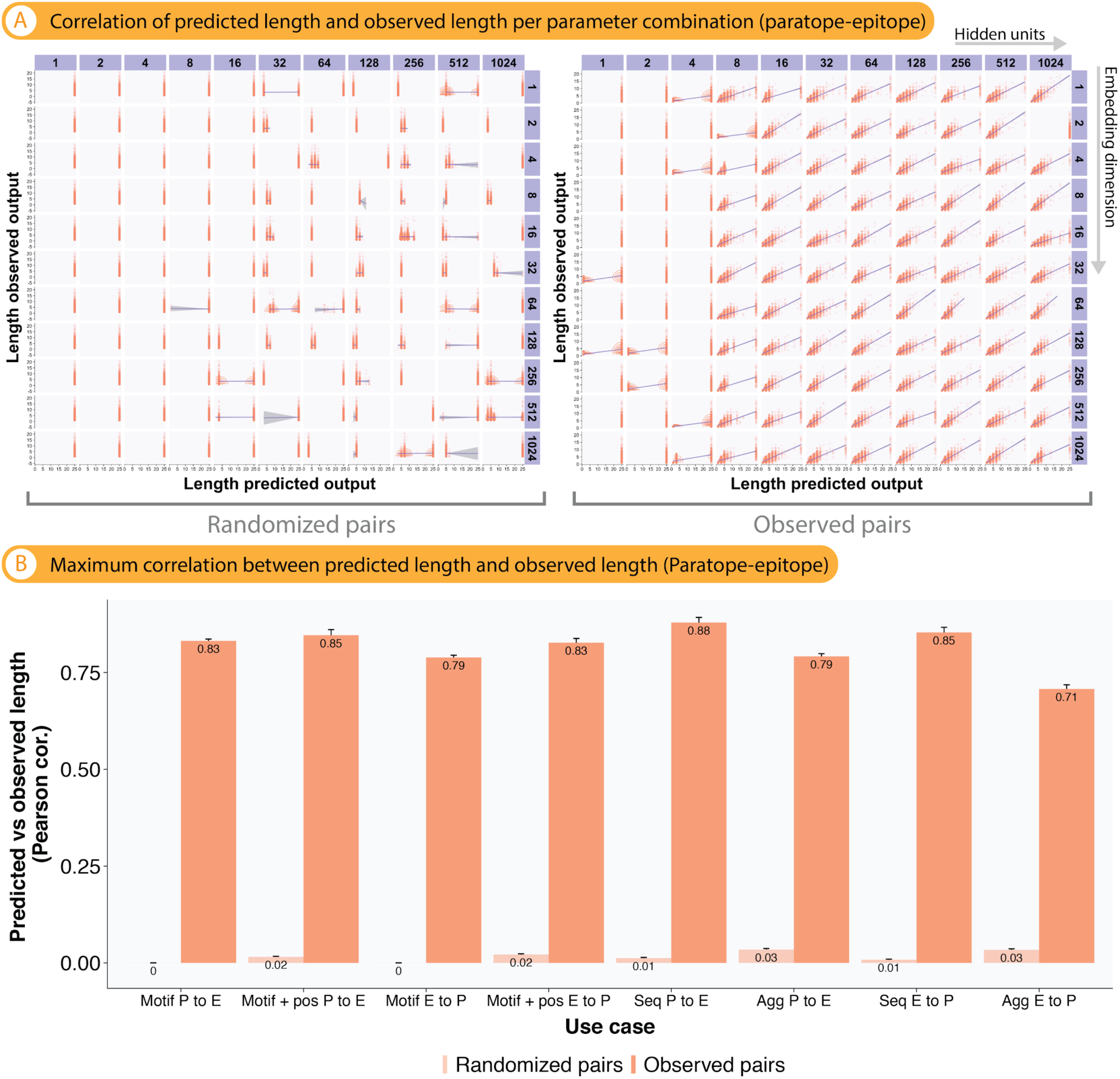
The lengths of interaction motifs and paratope and epitope sequences are learnable. Here shown for paratope-epitope data, PPI data behave similarly (not shown). Above plots is applicable only to our deep learning model as the model generates outputs character by character sequentially whereas the shallow model selects the most probable output based on the training data. **(A)** Example plot for the correlation between the predicted and observed length from one of the ten replicates in use case Sequence P→E as a function of embedding dimension and hidden units. Here shown are all pairwise combinations of the parameters (a total of 121) in each replicate. optimization Increasingly positive Pearson correlation values were recorded as the models approached optimal parameter pairs. **(B)**. Maximum predicted-observed length correlation. For the 10 replicates in each use case, the maximum correlation was obtained (the cell in (A) with the highest Pearson correlation) resulting in a total of 10 maximum correlation values. The mean of the maximum correlation values from each use case was computed and visualized as bar chart. The standard error of the mean of maximum correlation) was visualized as error bars. Use cases cover the bidirectional prediction tasks (paratope to epitope as well as epitope to paratope) of motif to motif, motif with position to motif with position, and finally amino acid sequence to amino acid sequence (see Table 1). Baseline prediction accuracies (control) were calculated based on label-shuffled data where antibody-antigen-binding partners were randomly shuffled (randomized pairs).

**Supplemental Figure S18.**
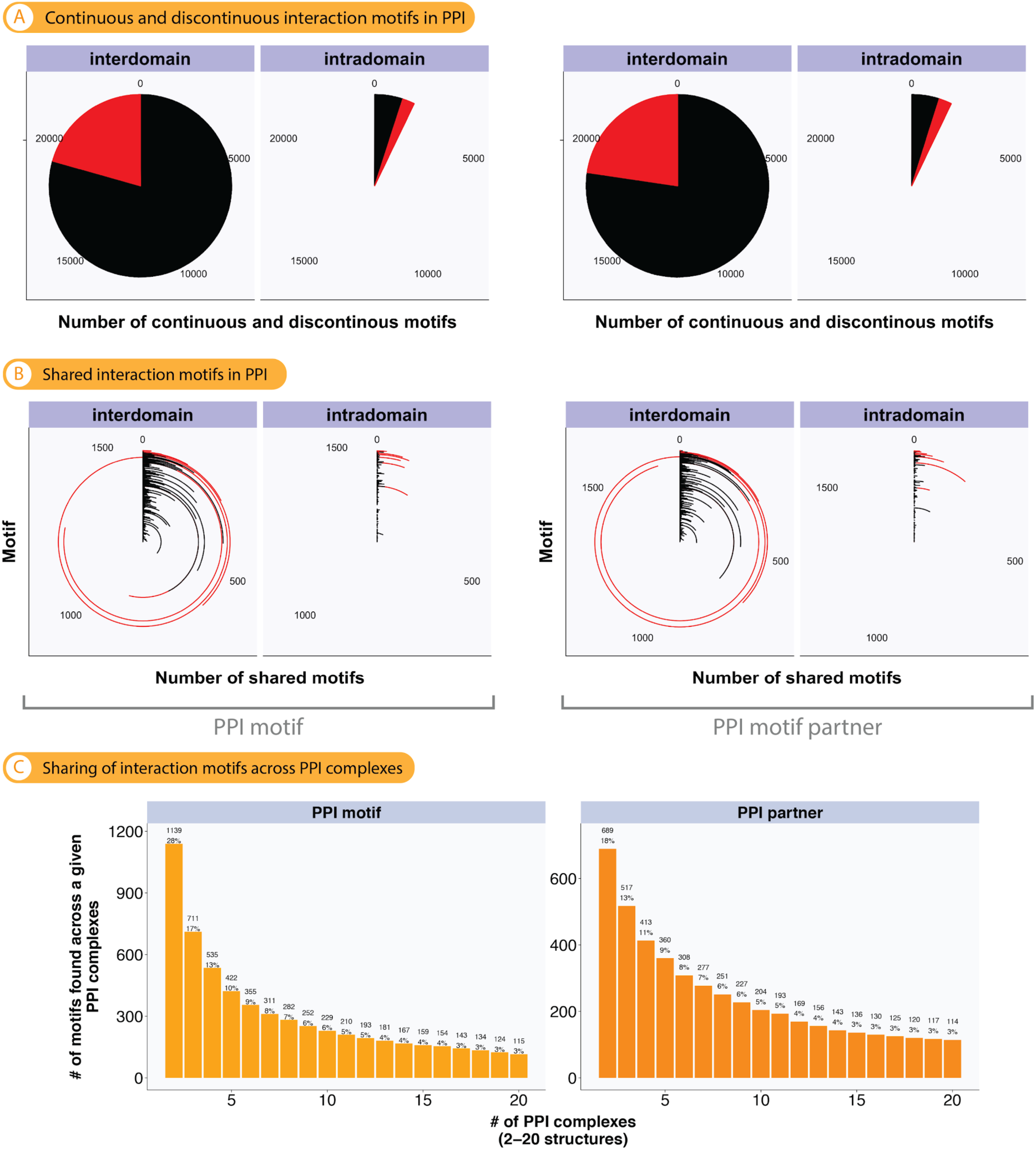
PPI interaction motifs tend to be discontinuous. Continuous motifs, however, tend to be shared. (A) Extent of discontinuous (at least one non-interacting residue) or continuous PPI interaction motifs across PPI complexes (see Fig. 3 for more methodological details). **(B)** Extent of sharing of discontinuous and continuous PPI interaction motifs across structures. (**C)** Absolute number and percentage of PPI structures a given interaction motif is found in.

**Supplemental Figure S19.**
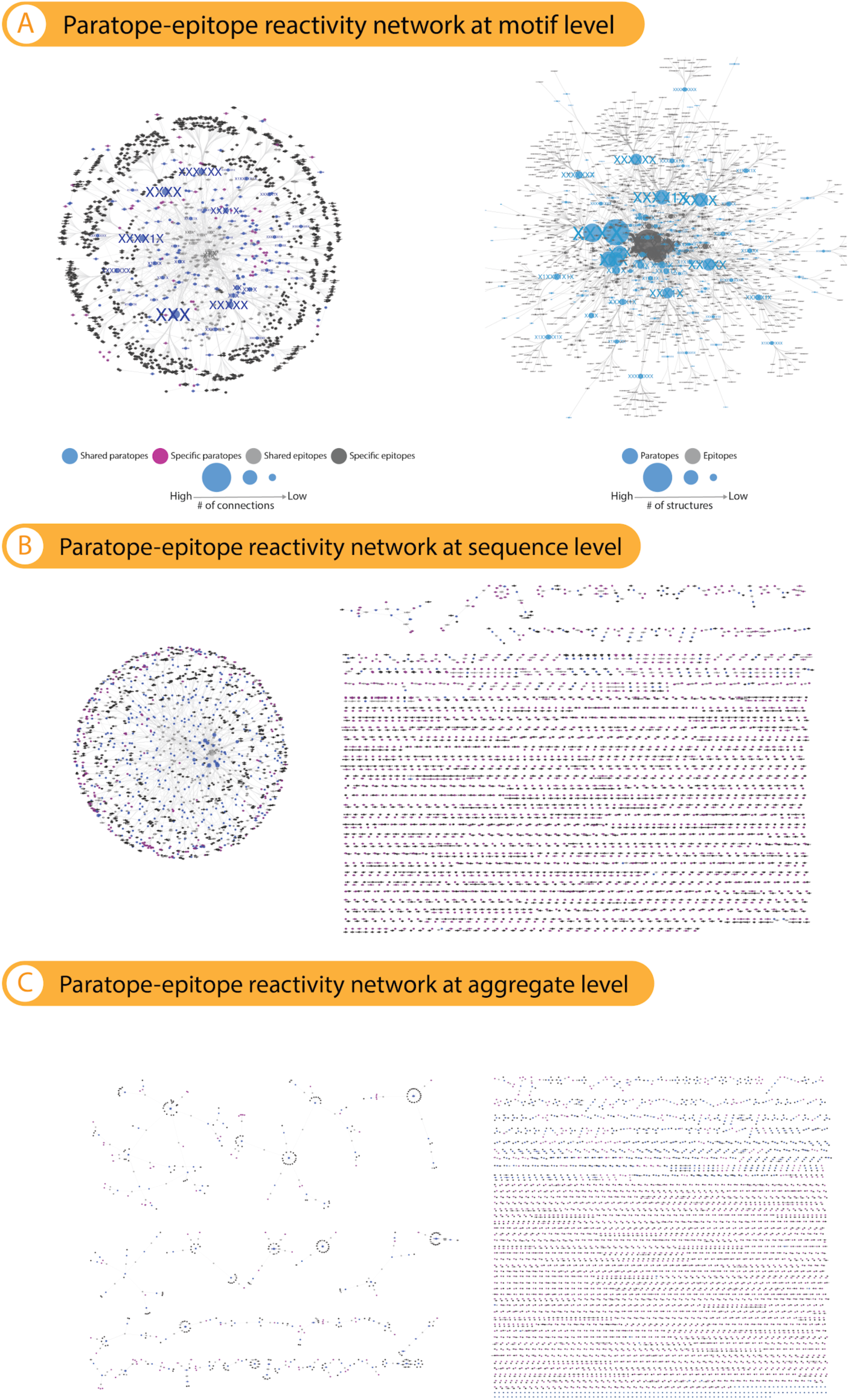
Reactivity networks according to different encodings. (A) Left: Bipartite reactivity network capturing paratope-epitope interaction constructed by connecting each paratope motif to its corresponding epitope motif (undirected edge), as in Figure 4A, including all clusters and visualizing the nodes according to their edge-betweenness (see Methods). This allows us to visually separate nodes by degree. Paratopes and epitope nodes are color-coded by specificity (specific: one exclusive binding partner, shared: >1 binding partners). Right: Reactivity network as described in (A) and Fig. 4A except that the nodes are scaled by the number of underlying structures instead of the number of connections. We performed this analysis to exclude the possibility that the motif frequency distribution has an effect on the reactivity network. **(B)** Bipartite reactivity network capturing paratope-epitope interaction at the sequence level: the sequence of binding amino acids inside motifs on the epitope or paratope are used as nodes. When the reactivity network is drawn on the sequence level, the hierarchy of nodes with paratope branches is lost. **(C)** Reactivity network drawn for an encoding where motif and sequence are merged (called “aggregate” in Figure 5). For example, the paratope RA2VG (R,A,V,G are interacting residues, and 2 residues are non-interacting) is encoded as RA--VG. Here, the network’s structure is pinned down to more independent clusters. All graphs represent the structure of datasets used to perform deep learning classifications in Figures 5 and Suppl Fig. 20.

**Supplemental Figure S20.**
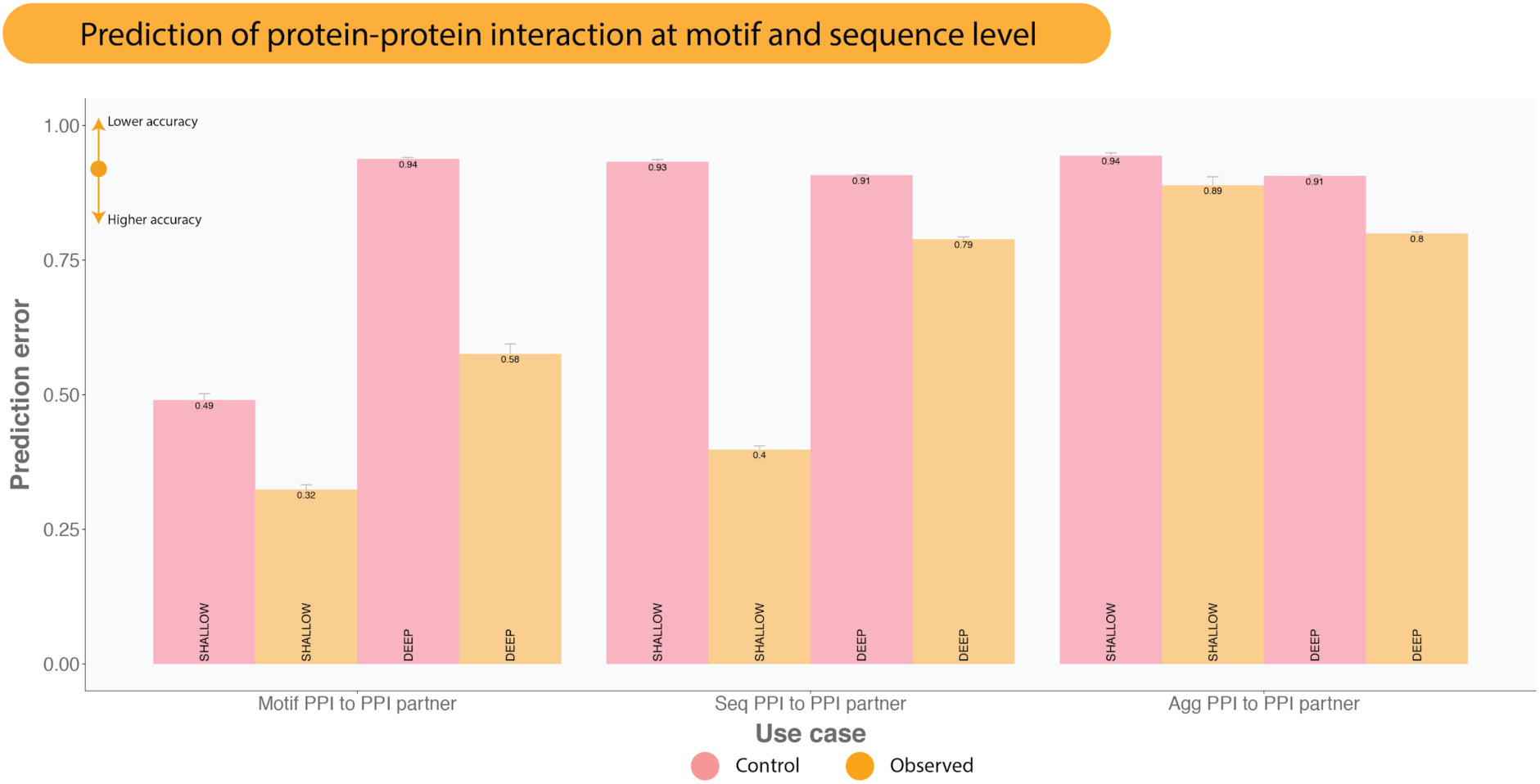
Predictability of protein-protein interaction mirrors the trends found in paratope-epitope interaction. Similar to paratope-epitope interaction, the interactions are more predictable at motif level (median errors: 32–58%; median baseline errors: 49–94%) compared to sequence (median errors: 0.4–0.79%: median baseline errors: 91–93%) and aggregate (median errors: 0.8–0.89%; median baseline errors: 91–94%) levels.

**Supplemental Figure S21.**
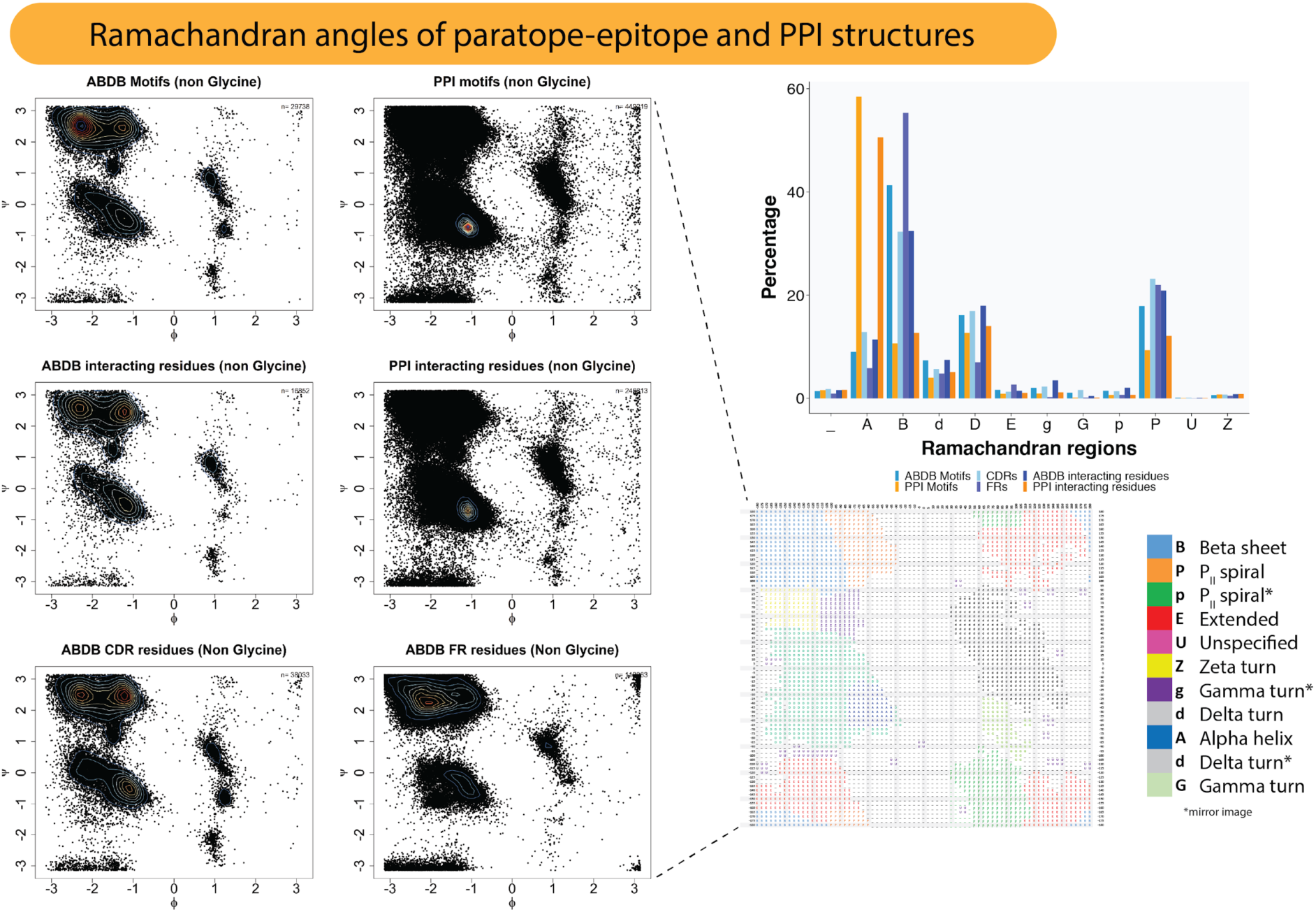
Ramachandran plot analysis: Antibodies are structurally distinct from non-immune proteins. Ramachandran angles, Phi (Φ) and Psi (Ψ), were extracted from each amino acid in motifs (both interacting residues and non-interacting residues) as compared to the set of interacting residues. These pairs of angles were used to group the residues into eight Ramachandran regions (bottom right corner) following Hollingsworth and colleagues (*77*). Here shown PPI (mostly manifests as alpha helix) and antibodies (mostly manifests as beta strand/sheet, P_II_ spiral, and delta turn) gravitates towards different angles thereby underlining the distinction between PPI and antibody-antigen interaction.

---

**Supplemental Table S1.**
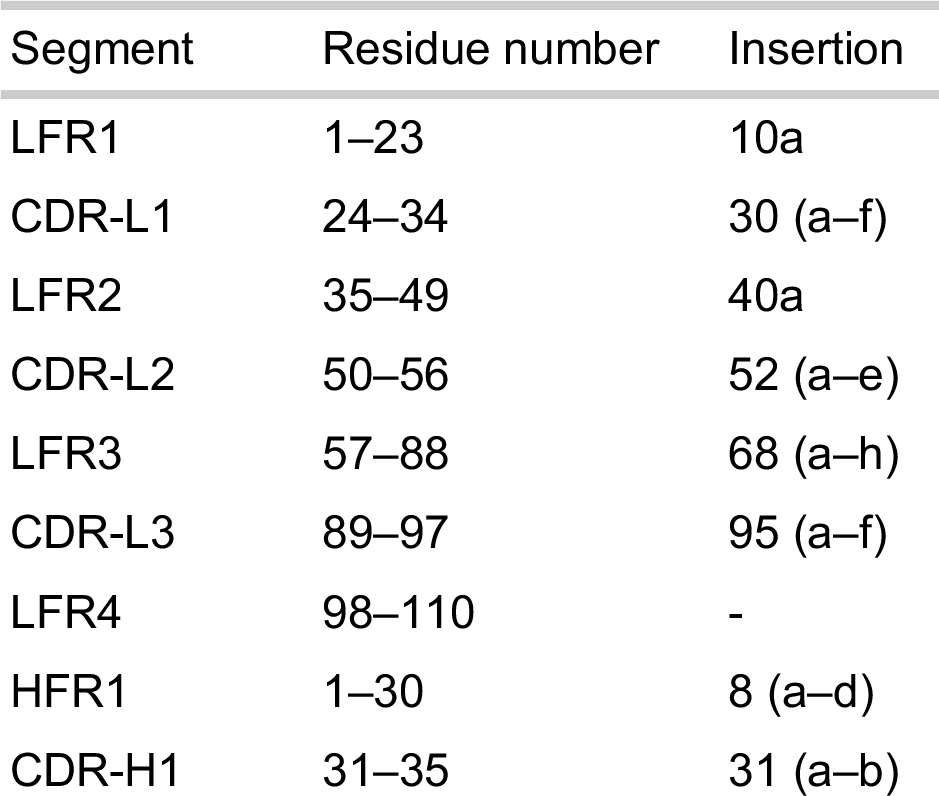
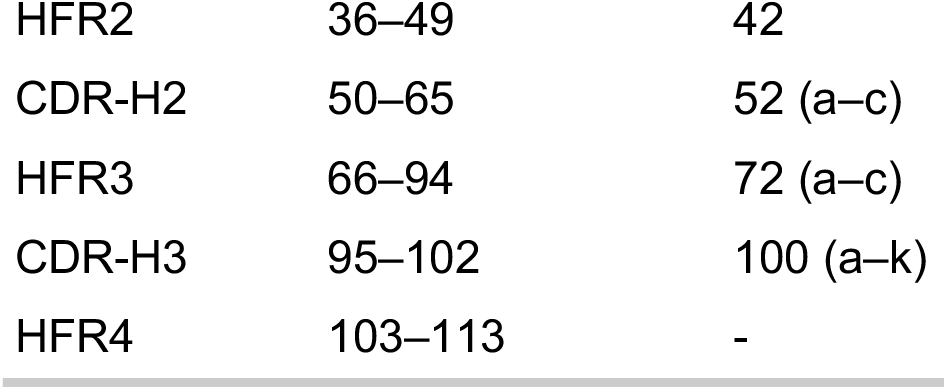
Antibody sequence numbering according to the Martin (*57*) numbering scheme.

**Supplemental Table S2.** Examples of antibody chains numbered according to the Martin (*57*) numbering scheme.

| 2EH8_1_H | 2EH8_1_L | 2LTQ_1_H | 2LTQ_1_L | 3JBA_1_H | 3JBA_1_L |
| --- | --- | --- | --- | --- | --- |
| Q-GLN-1 | D-ASP-1 | E-GLU-1 | D-ASP-1 | S-SER-7 | D-ASP-1 |
| V-VAL-2 | I-ILE-2 | V-VAL-2 | I-ILE-2 | G-GLY-8 | I-ILE-2 |
| Q-GLN-3 | Q-GLN-3 | Q-GLN-3 | V-VAL-3 | G-GLY-9 | V-VAL-3 |
| L-LEU-4 | M-MET-4 | L-LEU-4 | M-MET-4 | G-GLY-10 | M-MET-4 |
| V-VAL-5 | T-THR-5 | V-VAL-5 | S-SER-5 | L-LEU-11 | S-SER-5 |
| Q-GLN-6 | Q-GLN-6 | E-GLU-6 | Q-GLN-6 | V-VAL-12 | Q-GLN-6 |
| S-SER-7 | T-THR-7 | S-SER-7 | S-SER-7 | K-LYS-13 | S-SER-7 |
| G-GLY-8 | P-PRO-8 | G-GLY-8 | P-PRO-8 | P-PRO-14 | P-PRO-8 |
| A-ALA-9 | L-LEU-9 | G-GLY-9 | S-SER-9 | G-GLY-15 | S-SER-9 |
| E-GLU-10 | S-SER-10 | G-GLY-10 | S-SER-10 | G-GLY-16 | S-SER-10 |
| V-VAL-11 | L-LEU-11 | L-LEU-11 | L-LEU-11 | S-SER-17 | L-LEU-11 |
| V-VAL-12 | S-SER-12 | V-VAL-12 | A-ALA-12 | L-LEU-18 | A-ALA-12 |
| K-LYS-13 | V-VAL-13 | K-LYS-13 | V-VAL-13 | K-LYS-19 | V-VAL-13 |
| P-PRO-14 | T-THR-14 | P-PRO-14 | S-SER-14 | L-LEU-20 | S-SER-14 |
| G-GLY-15 | P-PRO-15 | G-GLY-15 | A-ALA-15 | S-SER-21 | V-VAL-15 |
| A-ALA-16 | G-GLY-16 | G-GLY-16 | G-GLY-16 | C-CYS-22 | G-GLY-16 |
| S-SER-17 | Q-GLN-17 | S-SER-17 | E-GLU-17 | E-GLU-23 | E-GLU-17 |
| V-VAL-18 | P-PRO-18 | L-LEU-18 | K-LYS-18 | A-ALA-24 | K-LYS-18 |
| K-LYS-19 | A-ALA-19 | K-LYS-19 | V-VAL-19 | S-SER-25 | V-VAL-19 |
| V-VAL-20 | S-SER-20 | L-LEU-20 | T-THR-20 | G-GLY-26 | T-THR-20 |
| S-SER-21 | I-ILE-21 | S-SER-21 | M-MET-21 | F-PHE-27 | M-MET-21 |
| C-CYS-22 | S-SER-22 | C-CYS-22 | S-SER-22 | T-THR-28 | S-SER-22 |
| K-LYS-23 | C-CYS-23 | A-ALA-23 | C-CYS-23 | F-PHE-29 | C-CYS-23 |
| A-ALA-24 | K-LYS-24 | A-ALA-24 | K-LYS-24 | S-SER-30 | K-LYS-24 |
| S-SER-25 | S-SER-25 | S-SER-25 | S-SER-25 | S-SER-31 | S-SER-25 |
| G-GLY-26 | S-SER-26 | G-GLY-26 | S-SER-26 | Y-TYR-32 | S-SER-26 |
| Y-TYR-27 | Q-GLN-27 | F-PHE-27 | Q-GLN-27 | A-ALA-33 | Q-GLN-27 |
| A-ALA-28 | S-SER-28 | A-ALA-28 | S-SER-28 | M-MET-34 | S-SER-28 |
| F-PHE-29 | L-LEU-29 | F-PHE-29 | L-LEU-29 | S-SER-35 | L-LEU-29 |
| S-SER-30 | L-LEU-30 | S-SER-30 | L-LEU-30 | W-TRP-36 | L-LEU-30 |
| S-SER-31 | Y-TYR-30 | S-SER-31 | N-ASN-30 | V-VAL-37 | Y-TYR-30 |
| S-SER-32 | S-SER-30 | Y-TYR-32 | S-SER-30 | R-ARG-38 | S-SER-30 |
| W-TRP-33 | N-ASN-30 | D-ASP-33 | R-ARG-30 | Q-GLN-39 | N-ASN-30 |
| M-MET-34 | G-GLY-30 | M-MET-34 | T-THR-30 | T-THR-40 | T-THR-30 |
| N-ASN-35 | K-LYS-30 | S-SER-35 | R-ARG-30 | P-PRO-41 | Q-GLN-30 |
| W-TRP-36 | T-THR-31 | W-TRP-36 | K-LYS-30 | E-GLU-42 | K-LYS-30 |
| V-VAL-37 | Y-TYR-32 | V-VAL-37 | N-ASN-31 | K-LYS-43 | N-ASN-31 |
| R-ARG-38 | L-LEU-33 | R-ARG-38 | Y-TYR-32 | R-ARG-44 | Y-TYR-32 |
| Q-GLN-39 | N-ASN-34 | Q-GLN-39 | L-LEU-33 | L-LEU-45 | L-LEU-33 |
| A-ALA-40 | W-TRP-35 | T-THR-40 | A-ALA-34 | E-GLU-46 | A-ALA-34 |
| P-PRO-41 | L-LEU-36 | P-PRO-41 | W-TRP-35 | W-TRP-47 | W-TRP-35 |
| G-GLY-42 | L-LEU-37 | E-GLU-42 | Y-TYR-36 | V-VAL-48 | Y-TYR-36 |
| Q-GLN-43 | Q-GLN-38 | K-LYS-43 | Q-GLN-37 | A-ALA-49 | Q-GLN-37 |
| G-GLY-44 | K-LYS-39 | R-ARG-44 | Q-GLN-38 | S-SER-50 | Q-GLN-38 |
| L-LEU-45 | P-PRO-40 | L-LEU-45 | K-LYS-39 | I-ILE-51 | K-LYS-39 |
| E-GLU-46 | G-GLY-41 | E-GLU-46 | P-PRO-40 | S-SER-52 | P-PRO-40 |
| W-TRP-47 | Q-GLN-42 | W-TRP-47 | G-GLY-41 | S-SER-53 | G-GLY-41 |
| I-ILE-48 | S-SER-43 | V-VAL-48 | Q-GLN-42 | G-GLY-54 | Q-GLN-42 |
| G-GLY-49 | P-PRO-44 | A-ALA-49 | S-SER-43 | G-GLY-55 | S-SER-43 |
| R-ARG-50 | K-LYS-45 | Y-TYR-50 | P-PRO-44 | N-ASN-56 | P-PRO-44 |
| I-ILE-51 | R-ARG-46 | I-ILE-51 | K-LYS-45 | T-THR-57 | K-LYS-45 |
| Y-TYR-52 | L-LEU-47 | S-SER-52 | L-LEU-46 | H-HIS-58 | L-LEU-46 |
| P-PRO-52 | I-ILE-48 | S-SER-52 | L-LEU-47 | Y-TYR-59 | L-LEU-47 |
| G-GLY-53 | Y-TYR-49 | G-GLY-53 | I-ILE-48 | P-PRO-60 | I-ILE-48 |
| D-ASP-54 | L-LEU-50 | G-GLY-54 | Y-TYR-49 | D-ASP-61 | Y-TYR-49 |
| G-GLY-55 | V-VAL-51 | G-GLY-55 | W-TRP-50 | S-SER-62 | W-TRP-50 |
| D-ASP-56 | S-SER-52 | S-SER-56 | A-ALA-51 | V-VAL-63 | A-ALA-51 |
| T-THR-57 | K-LYS-53 | T-THR-57 | S-SER-52 | K-LYS-64 | S-SER-52 |
| N-ASN-58 | L-LEU-54 | Y-TYR-58 | T-THR-53 | G-GLY-65 | T-THR-53 |
| Y-TYR-59 | D-ASP-55 | Y-TYR-59 | R-ARG-54 | R-ARG-66 | R-ARG-54 |
| A-ALA-60 | S-SER-56 | P-PRO-60 | E-GLU-55 | F-PHE-67 | E-GLU-55 |
| Q-GLN-61 | G-GLY-57 | D-ASP-61 | S-SER-56 | T-THR-68 | S-SER-56 |
| K-LYS-62 | V-VAL-58 | T-THR-62 | G-GLY-57 | I-ILE-69 | G-GLY-57 |
| F-PHE-63 | P-PRO-59 | V-VAL-63 | V-VAL-58 | S-SER-70 | V-VAL-58 |
| Q-GLN-64 | D-ASP-60 | K-LYS-64 | P-PRO-59 | R-ARG-71 | P-PRO-59 |
| G-GLY-65 | R-ARG-61 | G-GLY-65 | D-ASP-60 | D-ASP-72 | D-ASP-60 |
| K-LYS-66 | F-PHE-62 | R-ARG-66 | R-ARG-61 | N-ASN-72 | R-ARG-61 |
| A-ALA-67 | S-SER-63 | F-PHE-67 | F-PHE-62 | A-ALA-72 | F-PHE-62 |
| T-THR-68 | G-GLY-64 | T-THR-68 | T-THR-63 | R-ARG-72 | T-THR-63 |
| L-LEU-69 | S-SER-65 | I-ILE-69 | G-GLY-64 | N-ASN-73 | G-GLY-64 |
| T-THR-70 | G-GLY-66 | S-SER-70 | S-SER-65 | I-ILE-74 | S-SER-65 |
| A-ALA-71 | S-SER-67 | R-ARG-71 | G-GLY-66 | L-LEU-75 | G-GLY-66 |
| D-ASP-72 | G-GLY-68 | D-ASP-72 | S-SER-67 | Y-TYR-76 | S-SER-67 |
| K-LYS-72 | T-THR-69 | N-ASN-72 | G-GLY-68 | L-LEU-77 | G-GLY-68 |
| S-SER-72 | D-ASP-70 | A-ALA-72 | T-THR-69 | Q-GLN-78 | T-THR-69 |
| T-THR-72 | F-PHE-71 | K-LYS-72 | D-ASP-70 | M-MET-79 | D-ASP-70 |
| S-SER-73 | T-THR-72 | N-ASN-73 | F-PHE-71 | S-SER-80 | F-PHE-71 |
| T-THR-74 | L-LEU-73 | T-THR-74 | T-THR-72 | S-SER-81 | T-THR-72 |
| A-ALA-75 | K-LYS-74 | L-LEU-75 | L-LEU-73 | L-LEU-82 | L-LEU-73 |
| Y-TYR-76 | I-ILE-75 | Y-TYR-76 | T-THR-74 | R-ARG-83 | T-THR-74 |
| M-MET-77 | S-SER-76 | L-LEU-77 | I-ILE-75 | S-SER-84 | I-ILE-75 |
| E-GLU-78 | R-ARG-77 | Q-GLN-78 | S-SER-76 | E-GLU-85 | S-SER-76 |
| L-LEU-79 | V-VAL-78 | M-MET-79 | S-SER-77 | D-ASP-86 | S-SER-77 |
| S-SER-80 | E-GLU-79 | S-SER-80 | V-VAL-78 | T-THR-87 | V-VAL-78 |
| S-SER-81 | A-ALA-80 | S-SER-81 | Q-GLN-79 | A-ALA-88 | K-LYS-79 |
| L-LEU-82 | E-GLU-81 | L-LEU-82 | A-ALA-80 | M-MET-89 | A-ALA-80 |
| R-ARG-83 | D-ASP-82 | K-LYS-83 | E-GLU-81 | Y-TYR-90 | E-GLU-81 |
| S-SER-84 | V-VAL-83 | S-SER-84 | D-ASP-82 | Y-TYR-91 | D-ASP-82 |
| E-GLU-85 | G-GLY-84 | E-GLU-85 | L-LEU-83 | C-CYS-92 | L-LEU-83 |
| D-ASP-86 | V-VAL-85 | D-ASP-86 | A-ALA-84 | A-ALA-93 | A-ALA-84 |
| T-THR-87 | Y-TYR-86 | T-THR-87 | V-VAL-85 | R-ARG-94 | V-VAL-85 |
| A-ALA-88 | Y-TYR-87 | A-ALA-88 | Y-TYR-86 | G-GLY-95 | Y-TYR-86 |
| V-VAL-89 | C-CYS-88 | M-MET-89 | Y-TYR-87 | L-LEU-96 | Y-TYR-87 |
| Y-TYR-90 | V-VAL-89 | Y-TYR-90 | C-CYS-88 | Y-TYR-97 | C-CYS-88 |
| F-PHE-91 | Q-GLN-90 | Y-TYR-91 | K-LYS-89 | Y-TYR-98 | Q-GLN-89 |
| C-CYS-92 | G-GLY-91 | C-CYS-92 | Q-GLN-90 | G-GLY-99 | Q-GLN-90 |
| A-ALA-93 | T-THR-92 | A-ALA-93 | S-SER-91 | Y-TYR-100 | Y-TYR-91 |
| R-ARG-94 | H-HIS-93 | R-ARG-94 | Y-TYR-92 | D-ASP-100 | Y-TYR-92 |
| E-GLU-95 | F-PHE-94 | P-PRO-95 | N-ASN-93 | E-GLU-100 | S-SER-93 |
| Y-TYR-96 | P-PRO-95 | D-ASP-96 | L-LEU-94 | G-GLY-100 | Y-TYR-94 |
| D-ASP-97 | Q-GLN-96 | A-ALA-97 | Y-TYR-96 | S-SER-100 | P-PRO-95 |
| E-GLU-98 | T-THR-97 | M-MET-98 | T-THR-97 | D-ASP-100 | L-LEU-96 |
| A-ALA-101 | F-PHE-98 | D-ASP-101 | F-PHE-98 | F-PHE-100 | T-THR-97 |
| Y-TYR-102 | G-GLY-99 | Y-TYR-102 | G-GLY-99 | D-ASP-101 | F-PHE-98 |
| W-TRP-103 | G-GLY-100 | W-TRP-103 | G-GLY-100 | Y-TYR-102 | G-GLY-99 |
| G-GLY-104 | G-GLY-101 | G-GLY-104 | G-GLY-101 | W-TRP-103 | A-ALA-100 |
| Q-GLN-105 | T-THR-102 | Q-GLN-105 | T-THR-102 | G-GLY-104 | G-GLY-101 |
| G-GLY-106 | K-LYS-103 | G-GLY-106 | K-LYS-103 | Q-GLN-105 | T-THR-102 |
| T-THR-107 | V-VAL-104 | T-THR-107 | L-LEU-104 | G-GLY-106 | K-LYS-103 |
| L-LEU-108 | E-GLU-105 | S-SER-108 | E-GLU-105 | T-THR-107 | L-LEU-104 |
| V-VAL-109 | I-ILE-106 | V-VAL-109 | I-ILE-106 | T-THR-108 | E-GLU-105 |
| T-THR-110 | K-LYS-107 | T-THR-110 | K-LYS-107 | L-LEU-109 | L-LEU-106 |
| V-VAL-111 | R-ARG-108 | V-VAL-111 | A-ALA-108 | T-THR-110 | K-LYS-107 |
| S-SER-112 | T-THR-109 | S-SER-112 | D-ASP-109 | V-VAL-111 | R-ARG-108 |
| S-SER-113 | V-VAL-110 | S-SER-113 | A-ALA-110 | S-SER-112 | A-ALA-109 |

**Supplemental Table S3.** Top 10 protein domains and typical size sourced from Pfam (*72*) and 3did (*71*).

| Domain name | Pfam ID* | Typical number of amino acids** | Description |
| --- | --- | --- | --- |
| Proteasome | PF00227 | 190 | Protein degradation |
| C1-set | PF07654 | 80 | Ig-like constant |
| Hemagglutinin | PF00509 | 300 | Glycoprotein (Influenza) |
| V-set | PF07686 | 100 | Ig-like variable |
| WD40 | PF00400 | 40 | Structural motif |
| adh_short_c2 | PF13561 | 220 | Reductase |
| lectin_legB | PF00139 | 220 | Lectin proteins |
| EF-hand_7 | PF13378 | 220 | Enolase C-terminal domain-like |
| Ferritin | PF00210 | 140 | Ferritin-like domain |
| Catalase | PF00199 | 380 | Catalyzes the decomposition of hydrogen peroxide |

\*Pfam ID is a unique identifier to access an entry in the database of protein families (Pfam) (72).
\*\*The typical number of amino acids was obtained by mapping the Pfam domain to structures in Protein Data Bank (50) and sequences in UniProt (149).

